# Chemoenzymatic Synthesis of Genetically-Encoded Multivalent Liquid *N*-glycan Arrays

**DOI:** 10.1101/2022.08.05.503005

**Authors:** Chih-Lan Lin, Mirat Sojitra, Eric J. Carpenter, Ellen Susanah Hayhoe, Susmita Sarkar, Elizabeth Anne Volker, Alexei Atrazhev, Todd L. Lowary, Matthew S. Macauley, Ratmir Derda

## Abstract

A hallmark of cellular glycosylation is its chemical complexity and heterogeneity, which can be challenging to capture synthetically. Using chemoenzymatic synthesis on M13 phage, we produce a genetically-encoded liquid glycan array (LiGA) of biantennary complex type N-glycans. Ligation of azido-functionalized sialylglycosyl-asparagine derived from egg yolk to phage functionalized with 50–1000 copies of dibenzocyclooctyne produced divergent intermediate that can be trimmed by glycosidases and extended by glycosyltransferases to yield a library of phages with different *N*-glycans. Post-reaction analysis by MALDI-TOF MS provided a rigorous approach to confirm *N*-glycan structure and density, both of which were encoded in the bacteriophage DNA. The binding of this *N*-glycan LiGA by ten lectins, including CD22 or DC-SIGN expressed on live cells, uncovered an optimal structure/density combination for recognition. Injection of the LiGA into mice identified glycoconjugates with structures and avidity necessary for enrichment in specific organs. This work provides an unprecedented quantitative evaluation of the interaction of complex *N*-glycans with GBPs *in vitro* and *in vivo*.

## Introduction

The surface of every cell is coated with complex glycans, installed on lipids or as post-translational modifications on proteins, forming a glycocalyx at which cellular interactions occur^1–6^. Cell-surface glycans mediate biological processes as diverse as cell–cell adhesion, bacterial and viral infection, and immune regulation^7–9^. Interfering with these processes is a demonstrated strategy for drug action^10–13^. Understanding the recognition properties of glycan-binding proteins (GBPs) is necessary to unravel the role of glycans in cellular interactions and function^14–16^. A property of particular interest is density, which plays an essential role in glycan–GBP interactions^17–19^. Most GBPs have multiple glycan-binding sites, each specifically recognizing a glycan moiety or a fragment of a complex glycan^14, 20, 21^. The affinity between monovalent glycans and a GBP binding site is typically low^22^. Therefore, the spatial organization of multiple copies of the same glycan on the cell surface both influences the initial recognition event and binding, and generates clustered saccharide patches^5^ that enhance avidity via multivalency^20, 23^.

Previous efforts to probe the effect of glycan structure and density on GBP-recognition, have used microarrays^24, 25^ in which glycans are displayed on (usually) glass slides that, in principle, mimic their natural valency and spatial presentation.^26^ Despite their enormous utility, conventional glycan arrays have drawbacks; for example, they cannot assay interactions with intact cells. Furthermore, with the exception of BSA-conjugate arrays^18, 27^, it is challenging to systematically probe the effect of density using solid-phase arrays. The synthesis of homogeneous multivalent displays of glycans on polymers, dendrimers, liposomes, and other carriers offers a higher level of control of density^28–31^. Many such displays have been used to study cellular responses^19, 32, 33^. Unlike arrays, these multivalent displays cannot encode multiple types of glycans and, as a result, can probe only limited types of glycans. To bridge these technologies, we recently developed liquid glycan arrays (LiGAs)^34^, which employ phage virions as a display platform. These genetically-encoded libraries of structurally-diverse multivalent glycoconjugates are powerful probes of GBP–glycan interactions^34^. Compared to monovalent DNA-coded glycan arrays^35–38^, the LiGA strategy provides the ability to control and encode glycan density. LiGA can also probe interactions with cells both in *in vitro* and *in vivo*. In our initial report describing the LiGA technology^34^, we used relatively simple glycan structures, produced through chemical or chemoenzymatic synthesis prior to their incorporation onto M13 phage. Recent advances in the isolation of *N*-glycans from natural sources^39, 40^ and chemoenzymatic synthesis^41^ provides an attractive opportunity for building custom *N*-glycan LiGAs directly on phage.

The Flitsch group has employed biocatalysis for on-DNA glycan synthesis^36^. This pioneering study constructed carbohydrate-based libraries using enzymatic oxidation and/or glycosylation. The glycans generated were only monovalent; nevertheless, this report is an important precedent for chemoenzymatic glycan array synthesis. Later, the Cha and Reichardt groups independently used chemoenzymatic methods to construct multivalent displays of glycans on glass slides^42, 43^. Characterizing the completion of the enzymatic reactions was difficult and the authors inferred that most did not reach full conversion, even when performed under conditions that yielded full conversion of analogous glycans in solution. Most recently, independent reports from Wu and Capicciotti describe enzymatic remodeling of glycans on cells;^44–47^ the major bottleneck in such approaches is reaction characterization. Introduction of a new monosaccharide (e.g., Neu5Ac) on the cell surface is detected by increased binding of the cell to a lectin. However, direct detection of changes directly in cellular glycan composition by mass spectrometry is very challenging^48^. The challenges present in state-of-the-art enzymatic glycosylation on-cells, on-glass, and on-DNA prompted us to develop chemoenzymatic glycan synthesis “on-phage” with a complimentary analytical method to monitor reaction conversion. We anticipated that this approach would enable preparing genetically-encoded multivalent displays of glycans with defined structure and, importantly, quantitatively-defined densities. We describe here the successful implementation of this strategy, which has provided new LiGA components— bacteriophages equipped with DNA-barcodes displaying *N*-glycans. We also observed that on-phage enzymatic conversion in solution occurred more efficiently than on a two-dimensional (glass slide) display. Access to this library has, in turn, allowed us to study glycan–GBPs interactions *in vitro*, on cell surfaces, and in mice, with a particular focus on understanding the impact of glycan density or recognition.

## Results and Discussion

Previously^34^, we assembled a library of glycan-coated phages using strain-promoted azide–alkyne cycloaddition (SPAAC) to ligate oligosaccharides with alkyl-azido linkers to dibenzocyclooctyne (DBCO)-modified M13 phage, each containing a distinct DNA barcode. We chose to employ this approach again to prepare an *N*-glycan library starting using a heterogenous sialylglycopeptide (SGP, **1**, Fig. 1a and Fig.S1-S7) from egg yolk^39, 40^, a commonly employed starting material for chemoenzymatic *N*-glycan preparation^41^.

**Fig. 1:**
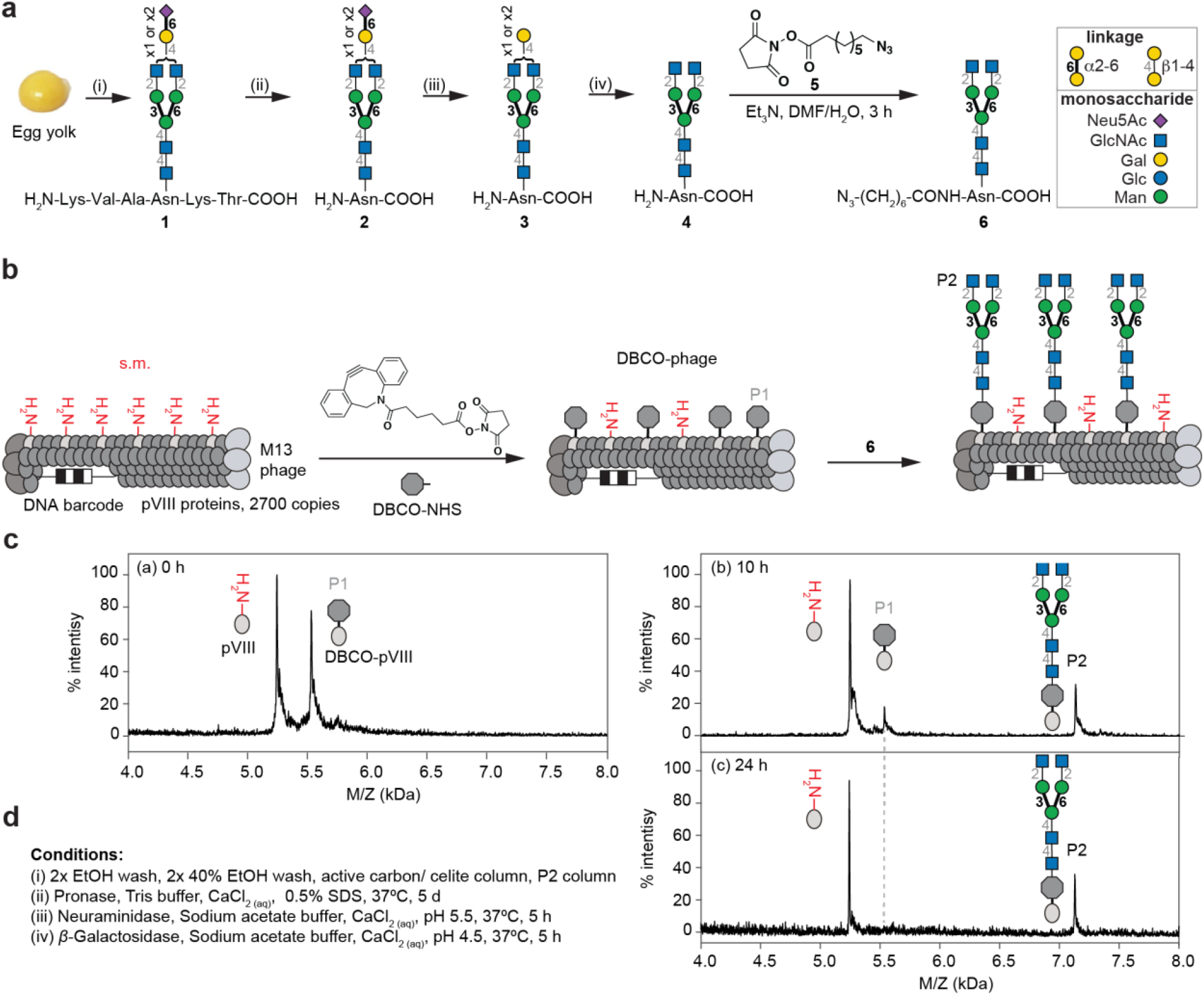
Synthesis and characterization of LiGA components. **a,** Isolation of SGP from egg yolk and trimming steps to afford homogeneous azido-functionalized *N*-glycan **6**. **b,** Representation of two-step linkage of N-glycan to phage. **c,** MALDI-TOF MS characterization of starting material (pVIII protein), alkyne-functionalized product (DBCO-pVIII, P1), and glycoconjugate product (P2). **d,** Conditions of each step to afford product **6**.

Initially, we used a published route^40^ to trim **1** to a homogenous *N*-glycan and ligate it to phage. To do this, **1** was treated with pronase to provide a mixture of asparagine (Asn)-linked biantennary oligosaccharides **2** (Fig. 1a and Fig. S2). Subsequent treatment of **2** with neuraminidase and then β-galactosidase resulted in homogeneous GlcNAc-terminating biantennary structure, **4** (Fig. S3-4), which was *N*-acylated on the amine of Asn with 8-azido-octanoic acid NHS-ester **5**^49^ to yield **6** (Fig. S5). Biantennary glycan **6** retained its natural *N*-linkage to Asn, whereas the azido-linker allowed ligation to DBCO-modified M13 phage by SPAAC. Monitoring the SPAAC reaction by MALDI-TOF MS (Fig. 1c) showed that ligation of *N*-glycan **6** required longer times (24 h) compared to the 1–2 hour reaction times needed for smaller glycans^34^.

We then used similar steps to install a heterogeneous glycosyl asparagine derivative. Thus, Asn-linked *N*-glycans **2** were acylated with **5** to yield a mixture of *N*-glycans **7**, which was ligated to DBCO-modified M13 phage (Fig. 2a). MALDI-TOF MS confirmed the modification (Fig. 2c, d and see Fig. S8 for optimization of MALDI conditions): The peaks S1 and S2 correspond to natural symmetric and asymmetric biantennary structures. The peak S2’ represents the cleavage of one sialic acid from a symmetric glycan during MALDI-TOF MS detection as confirmed by enzymatic treatment described below.

**Fig. 2:**
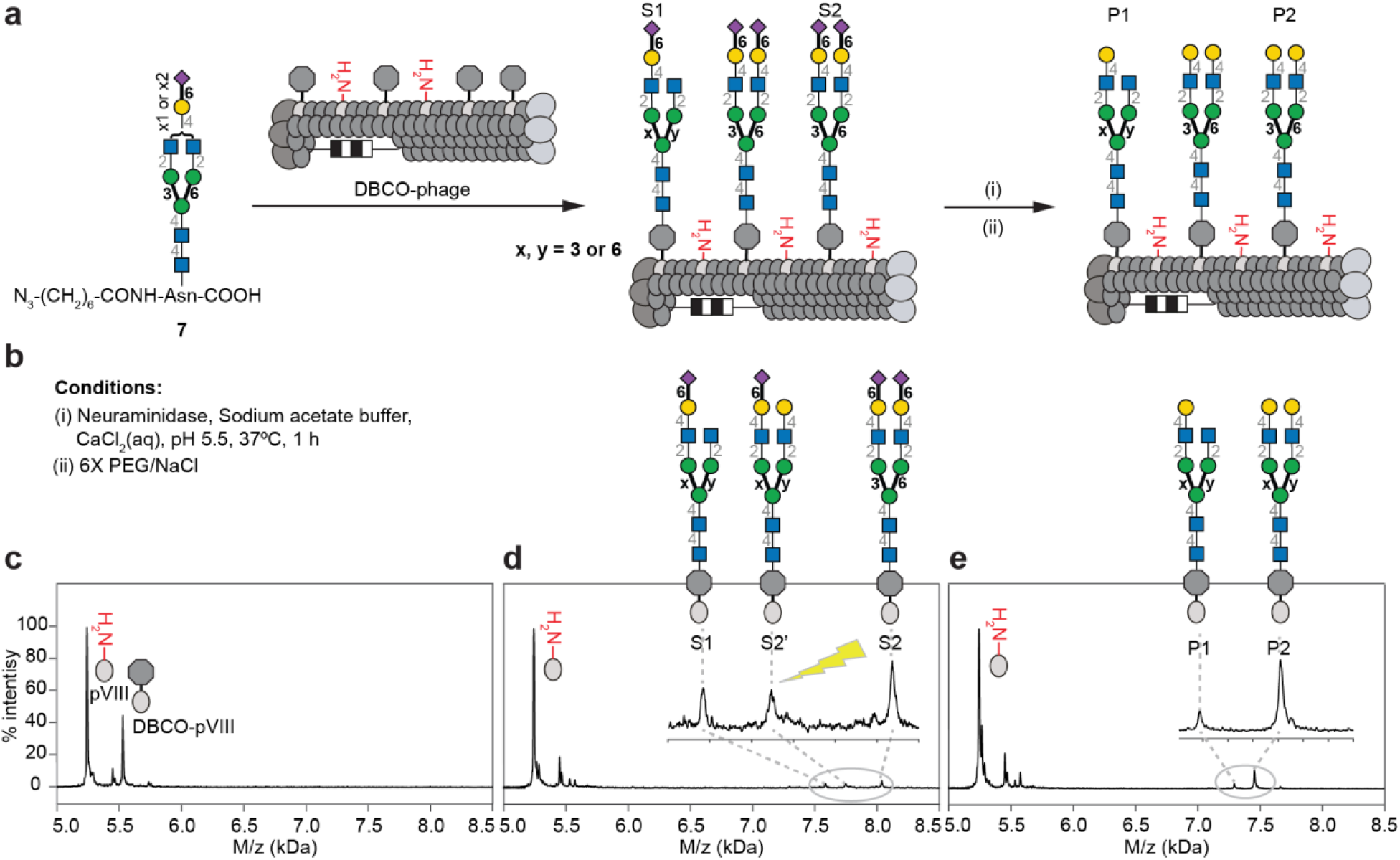
On-phage enzymatic trimming of sialic acid using neuraminidase. **a,** Preparation and attachment of azido-modified Asn-linked biantennary SGP **7** to M13 phage. **b,** neuraminidase treatment to afford mixtures of conjugates with either one or two terminal galactose residues on pVIII. **c,** MALDI-TOF MS characterization of conjugation of DBCO-NHS to pVIII. **d,** S1 and S2 products of ligation of **7**, and S2’, a MALDI-TOF MS artefact. **e,** P1 and P2 phage-displayed glycans generated by neuraminidase treatment.

Phages decorated by either homogeneous or heterogeneous glycans can be used for chemoenzymatic glycan modification. Such on-phage trimming or elongation of glycans facilitates the preparation of glycoconjugates with consistent densities across a range of structures. MALDI-TOF MS confirmed that *β*-galactosidase treatment quantitatively removes terminal galactose residues from glycans on phage (Extended Fig. 1). Similarly, neuraminidase trimming of sialic acids in phage-displayed SGP yielded *N*-glycans with either one or two terminal galactose residues (peaks P1 and P2, Fig. 2e). Subsequent *β*-galactosidase treatment (Fig. 3a) revealed progressive cleavage of both galactose residues with complete disappearance of the symmetric structure after ~2 hours (Fig. 3c), transient accumulation of mono-galactosylated glycans I1, and then their disappearance after ~4 hours (Fig. 3c). Tandem neuraminidase and *β*-galactosidase treatment of heterogeneous **7** on phage, thus, quantitatively gave a homogeneous glycosylated product (Fig. 3a). In contrast, direct *β*-galactosidase treatment of **7** on phage yielded no observable changes (Fig. S9), confirming that the glycan contains no species with terminal galactose residues and that the P2’ peak observed by MALDI-TOF MS are indeed “ghost” species generated by sialic acid cleavage during analysis (Fig. S9). Further evidence comes from model 6’SLN (α-Neu5Ac-(2→6)-LacNAc) glycans ligated to phage, which can be cleaved by *β*-galactosidase; such cleavage was blocked by sialylation of the galactose residues (Extended Data Fig. 2). The homogenous biantennary glycan with terminal *N*-acetylglucosamine (GlcNAc) was further treated with *β-N*-acetylglucosaminidase to cleave the GlcNAc residues yielding a homogenous *N*-glycan structure with terminal mannose (paucimannose) (Extended Data Fig. 3). These results confirm that efficient multi-step enzymatic trimming of *N*-glycans on phage is possible.

**Fig. 3:**
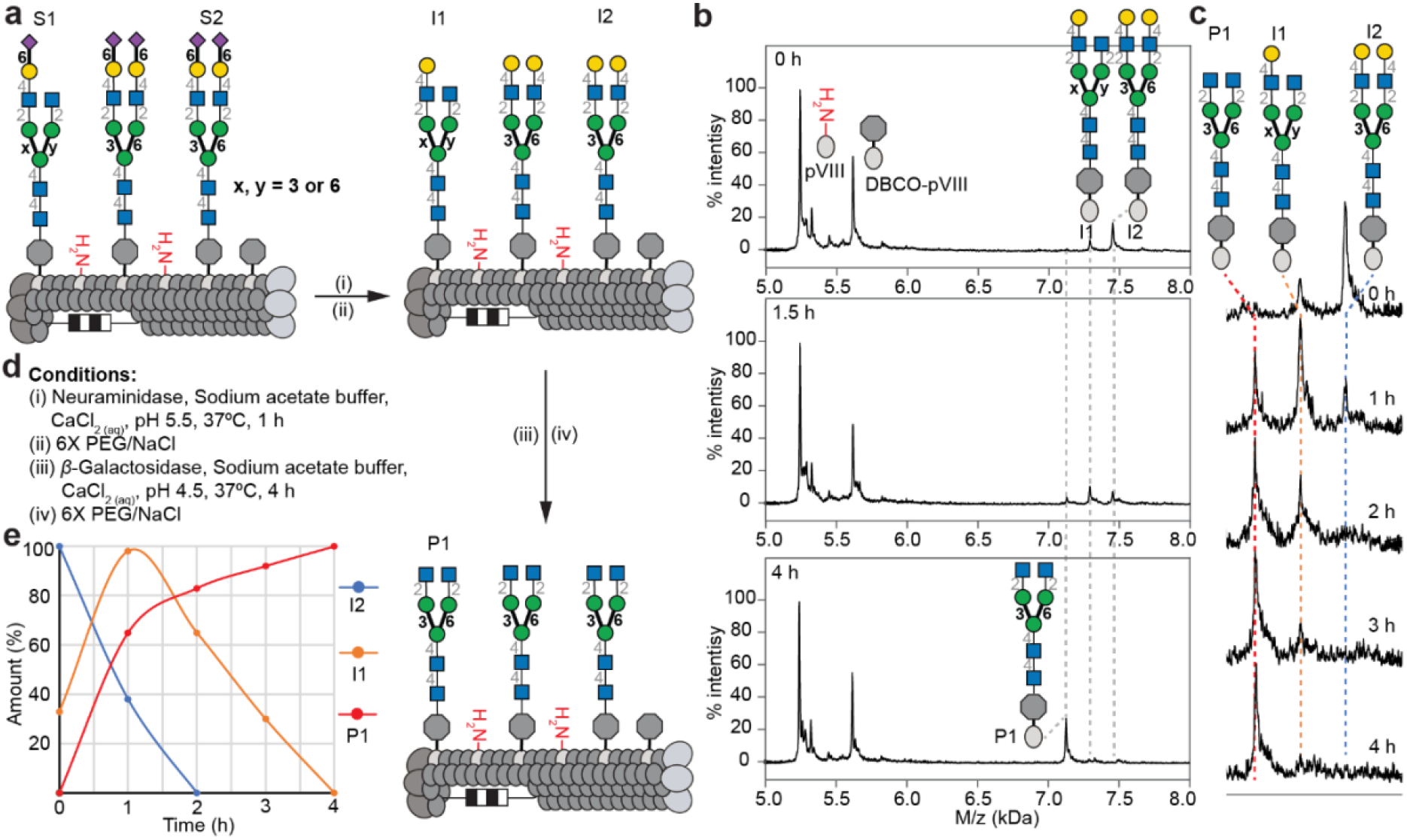
On-phage two-step enzymatic trimming of biantennary *N*-glycans by neuraminidase and β-galactosidase. **a,** Biantennary N-glycans S1, S2 were cleaved by neuraminidase to yield galactose-terminated intermediates I1 and I2. Subsequent treatment with *β*-galactosidase yielded a homogeneous terminal GlcNAc heptasaccharide product P1 on pVIII. **b,** MALDI-TOF MS characterization after *β*-galactosidase treatment. **c,** Progress of β-galactosidase treatment as monitored by MALDI-TOF MS over time. **d,** Conditions of each glycosidase treatment. **e,** Plot of time course results with the amount (%) of I1, I2 and P1 over time (h).

We also explored on-phage *N*-glycan synthesis using glycosyltransferases. In model studies, phages with *N*-glycans terminating with GlcNAc on phage (Extended Data Fig. 4) or in solution (Fig. S7) were treated with *β*-(1→4)-galactosyltransferase (B4GalT1) and uridine 5’-diphosphogalactose (UDP-Gal)^41^ to give, after ~40 hours, phages with lactosamine (LacNAc)-terminating structures (Extended Data Fig. 4b). We also applied these conditions to transfer galactose to heterogeneous asialo-SGP on phage (Extended Data Fig. 5), to provide a homogenous biantennary *N*-glycan with terminal galactose residues. MALDI-TOF MS confirmed a time-resolved conversion of peak P1 (asymmetric glycan) to species P2 (symmetric glycan) over the course of eight hours (Extended Data Fig. 5b). In model studies of sialylation, quantitative addition of sialic acid to LacNAc- and lactose-phages was achieved using recombinant α-(2→6)-sialyltransferase from *Photobacterium damselae* (Pd26ST) and cytidine-5’-monophosphate-*N*-acetylneuraminic acid (CMP-Neu5Ac). After nine hours, α-Neu5Ac-(2→6)-LacNAc and α-Neu5Ac-(2→6)-Lac-phages were formed quantitatively (Extended Data Fig. 2b and 6b). Pd26ST also transferred a sialic acid derivative bearing a 3-butynamide group at C-5^50, 51^ to Lac-phage (Extended Data Fig. 6c). This alkyne handle can be exploited for further chemical derivatization of phage-displayed glycans.

The loss of sialic acid during the MS analysis made it difficult to confirm reaction completion by this technique alone (Extended Data Fig. 2b and 6b). After the sialylation, the phage was therefore treated with *β*-galactosidase to cleave the terminal galactose in any unreacted LacNAc or Lac moieties (Extended Data Fig. 2b and 6g). Using this approach, we confirmed that sialylation proceeds to completion and that asialoglycans observed in MALDI-TOF MS are “ghost” peaks. If necessary, tandem cleavage of Gal and then GlcNAc can also distinguish partially and fully sialylated glycans (Extended Data Fig. 7). We also observed a reduction of MALDI-TOF MS signal intensity upon sialylation. To ensure that this decrease is not due to glycan degradation during the enzymatic reaction, we performed a tandem Pd26ST-catalyzed sialylation and neuraminidase de-sialylation. The intensity for the LacNAc-pVIII conjugate in the mass spectrum was similar before and after the sialylation/de-sialylation cycle (Extended Data Fig. 8) confirming that the decrease in intensity is due to decreased ionization capacity^52^. Using these optimized synthesis and monitoring procedures, we performed a two-step on-phage enzymatic extension using B4GalT1 and Pd26ST to yield a symmetric biantennary sialylated *N*-glycan (Fig. 4). Phage-bound **7** was first extended by B4GalT1 and UDP-Gal (Fig. 4b and 4d). After purification by PEG-precipitation, Pd26ST-catalyzed transfer of Neu5Ac yielded a homogeneous product P2 on phage (Fig. 4b and 4d). These results confirm that multi-step enzymatic glycan extension can be used to create *N*-glycans directly on phage.

**Fig. 4:**
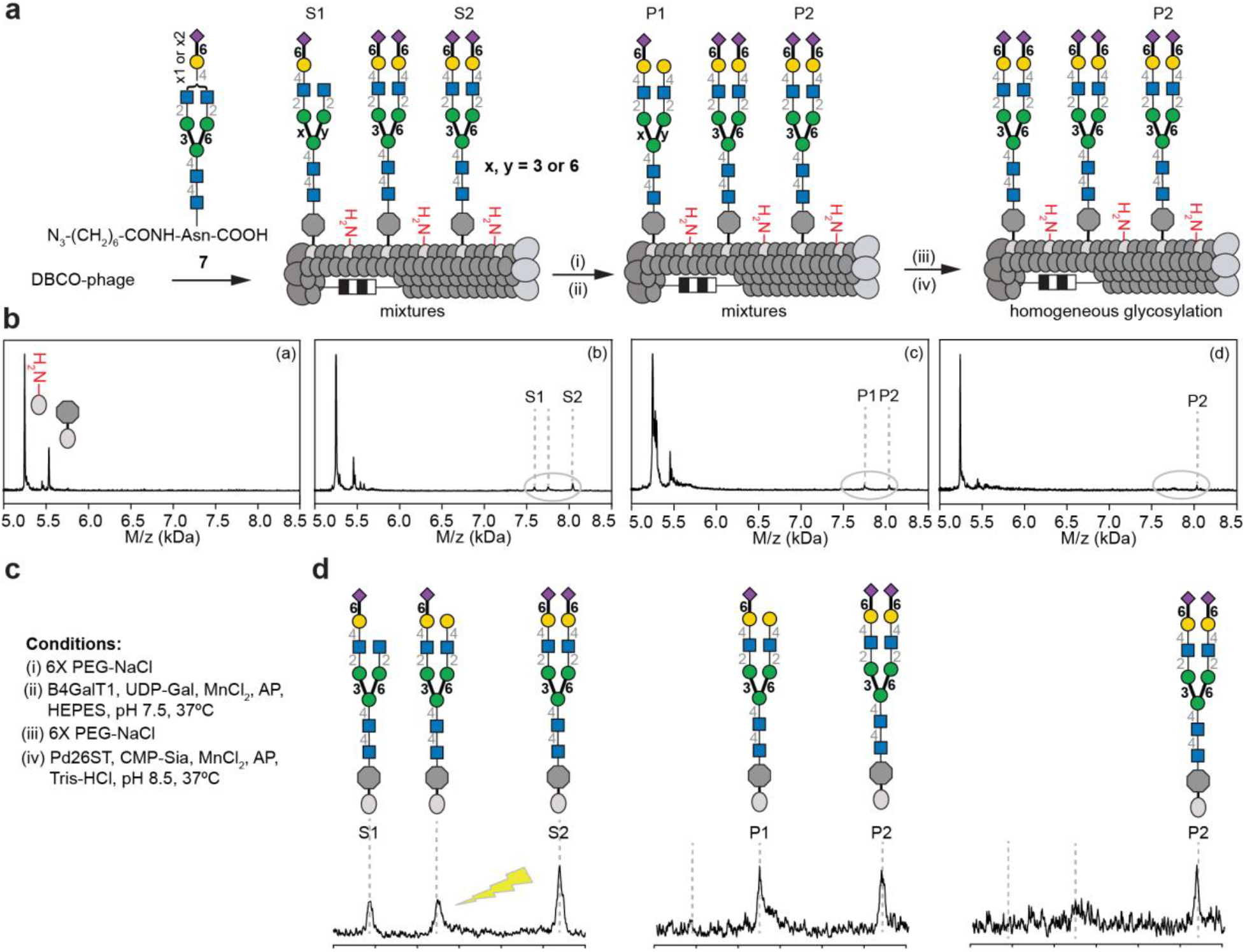
On-phage two-step enzymatic extension of bi-antennary N-glycan by B4GalT1 and Pd26ST. **a,** Sequence of β-1,4-galactosylation and α-2,6-linked sialylation to afford homogeneous sialylated product P2. **b,** MALDI-TOF MS: (b) substrates S1, S2 were products of the ligation reaction. (as described in Fig. 2) (c) B4GalT1-catalyzed galactosylation converted S1 to P1. (d) Pd26ST-catalyzed sialylation converted P1 to P2 yielding a homogeneous display of P2. **c,** Conditions for enzymatic reactions. **d,** Detailed changes of each peak on MALDI-TOF MS. The lightning symbol indicate that these peaks are “ghost” peaks that do not respond to β-galactosidase treatment.

Developing a robust method to synthesize phage-displayed *N*-glycans made it possible to study the effect of *N*-glycan structure and density on GBP binding. To this end, we synthesized a library of six *N*-glycans displayed at five different densities (50, 150, 500, 750 and 1000 glycans/phage) (Fig. S10–S15). An example of our ability to control the glycan density is shown in Extended Data Fig. 9. The density was set by installing a range of 50–1000 copies of DBCO per phage (confirmed by MALDI-TOF MS), followed by complete conjugation with **7** and, finally, quantitative chemoenzymatic conversion of phage-SGP to the desired structures (again confirmed by MS). The resulting library, dubbed “LiGA6×5”, was used to analyze binding to ten lectins: Concanavalin A (ConA), *Sambucus nigra*-I (SNA-I), *Ricinus communis* agglutinin (RCA-I), *Lens culinaris* hemagglutinin (LCA), *Pisum sativum* agglutinin (PSA), *Galanthus nivalis* (GNL), *Erythrina cristagalli* agglutinin (ECL), Galectin-3 (G3C), Wheat Germ agglutinin (WGA) and CD22 (Fig.5 and Fig. S16–S24) as well as cells overexpressing CD22 and DC-SIGN (dentritic cell-specific intercellular adhesion molecule-3-grabbing non-integrin) (Fig. 5).

**Fig. 5:**
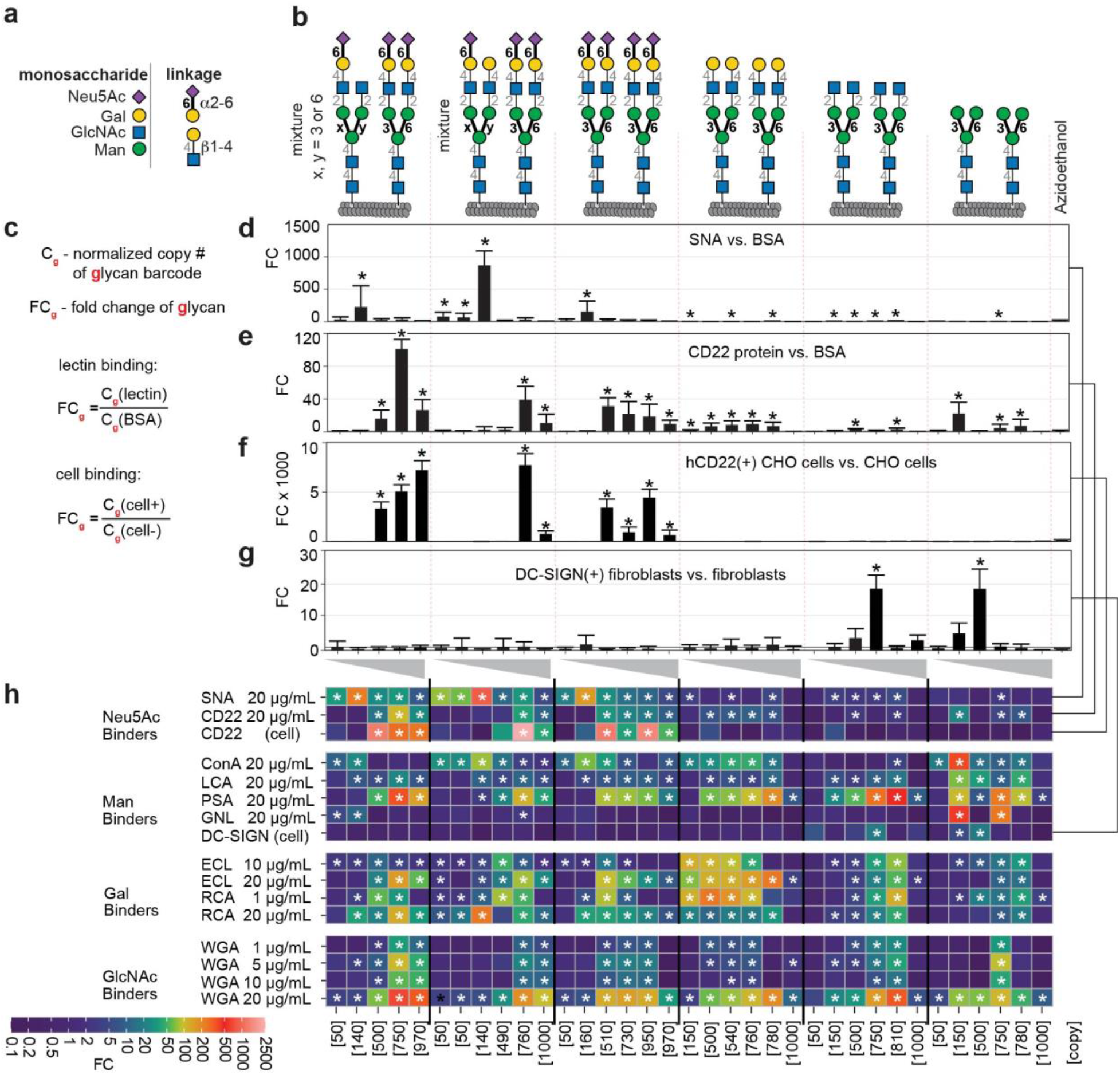
Interaction of LiGA6×5 with SNA, CD22 and cells overexpressing CD22 or DC-SIGN measure affinity- and avidity-based responses. **a,** Glycan abbreviations. **b**, LiGA6×5 is composed of six *N*-glycans each at five display densities. Two of the six are mixtures. **c,** Calculations of the fold change (FC) in experiments. FC was calculated by edgeR; normalization was by assuming equality of azidoethanol labeled phages; false discovery rates (FDR) are from Benjamini-Hochberg adjustment; **d-e,** Binding of LiGA6×5 to wells coated with SNA-I (**d**) or CD22 protein (**e**). **f-g,** Binding of LiGA6×5 to CD22 lectin overexpressed on CHO cells (**d**) and DC-SIGN expressed on rat fibroblasts (**e**); normalization in (**f**) was done using Trimmed Mean of M-values (TMM) due to the low abundance of reference reads in this dataset. **h,** Heatmap representation of the binding of 10 purified lectins and 2 cell-displayed lectins to LiGA6×5. Data from **d-g** (SNA, CD22, CD22^+^ and DC-SIGN^+^ cells) is repeated in the heatmap for consistency. In **d-h**, asterisks (*) indicate FDR≤0.05, n≥5 independent experiments for all data, except n=2 for CD22-cell; in **d-g**, error bars represent standard deviation propagated from the variance of the normalized sequencing data.

In each experiment, LiGA6×5 was incubated in wells coated with each lectin before unbound phages were removed by washing. Bound particles were eluted with 1M HCl and then analyzed by next generation sequencing (NGS). As a metric, we used the fold change (FC) difference^34^ in copy number of each phage with respect to its copy in LiGA6×5 incubated in BSA-coated wells, which underwent analogous processing (wash, elution, NGS).

SNA-I lectin^53^ and CD22 (Siglec-2) both recognize α-(2 →6)-sialylated *N*-glycans and, as expected, we observed interaction of only α-(2 →6)-sialylated components of LiGA with these proteins (Fig. 5d-f and Supplementary Fig. S16, S17). SNA-I bound strongly to a medium density of glycans (~150 glycans per phage), whereas phage with ≤50 or ≥500 glycans bound significantly lower (these phages exhibited minor, statistically significant, binding to SNA-I, at a factor 20–100 fold lower than binding with 150 glycans/phage). In contrast, CD22 required ≥500 glycans per phage for significant binding (Fig. 5e). The same density dependence was seen in a more biologically-relevant environment: CD22 expressed on the surface of cells (Fig. 5f). Asialoglycans exhibited no statistically significant binding to SNA-I and CD22 proteins or CD22^+^ cells at any density. We are not aware of any reports describing non-overlapping density dependence of SNA-I and CD22 proteins, which may stem from differences in the accessibility of protein clusters on the plate or cell surface as well as 50–100x weaker affinity of interaction of sialylated glycans with CD22 when compared to SNA (75 μM and 0.77 μM respectively).^54, 55^

An interplay of structure and density was also observed in a mannose binding lectin family (ConA, LCA, PSA, GNL and DC-SIGN). ConA recognizes a wide range of biantennary *N*-glycans and tolerates multiple extensions,^53^ and LiGA6×5 detected binding of ConA to nearly all biantennary *N*-glycans on phage with the exception of paucimannose with terminal GlcNAc (Fig. 5 and Supplementary Fig. S18). Binding occurred at medium density, 150–750, depending on the glycan, but at 1000 glycans per phage there was no detectable binding to any *N*-glycan. Lack of ConA binding to paucimannose was in agreement with earlier observations^26^ but our data further shows that no binding occurs at any glycan density. Unlike ConA, the mannose binding lectins LCA and PSA bound to the core Man3 epitope in all six *N*-glycans (Fig. 5h and Supplementary Fig. S19, S20). Some glycan array studies have suggested that core fucosylation is required for LCA or PSA to recognize the Man3 epitope^26^ (Supplementary Fig. S19d, S20d) whereas other investigations^53^ observed binding without (Supplementary Fig. S19e, S20e). Affinities of Man3 for LCA (*K*_d_ = 10 μM) and PSA (*K*_d_ = 15 μM) have also been measured by calorimetry.^56^ Notably, PSA recognized α-(2 →6)-sialylated glycans at 1000 glycans/phage whereas binding to asialoglycans at 1000 glycan/phage density was significantly decreased (n = 5 independent experiments). A possible explanation for this observation is a change the conformation/accessibility of Man3 with and without the negatively charged sialic acid epitopes. Finally, the GNL lectin, which is known to recognize paucimannose, bound only to phages that display 150 and 750 copies of this structure and not to any other *N*-glycan at any density (Figure 5h, Supplementary Fig. S21). The double bimodal binding profile could suggest that GNL lectin can bind paucimannose in two different ways, but such hypothesis would have to be confirmed by mode detailed investigations.

Recognition of LiGA6×5 by DC-SIGN was dramatically different from ConA, LCA, PSA or GNL. Using DC-SIGN^+^ rat fibroblasts, we observed that DC-SIGN does not tolerate terminal LacNAc or sialyl-LacNAc on the core Man3, regardless of glycan density (Fig. 5g and 5h). In line with our previous report,^34^ a narrow range of density – 500 Man3 epitopes/phage – was optimal for interaction with DC-SIGN^+^ cells. This preference shifted to higher density for GlcNAc-terminated Man3, corroborating an earlier study^57^ that showed that DC-SIGN recognition requires a high density of GlcNAc-terminated Man3 epitopes.^57^ All glycan binding was ablated at ≥1000 glycans/phage, presumably due to steric occlusion of tightly packed epitopes.^34^

In the Lac/LacNAc binding lectin family, RCA-I lectin^53^ bound to LiGA6×5 components decorated with galactose-terminated *N*-glycans and those with α-(2 →6)-linked sialic acid (Fig. 5h and Supplementary Fig. S23). LiGA measurements matched prior observations^26^ and uncovered a previously unknown bimodal density dependence of RCA-I binding (Supplementary Fig. S23). ECL, which is known to recognize terminal β-(1→4)-linked galactose,^53^ bound selectively at low concentration of lectin ([ECL] =10 μg/mL) to phage displaying terminal galactose. At high concentration ([ECL] = 20 μg/mL), this lectin also recognized phages with α-(2 →6)-sialylated galactose (Fig. 5h and Supplementary Fig. S22). WGA, which binds to various terminal^53^ and internal GlcNAc^58^, exhibited binding to all LiGA components including Man3GlcNAc2 with internal GlcNAc (Fig. 6c, Supplementary Fig. S24). Binding to specific *N*-glycans was dictated by density of the glycan on phage and attenuated by the concentration of WGA used in the experiment (Supplementary Fig. S24). The recognition preferences of ECL and WGA aligned with some earlier glycan array experiments^53^ (Supplementary Fig. S22c, S22e, and S24c, S4e) but diverged from others (Supplementary Fig. S22d, S24d).^26 58^ In these latter cases, the binding of ECL or WGA to sialylated-*N*-glycans was not observed and we note that these lectins bound to sialylated structures in LiGA6×5 only at high density (750–1000 glycans/phage). In contrast, binding of asialoglycans to ECL and WGA was bimodal; binding was optimal at intermediate glycan densities. The complex interplay of glycan density and glycan structure might explain the inconsistent binding preferences between prior glycan array experiments. These LiGA experiments thus emphasize the importance of testing multiple glycan densities as GBP–glycan binding depends on not only on lectin concentration but also on the spatial arrangement (density) of glycans.

**Fig. 6:**
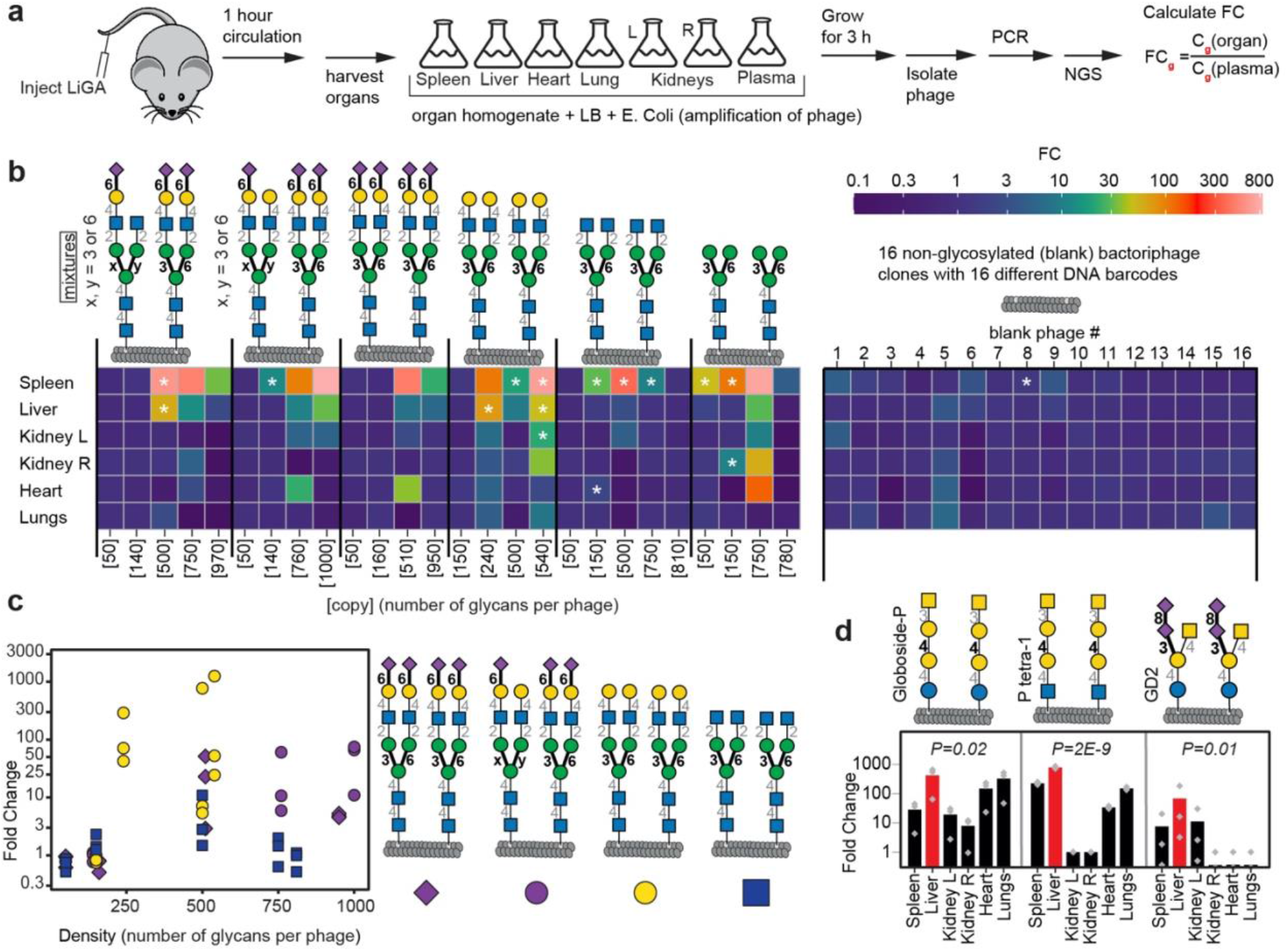
Interaction of LiGA with organs *in vivo*. **a,** LiGA was injected into mice via the tail vein (n=3 animals). After 1 hour, mice were perfused with saline and euthanized. Plasma and organs were collected; each organ homogenate and plasma were incubated with *E. coli* for 3 hours to amplify the phage, followed by PCR and NGS of phage DNA. Fold change (FC) between organ and plasma was calculated by edgeR after normalizing by scaling the unmodified phages to unity, and B-H correction for FDR. **b,** Heatmap of the enrichment of *N*-glycan-decorated and unmodified (blank) phages in organs. Asterisks designate enrichment with FDR ≤0.05. See **Supporting Figs. S25-29** for a detailed summary per organ. **c,** Summary of enrichment of four classes of *N*-glycans in liver, n=3 data points represent data from n=3 animals (data mirrored from heatmap **b**). A Mann-Whittney U-test indicates significant (*P*=0.02) liver enrichment of phage galactose-terminated N-glycans when compared to completely or partially sialylated *N*-glycans or GlcNAc terminated N-glycans **d,** Phage decorated by three glycans with terminal GalNAc that show significant enrichment in liver when compared to other organs, p-values from ANOVA test. See **Supporting Fig. S30** for summary of other glycans enriched in liver (vs. plasma).

Previously we employed LiGA to measure interactions between glycans and receptors on the surface of B cells in live animals.^34^ Here, we injected LiGA into the mouse tail vein (n=3 animals), recovered organs after 1 hour, amplified phages from the organs and plasma and employed NGS of phage populations to determine the structure and density of glycans associated with each organ (Fig. 6 and Supplementary Fig. S25–S30). We compared the Fold Change (FC) difference in copy number and its significance (False Discovery Rate, FDR<0.05) for each glycophage in each organ with respect to the same glycophage recovered from plasma. The injected LiGA contained N-glycan glycophages (components of LiGA6×5 in Fig. 5), previously described glycophages that displayed synthetic glycans^34^, 10 phage clones in which the DBCO handle was capped by azidoethanol and 16 unmodified phage clones. The latter 16+10 “blank” clones served as important baseline of the *in vivo* homing experiment and exhibited only a minor fluctuation in FC across all organs (Fig. 6b). A significant (FDR<0.05) spleen enrichment of diverse N-glycans (Fig. 6b) and synthetic glycans (Fig. S25) was anticipated because lymphocytes, macrophages, dendritic cells, and plasma cells residing in spleen express the most diverse array of cell-surface lectins. In contrast, few N-glycans enriched in kidneys, heart and lungs; high FC ratio of paucimannose-conjugated phage was detected in kidneys and heart but its significance could not be inferred; no significant enrichment of any N-glycan at any density was observed in the lungs. In liver homing, phage particles that displayed various densities of de-sialylated N-glycans with terminal galactose exhibited a significant (*P*=0.02) accumulation in the liver when compared to N-glycans in which galactose was fully or partially capped by sialic acid (Figure 6b-c). Removal of terminal galactose to expose terminal GlcNAc abrogated liver targeting (Figure 6b-c). This observation resembled natural clearance of desialylated red blood cells and platelets by liver. Specifically, aged, desialylated platelets, are cleared by the hepatic Ashwell Morell Receptor (AMR) complex composed of two asialoglycoprotein receptors (ASGPR) 1 and 2^59, 60^. From 12 glycans significantly enriched in liver when compared to plasma (FDR<0.05), three synthetic glycans—P1 tetra (**<u>GalNAc(β1</u>**-3)Gal(α1-4)Gal(β1-4)GlcNAc(β-), Globoside P (**<u>GalNAc(β1</u>**-3) Gal(α1-4)Gal(β1-4)Glc(β-Sp) and GD2 (**<u>GalNAc(β1</u>**-4)ΓNeu5Ac(α2-8)Neu5Ac(α2-3)lGal(β1-4) Glc(β-Sp—enriched in liver significantly more than in any other tested organs (p<0.05). All three contained terminal beta-linked **<u>GalNAc</u>** residue. This observation mirrors well-known delivery of GalNAc-conjugates to ASGR receptors in liver used in FDA-approved drugs (Givlaari) and 28 other GalNAc-conjugated oligonucleotides tested in phases I-III of clinical trials.^61^ Biodistribution of glycophages—components of LiGA—thus mirrors a number of well-known biological mechanisms. These results highlight the possibility of using LiGA to identify both the structure and density of glycans necessary for homing of glycoconjugates to a specific organ paving the route to discovery of new strategies for delivery of therapeutics and uncovering mechanisms that govern glycan-driven biodistribution *in vivo*.

## Discussion

The first LiGA^34^ employed 70–90 small synthetic glycans and represented a powerful new approach to probe glycan–GBP binding. However, interactions of larger glycans with GBPs cannot always be accurately extrapolated from data obtained with smaller structures^62^. In this paper, we address this gap by expanding the LiGA technology to larger molecules, in particular *N*-glycans. To increase synthetic efficiency, we developed “on-phage” enzymatic trimming and extension starting from a readily available *N*-glycan, resulting in a flexible strategy for producing a soluble array in which glycan density can be controlled. A major advantage of this approach is the ability to modify precursors displayed at a defined density leading to products displayed at the same density. Success required developing an analysis strategy where MALDI-TOF MS is used to ensure completion of each enzymatic reaction and to profile and improve reaction conditions. For example, MALDI-TOF MS uncovered that extension and trimming of glycans proceeds more slowly with a higher density glycan precursor. This examination of density vs. enzymatic conversion is the first of its kind and provides insight into enzymatic remodeling of glycans in a complex milieu, here a bacteriophage. Analogous in-situ MS analysis is extremely difficult for on-glass or on-cell synthesis. On-phage enzymatic modification also provides economic benefits: an order of magnitude decrease in the amount of starting glycan needed to manufacture LiGAs compared to glass-based arrays (see Fig. S32).

Using LiGA6×5, we uncovered a previously uncharacterized interplay between *N*-glycan structure and density in lectin recognition not only *in vitro* with pure proteins, but also when they are displayed on cells *ex vivo* and on organs *in vivo*. It has been long postulated that GBP–glycan binding is dictated not only by glycan structure, but also by their density on the cell surface^21, 23, 25^. Nevertheless, efforts to address this issue have been limited to specialized solid-phase arrays^18, 27^ or single glycans displayed on polymers^28^ or liposomes^31^. The approach described here, which involves deploying a range of glycans at different densities in a single experiment, provides a more complete picture of glycan–GBP interactions. Such experiments help to explain discrepancies in previous GBP–glycan binding data originating from different arrays, which we postulate arise, in part, from density effects.

LiGA constructs—glycosylated bacteriophages or “glycophages”—like polymers or liposomes have an optimal ‘soluble’ format suitable for cell-based assays but differ from these more traditional formats, as they can be genetically encoded. Other cell-based glycoarrays^44, 47, 63^ can potentially give rise to similar, DNA-encoded multivalent constructs displayed in a natural milieu. However, precise control of density and composition of a desired glycan structure in cell-based approaches is difficult. LiGA thus represents a heretofore unavailable tool to study GBP–glycan interactions *in vitro* and *in vivo*, which combines ease of manufacture and characterization, with the ability to display diverse synthetic and natural glycans at defined density.

