## Supplementary Information for "Chemoenzymatic Synthesis of Genetically-Encoded Multivalent Liquid *N*-glycan Arrays"

**Figure S32.** Comparison of “on-phage” enzymatic synthesis of liquid glycan array (**a**) with in solution enzymatic synthesis of glycans used to construct glass-based glycan arrays (**b**).50
**References:**........................................................................................................................ 51

**List of abbreviations**:

| | |
|---|---|
| BSA | Bovine Serum Albumin |
| CHO | Chinese Hamster ovary [cells] |
| CD | cluster of differentiation |
| ConA | Concanavalin A |
| CuAAC | Copper-catalyzed azide alkyne cycloaddition |
| DBCO | dibenzylcyclooctyne |
| DNA | Deoxyribonucleotide |
| DMEM | Dulbecco's Modified – Eagle's Medium |
| dsDNA | Double stranded DNA |
| ECL | *Erythrina cristagalli* lectin |
| EDTA | ethylenediaminetetraacetic acid |
| ELISA | Enzyme linked immuno-assay |
| FACS | Fluorescence activated cell sorting |
| Gzip | file format used for file compression / decompression |
| GNL | *Galanthus nivalis* Lectin |
| G3C | Galectin-3 |
| FASTQ | test-based format for storing DNA sequence and its **Q**uality score |
| HEPES | Hydroxyethyl piperazineethanesulfonic acid |
| HRPO | Horseradish peroxidase |
| LB | Lysogeny Broth |
| LCA | *Lens culinaris* agglutinin |
| MALDI-TOF | Matrix-assisted laser desorption/ionization-Time of flight |
| MQ | MilliQ |
| MS | Mass spectrometry |
| MWCO | Molecular weight cutoff |
| NHS | N-hydroxysuccinimide |
| qPCR | Quantitative polymerase chain reaction |
| PBS | Phosphate buffered saline |
| PCR | Polymerase chain reaction |
| PEG | Poly(ethylene glycol) |
| PFU | Plaque forming units |
| | *Photobacterium damselae* |
| PSA | *Pisum sativum* agglutinin |
| RCA-1 | *Ricinus communis* agglutinin I |
| rt | room temperature |
| rSAP. | Shrimp Alkaline Phosphatase |
| SNA-1 | *Sambucus nigra* agglutinin |
| TRIS | Tris(hydroxymethyl)aminomethane |
| WGA | Wheat Germ agglutinin |

### 1. Biochemical Methods

#### **1.1.** *Materials and general information*

Chemical reagents were purchased from Sigma–Aldrich and Thermo Fisher Scientific unless noted otherwise. Pronase from *Streptomyces griseus* was purchased from Roche. Neuraminidase from *Clostridium perfringens* and galactosidase from *Aspergillus niger* were purchased from Sigma–Aldrich. *β-N*-Acetylglucosaminidase S from *Streptococcus pneumoniae* was purchased from New England BioLabs. Glycosyltransferase Pd26ST and B4GalT1 were expressed and purified as described (details below).

Reactions performed in solution were confirmed by LC-MS. For the detection of products, reverse phase high performance liquid chromatography followed by detection using MS (RP-HPLC-MS) was performed using an Agilent 1200 SL HPLC System with a Phenomenex Luna Omega Polar C18, 1.6 µm, 100 Å, 2.1x50 mm column (Phenomenex, Torrance, USA) with guard thermostated at 50 °C. A buffer gradient system composed 0.1% formic acid in water as mobile phase A and 0.1% formic acid in acetonitrile (ACN) as mobile phase B was used. An aliquot of 2 µL of sample was loaded onto the column at a flow rate of 0.5 $mLmin^{-1}$ and an initial buffer composition of 100% mobile phase A and 0% mobile phase B. After injection, the column was washed using the initial loading conditions for 1 min to effectively remove salts. Elution of the analytes was done by using a linear gradient from 0% to 98% mobile phase B over a period of 5.0 min, kept at 98% mobile phase B over a period of 1.5 min and back to the initial condition in 0.5 min. Mass spectra were acquired in positive mode of ionization using an Agilent 6220 Accurate-Mass TOF HPLC/MS system (Santa Clara, CA, USA) equipped with a dual sprayer electrospray ionization source with the second sprayer providing a reference mass solution. Mass correction was performed for every individual spectrum using peaks at m/z 121.0509 and 922.0098 from the reference solution. Mass spectrometric conditions were drying gas 10 L/min at 325 °C, nebulizer 30 psi, mass range 100-3200 Da, acquisition rate of ~1.03 spectra/sec, fragmentor 175 V, skimmer 65 V, capillary 4000 V, instrument state 4 GHz High Resolution. Data analysis was performed using the Agilent Mass Hunter Qualitative Analysis software package version B.07.00 SP2.

MALDI-TOF MS spectra were recorded on AB Sciex Voyager Elite MALDI MS equipped with a MALDI-TOF pulsed nitrogen laser (337 nm) (3 ns pulse up to 300 µJ/pulse) operating in Full Scan MS in positive ionization mode. Nanodrop™ (Thermo Fisher) was used to measure the absorbance of protein and DNA solutions.

High Resolution Mass Spectrometry (HRMS), the electrospray ionization spectra were recorded on an Agilent Technologies 6220 TOF spectrometer with samples dissolved in $CH_3OH$ or $H_2O$.

Concanavalin-A (ConA, #C2010) and Wheat Germ agglutinin (WGA, #L9640) and *Erythrina cristagalli* lectin (ECL, #L5390) were purchased from Sigma–Aldrich. *Galanthus nivalis* lectin (GNL) was purchased from GlycoMatrix (#21510244-1). *Sambucus nigra* agglutinin (SNA-1), *Lens culinaris* agglutinin (LCA) was from EY laboratories, *Pisum sativum* agglutinin (PSA) was from GlycoMatrix, *Ricinus communis* agglutinin I (RCA-I), were a generous gift from Prof. Lara Mahal (University of Alberta).

Sanger sequencing and deep sequencing was performed at the Molecular Biology Service Unit (University of Alberta) using an Illumina NextSeq500 system. All DNA primers were ordered from Integrated DNA Technologies. Biochemical reagents were purchased from Thermo Fisher Scientific unless noted otherwise. HEPES buffer contains 20 mM HEPES, 150 mM NaCl, 2 mM $CaCl_2$, pH 7.4. PBS buffer contains 137 mM NaCl, 10 mM $Na_2HPO_4$, 2.7 mM KCl, pH 7.4. Solutions used for phage work were sterilized by filtration through 0.22 µm filters.

**1.2.** *Extraction, Isolation and Trimming of SGP*

**Sialylated Biantennary Glycopeptide (SGP) Extraction**

SGP **1** was extracted according to reported procedures[1]. Commercially available egg yolk powder (100 g, Rembrandt Foods, Inc) was suspended twice in 95% ethanol (1.5 L) and stirred for 2 h at rt to extract lipids and other organic soluble components. The suspension was filtered, dried via vacuum filtration and the filtrate was discarded to yield an off-white, dry powder. To extract SGP, the off-white powder was suspended twice in aqueous ethanol (40% v/v ethanol, 1.5 L) solution and stirred for 2 h at rt. The insoluble material was discarded and the filtrate was concentrated under reduced pressure at 40 °C. Cold aqueous ethanol (40% v/v ethanol, 0.5 L) was added to the concentrated solution to precipitate proteins, which were removed by centrifugation (3000 rpm, 10 min). The resulting translucent liquid was purified on an active carbon / Celite column (50 g of active carbon and 50 g Celite). Impurities were removed by flushing the column with 100 mL of water (0.1% v/v TFA), 100 mL L of 5% acetonitrile in water (0.1% v/v TFA), and 100 mL 10% acetonitrile in water (0.1% v/v TFA). The SGP was eluted from the column using a solution of 25% acetonitrile in water (0.1% v/v TFA), and fractions containing the product were collected and concentrated under reduced pressure. The resulting white powder was subjected to size-exclusion chromatography (Sephadex$^{TM}$ G-25 Superfine Gel, fine particle size 15–88 µm, column dimensions 5.0 cm x 80 cm, 250 mL fractions) eluting with 0.1 M ammonium bicarbonate to yield SGP **1** as white powder (0.75 mg SGP/g egg yolk powder).

**Trimming and Modification of SGP 1**

The procedures used followed a previous report[1]. Isolated SGP **1** (20 mg) was dissolved in 5 mL Tris buffer (100 mM, pH8.0) containing 5 mM $CaCl_2$. Pronase (Conc. 2 mg/100 µL) from *Streptomyces griseus* (Roche # 10165921001-1G) was added, and the reaction was incubated for 5 days at 37 °C with shaking. The mixture was heated at 80 °C for 20 min followed by Pronase removal using an Amicon Ultra-10 (MWCO-10k) centrifugal filter. The filtrate was lyophilized and purified by size exclusion chromatography (Sephadex$^{TM}$ G-25 Superfine Gel, fine particle size 15–88 µm), eluting with a 0.1 M ammonium bicarbonate. The fractions containing the glycosylated asparagine **2** (7 mg) were collected and lyophilized.

The glycosylated asparagine **2** (7 mg) was dissolved in 1 mL of sodium acetate buffer (50 mM, pH5.5) containing 5 mM $CaCl_2$ and treated with neuraminidase from *Clostridium perfringens* (Sigma–Aldrich, 10 µL, 0.2 units). The reaction was incubated at 37 °C with shaking and monitoring by TLC (*i*PrOH–$NH_4OH$–$H_2O$ = 4:3:1). After 4 h, the mixture was concentrated under reduced pressure and purified by size exclusion chromatography (Sephadex$^{TM}$ G-25 Superfine Gel), eluting with a 0.1 M ammonium bicarbonate. The fractions containing the mixture of symmetric and asymmetric products (3.2 mg) were collected and lyophilized. Products were further dissolved in 0.5 mL of sodium acetate buffer (50 mM, pH4.5) containing 5 mM $CaCl_2$ and *β*-galactosidase (30 µL, 5 units) from *Aspergillus niger* (Sigma-Aldrich) was added to the reaction mixture. The reaction was incubated at 37 °C. The reaction was monitored by TLC (*i*PrOH–$NH_4OH$–$H_2O$ = 4:3:1). After 3 h, the enzymes were removed using an Amicon Ultra-10 (MWCO-10k) centrifugal filter. The filtr.ate was concentrated under reduced pressure and purified by size exclusion chromatography (Sephadex$^{TM}$ G-25 Superfine Gel), eluting with a 0.1 M ammonium bicarbonate. The fractions containing the desired core structure (3.0 mg) was collected and lyophilized. The results were confirmed by LC-MS analysis.

**1.3.** *Expression and purification of enzymes*

**Expression and purification of Histag-Pd26ST ($\alpha$2→6 sialyltransferase) in *E. coli***

A truncated Pd26ST gene encoding for amino acid residues 16–497 of the full-length protein was cloned into a pET15b vector and expressed as a N-His6-tagged protein. LB broth powder (27.5 g) was added to MQ water (1.1 L) and the mixture was agitated until the powder completely dissolved. This LB solution (100 mL) was added to a clean baffled 500 mL Erlenmeyer flask and the remaining solution (1 L) was added to a clean baffled 4 L Erlenmeyer flask. The solutions in both flasks were sterilized by autoclaving for 15 min. Prior to inoculation, ampicillin (100 µL of a 100 µg/mL solution) was added to LB media (100 mL). Then, the plasmid-bearing *E. coli* strain was cultured in LB medium at 37 °C overnight with shaking at 200 rpm. Overexpression of the target protein was achieved by inducing the *E. coli* culture with 0.2 mM of isopropyl 1-thio-$\beta$-D-galactopyranoside (IPTG) when the $OD_{600nm}$ of the culture reached 0.4–0.6 and incubating at 20 °C for 20 h with vigorous shaking at 250 rpm in an incubator shaker. The cell culture was centrifuged at 6,000 RPM (JLA8.1, Beckman Coulter) for 30 min. The cell pellet was resuspended in cold resuspension buffer (50 mM Tris-Cl, 0.2% Triton X-100, 10 mM Imidazole, 300 mM NaCl, pH 8.0). To the resuspended cell pellet protease inhibitor (What inhibitor) cocktail tablet was mixed. The cell suspension was passed through cell disruptor at 20,0000 PSI. The cell lysate was collected and kept at 4 °C. The cell lysate was then centrifuged (36,000 rpm, 60 min, Ti45 rotor) and the supernatant was decanted and stored at 4 °C. The cell lysate containing the His-tagged Pd26ST was passed through pre-equilibrated (5x3 mL, 5 mM imidazole, 0.5 M NaCl, 50 mM Tris-HCl, pH 7.5) Ni-NTA superflow/agarose column. The column was washed (vol, 50 mM Tris-Cl, 20 mM imidazole, 300 mM NaCl, pH 8.0). The His tagged Pd26ST was eluted using elution buffer (vol, 50 mM Tris-Cl, 250 mM imidazole, 300 mM NaCl, pH 8.0). The fractions containing the protein (absorbance at A280) were combined and stored at -20 °C. The final yield was determined to be 1.29 mg/mL.

**Recombinant expression of β-(1→4)-galactosyltransferase 1 (B4GalT1) in *E. coli***

Expression procedures: a single colony of freshly transformed MBPT-HP0826 in E. coli K12 TB1 was grown at 37 °C in LB broth containing 100 µg/ml Ampicillin. After the overnight cultures were inoculated into the scale-up cultures (LB broth containing 100 µg/ml Ampicillin, 0.2% Glucose), the cells were induced with 0.3 mM IPTG at OD600 reaching around 0.5. The culture was incubated at 20 °C with shaking and harvested cells after induction 22 h. The weight of the cell pellets is 12 g/L culture.

Purification procedures: The cell paste (1 L culture) was resuspended with 50 mM HEPES, pH7.5 containing 1 mM EDTA and 300 mM NaCl supplemented with 1 EDTA free protease inhibitor cocktail. Passed through the cell disruptor at 20,000 psi and collected the lysate on ice. The lysate was centrifuged at 40000 rpm for 1 h in order to remove cell debris and membrane proteins. The supernatant was filtered with a Millex 0.22 um GV-13 filter and loaded onto a MBP-Trap HP affinity column (GE) and followed by the washing step until the UV profile go back the baseline. The column was eluted with the elute buffer (50 mM HEPES, pH7.5 containing 300 mM NaCl and10 mM maltose). The eluted fractions were aliquoted and stored at -80 freezer. The yield is 11.6 mg/L culture.

**1.4.** *SDB clone isolation and amplification*

The M13-SDB-SVEKY library described in a previous report[2] was used to isolate the individual phage clones with built-in silent double barcodes. The results of Illumina sequencing of the isolated clones are available at http://ligacloud.ca/ by searching for specific SDB number. Example to search for "SDB25": http://ligacloud.ca/searchDB?search=SDB25

**1.5.** *Analysis of glycosylation of phage samples by MALDI-TOF MS*

Sinapinic acid matrix[3] was formed by deposition of two layers. Layer 1 was prepared from a solution of sinapinic acid (Sigma, #D7927, 10 mg/mL) in acetone–methanol (4:1). Layer 2 was prepared from a solution of sinapinic acid (10 mg/mL) in acetonitrile–water (1:1) with 0.1% TFA. In a typical sample preparation, 2 μL of phage solution in PBS was combined with 4 μL of Layer 2, then a 1:1 mixture of Layer 1: Layer 2+phage was deposited in that order onto the MALDI inlet plate ensuring that Layer 1 was completely dry before adding Layer 2+phage. Spots were washed with 0.1% TFA in water (10 μL) to remove salts from the PBS. To estimate the ratio of modified to unmodified pVIII, we fit and plotted the data using MatLab.

An essential part of LiGA and the LiGA-dictionary (see below) was the copy number of each glycan in the mixture; these numbers were determined for each clone by MALDI-TOF MS. We implemented an automated pipeline for processing of raw MALDI *.txt files to images and integration data. This task was performed by plotMALDI.m MatLab script. Detailed information was described in a previous paper[2].

**1.6.** *Chemical modification of phage clones with glycans to build components of LiGA*

A solution of SDB phage clone ($10^{12}$–$10^{13}$ PFU/mL in PBS) was combined with DCBO–NHS (20 mM in DMF) to afford a 0.2–2.0 mM concentration of DCBO-NHS in the reaction mixture, which typically yields 5–50% of pVIII modification after 45 min of incubation. After conjugation of DBCO-NHS, each clone was individually purified on a Zeba™ Spin Desalting column (7K MWCO, 0.5 mL, Thermo Fisher, #89882) following the manufacturer instructions. Solutions of azido-glycans (10 mM stock in Nuclease Free $H_2O$) were added to the filtrates to afford a 2 mM concentration of the azido-glycan and the solutions were further incubated overnight at 4 °C. All chemical reactions were verified and quantified by MALDI-TOF MS as described in previous sections. If reactions were incomplete and residual pVIII-DBCO peak was detected by MALDI-TOF MS, we added an additional amount of azido-glycan and extended the incubation time. If reactions were complete, the conjugates were purified by on a Zeba™ Spin Desalting column and stored at 4 °C or supplemented with glycerol and stored as a 50% glycerol stock at –20 °C.

**1.7.** *Preparation of LiGA from glycosylated clones*

In a typical protocol, a LiGA was prepared by mixing $10^8$ PFU of desired glycan-phage conjugates in a single tube. The mixture was characterized by titering and $N \times 10^6$ PFU (N = glycan-phage conjugates) was used for a typical lectin or cell-binding experiment. Each unique LiGA mixture was assigned a two-letter identifier (e.g., "SC") and a "dictionary" (e.g., SC.xlsx), a table that describes the correspondence between the DNA barcodes and the glycans in the LiGA mixture (**Table S1**). These dictionaries were subsequently used to translate from nucleotide sequences in the deep-sequencing files to the corresponding glycans (including density). Examples of dictionaries are available as part of the Supporting Information **Table S1**.

In characterizing the LiGA mixture by deep sequencing, we noted that the mixing of the solutions matched by the titer of phage stock did not afford uniform distribution of barcodes after sequencing. The composition of each naïve library or a naïve library binding to a control target, thus, was used as normalization factor in each experiment. Dictionaries for LiGA mixtures relevant to this manuscript are available in Data/LiGA Dictionaries/ folder. Naïve compositions of these mixtures and references to the deep-sequencing data on http://ligacloud.ca are available in **Table S1**. Locations of "silent barcode" (SB) regions SB1 and SB2 in M13 genome is illustrated in a previous published [2] Supplementary Table 1.

To access the data concatenate URL as http://ligacloud.ca/searchLibInfo?f=0&b=0&d=20210430-87EDcaBI-CT

| LiGA Dictionary File | Input composition | Experimental data (Output) | Used in |
|---|---|---|---|
| SC.xlsx | 20211216-87SCbsBI-CT<br>20220219-87SCbsBI-CT | 20211216-87SCsnaBI-CT | SNA-20 µg |
| SC.xlsx | 20211216-87SCbsBI-CT<br>20220219-87SCbsBI-CT | 20211216-87SCkhBI-CT | CD22-20 µg |
| SC.xlsx | 20211216-87SCbsBI-CT<br>20220219-87SCbsBI-CT | 20211216-87SCcaBI-CT<br>20210219-87SCcaBI-CT | ConA-20 µg |
| SC.xlsx | 20211216-87SCbsBI-CT<br>20220219-87SCbsBI-CT | 20210219-87SClaaBI-CT | LCA-20 µg |
| SC.xlsx | 20211216-87SCbsBI-CT<br>20220219-87SCbsBI-CT | 20210219-87SCqbBI-CT | PSA-20 µg |
| SC.xlsx | 20211216-87SCbsBI-CT<br>20220219-87SCbsBI-CT | 20220314-87SCgnlBI-CT | GNL-20 µg |
| SC.xlsx | 20211216-87SCbsBI-CT<br>20220219-87SCbsBI-CT | 20210219-87SCeaaBI-CT | ECL-20 µg |
| SC.xlsx | 20220408-87SCbsBI-CT | 20220408-87SCeaaBI-CT | ECL-10 µg |
| SC.xlsx | 20211216-87SCbsBI-CT<br>20220219-87SCbsBI-CT | 20211216-87SCrcaaBI-CT | RCA-20 µg |
| SC.xlsx | 20220408-87SCbsBI-CT | 20220408-87SCrcaaBI-CT | RCA-1 µg |
| SC.xlsx | 20211216-87SCbsBI-CT<br>20220219-87SCbsBI-CT | 20210219-87SCwgaBI-CT | WGA-20 µg |
| SC.xlsx | 20211216-87SCbsBI-CT<br>20220219-87SCbsBI-CT | 20210314-87SCwgaaBI-CT | WGA-10 µg |
| SC.xlsx | 20211216-87SCbsBI-CT<br>20220219-87SCbsBI-CT | 20210314-87SCwgabBI-CT | WGA-5 µg |
| SC.xlsx | 20211216-87SCbsBI-CT<br>20220219-87SCbsBI-CT | 20210314-87SCwgacBI-CT | WGA-1 µg |
| SC.xlsx | 20211216-87KErbZW-MS | 20211216-87KEkhZW-MS | CD22 on cell |
| SC.xlsx | 20220314-87SCrfZW-MS | 20220314-87SCrdZW-MS | DC-SIGN on cell |

**Table S1:** LiGA mixtures used in this paper.
References to the deep-sequencing data, instructions for online access by URL-concatenation.

**1.8.** *Methods to measure binding of LiGA components*

**Binding of LiGA to lectins immobilized on plate**

The procedures were followed those described previously[2]. Lyophilized Concanavalin A (Sigma-Aldrich, #C2272) was dissolved in PBS at final concentration of 1 mg/mL. The solution was then diluted with PBS to afford a final concentration of 20 μg/mL and 50 μL was added to each well of a 96 well plate (Corning®, #CLS3369). The plate was covered with sealing tape (Thermo Scientific™, #15036) and incubated overnight at 4 °C. The following day, the wells were washed 3x by adding washing buffer (200 μL, 0.1% Tween-20 in PBS) in the wells and discarding the solution by inverting the plate on top of a paper towel. Thereafter, blocking solution (100 μL, 20 μg/uL BSA in PBS) was added to the wells and incubated for 1 h at rt. The solution was discarded by inverting the plate, the wells were then washed three times with washing buffer. After the incubation with blocking solution and triple washing, 50 μL of LiGA6×5 ($8\times10^8$ PFU/mL in PBS) was added to the wells. The solution was incubated for 1 h at rt and discarded by inverting the plate. The wells were washed 2x with washing buffer and 1x with PBS (200 μL). To elute bound phage, 50 μL of HCl (pH 2.0) was added to the well, incubated for 9 min at rt, and the content of each well was transferred to an Eppendorf tube containing 25 μL of 5× Phusion HF buffer (NEB, #M0530S). The neutralized solution was used for titer and as DNA template for PCR and Illumina sequencing. Binding of LiGA to other lectins was performed in similar fashion.

**Binding of LiGA to CHO cells expressing CD22**

Production and maintenance of CHO cells expressing CD22 were described previously[2]. The binding procedure is adapted from a previous publication[2]. Confluent CHO-CD22(+) and CHO-wt cells were detached from culture flask using PBS containing EDTA (5 mM) and centrifuged for 5 min at 300xg. The supernatant was decantated and the pellet was washed twice by resuspending it in PBS (5 mL) and centrifuged for 5 min at 300xg. After final washing, the cell pellet was resuspended in incubation buffer (1% BSA in HEPES buffer) at 2 $x10^6$ cells/mL. The cells were aliquoted (500 μL) to a FACS tube (Corning, #352054), which afforded 1 million cells per FACS tube. Thereafter, LiGA was added to each FACS tube at $10^8$ PFU, which afforded approximately $10^6$ PFU of each clone in the incubation solution. The solution was incubated for 1 h at 4 °C. After incubation, the cells were gently vortexed (Speed 1, Fisher Vortex Genie 2™, #12-812) and wash buffer (3 mL, 0.1% BSA in HEPES buffer) was added to each FACS tube using a small squirt bottle. The solution was centrifuged at 218 ×g for 5 min at 4 °C in a swinging bucket rotor. The supernatant was decanted by inverting the FACS tubes and blotting on a Kimwipe. Two additional washes were performed: during each wash, the tube was filled with 3 mL of wash buffer, centrifuged and inverted to discard the supernatant in same manner as described above. After the last wash, the pellet was resuspended in 1 mL of HEPES buffer, transferred to a microcentrifuge tube and centrifuged for 5 min at 218 ×g at 4 °C in a swinging bucket rotor. The supernatant was discarded by pipetting and the pellet was resuspended in nuclease-free water (30 μL). An aliquot of this solution (2 μL) was sampled and combined with PBS (500 μL, pH 7.4) for tittering. The remaining solution was incubated at 90 °C for 15 min, centrifuged at 21,000 ×g for 10 min, and the supernatant (25 μL) was used as template for PCR reaction as described in PCR protocol section.

### Binding of LiGA to Fibroblasts cells expressing DC-SIGN.

Production and maintenance of rat fibroblast cells expressing DC-SIGN was described previously[2]. The binding procedure is adapted from a previous publication[2]. The Rat-6 fibroblast DC-SIGN(+) and DC-SIGN(–) fibroblast cells were detached from a culture flask using TrypLE (ThermoFisher, # 12605036) and resuspended in incubation buffer (20 mM HEPES, 150 mM NaCl, 2 mM $CaCl_2$, pH 7.4, 1% BSA,) to produce $2x10^6$ cells/mL suspension. The cells were aliquoted (500 µL) to a FACS tube (Corning, #352054), which afforded 1 million cells per FACS tube. Binding of LiGA to these cells was performed using the steps identical to the protocol above mentioned ("**Binding of LiGA to CHO cells expressing CD22**").

**1.9.** *PCR protocol section*

The DNA template solution (25 µL) in nuclease-free water was amplified in total volume of 50 µL with 1x Phusion® buffer, 50 µM of each dNTPs, 500 µM $MgCl_2$, 1 µM forward barcoded primer, 1 µM reverse barcoded primer and one unit of Phusion® High-Fidelity DNA Polymerase (NEB, #M0530S). A typical 50 µL reaction mixture contained:

| | |
|---|---|
| a) 5x Phusion buffer | 10 µL |
| b) 10 mM dNTPs | 1 µL |
| c) Phusion® Polymeras | 0.5 µL |
| d) Forward primer (10 µM) | 2.5 µL |
| e) Reverse primer (10 µM) | 2.5 µL |
| f) Template solution | 25 µL |
| g) Nuclease-free water | 8.5 µL |

Exceptions: in amplification of clonal phages and naïve libraries, the volume of the template (phage solution) was 2 µL. In panning against the intact cells, protein coated beads, and protein coated wells the volume of solution containing the template was 25 µL.

Cycling was performed using the following thermocycler settings:

a) 98°C 3 min,
b) 98°C 10 s,
c) 50 °C 20 s,
d) 72 °C 30 s,
e) repeat b)-d) for 10 cycles,
f) 98 °C 10 s,
g) 72 °C 30s,
h) repeat f)-g) for 20 cycles,
i) 72 °C 5 min, i) 4 °C hold

**1.10.** *Illumina sequencing*

The PCR products described in "1.9. PCR Section" were quantified by 2% (w/v) agarose gel in Tris-Borate-EDTA buffer at 100 volts for ~35 min using a low molecular weight DNA ladder as standard (NEB, #N3233S). PCR products that contain different indexing barcodes were pooled allowing 10 ng of each product in the mixture. The mixture was purified by eGel, quantified by quBit and sequenced using the Illumina NextSeq paired-end 500/550 High Output Kit v2.5 (2x75

Cycles). Data was automatically uploaded to BaseSpace™ Sequence Hub. Processing of the data is described in section "1.12. Processing of Illumina data".

**1.11.** *General data processing methods*

General data processing methods were reported in a previous paper[2]. Data analysis was performed in Python, Matlab or R/Bioconductor. Comparison and testing differences for significance in LiGA data was performed essentially as differential enrichment analyses of phage displayed library sequencing described in our previous reports[4, 5] and based on differential expression (DE) analysis implemented in edgeR[4, 5]. In DE-analysis three factors were considered: (i) modeling of the observed counts using a negative binomial model; (ii) Benjamini–Hochberg (BH) adjustment to control the false discovery rate (FDR) at $\alpha = 0.05$[5]; (iii) normalization of data across multiple samples using the Trimmed Mean of M-values (TMM) normalization[5]. Core scripts are available as part of the supporting information or on GitHub.

To assess the significance of a glycan binding in a specific experiment the differential enrichment of the levels of the DNA barcode associated with that glycan in "test" sets of DNA read was compared to the levels of the same read in "control" sets. For example, in cell-based experiments, the "test" dataset was association of LiGA with receptor (+) cell line, whereas the control was the dataset was association of the identical LiGA with isogenic cell line that contained no target receptor. In binding to lectins, the "control" dataset was association of LiGA with blank carriers (beads or plate). Prior to DE-analysis, "test" and "control" data sets were retrieved from the http://ligacloud.ca/ server as tables of glycans, DNA, and raw sequencing counts. DNA reads that could not be mapped to any entries in LiGA dictionary were discarded.

**1.12.** *Processing of Illumina data*

The processing of FASTQ files downloaded from BaseSpace™ Sequence Hub to reads and frequencies of these glycans was performed as previously described[6-8]. A LiGA-specific lookup table ("LiGA dictionary") was used to convert the identified SDB to glycans and display density. Abbreviated names of glycans are based on those recommended by the Consortium for Functional Glycomics:
http://www.functionalglycomics.org/static/consortium/resources/resourcecored2.shtml

Translated files with raw DNA reads, raw counts, and mapped glycans were uploaded to the http://ligacloud.ca/ server. All LiGA sequencing data is publicly available at http://ligacloud.ca/ server. Each experiment has a unique alphanumeric name (e.g., **20180711-87YOrdRB-JM**) and unique static URL (Highlighted portion of the URL is constant for all files):
http://ligacloud.ca/searchLibInfo?f=0&b=0&d=20180711-87YOrdRB-JM

**1.13.** *Panning of LiGA in Mice*

All procedures and experiments involving animals were carried out using a protocol approved by the Health Sciences Laboratory Animal Services (HSLAS) at the University of Alberta. The protocol was approved as per the Canadian Council on Animal Care (CCAC) guidelines. All mice were maintained in pathogen-free conditions at the University of Alberta breeding facility.

Animals were injected with LiGA (0.2 mL, 1 x $10^{11}$ PFU/mL in PBS). One-hour post-injection mice were euthanized with $CO_2$ and blood (0.5 mL) was drawn out and stored on ice. Internal organs (heart, liver, kidney, lung and spleen) were collected and stored in cold DMEM (Thermo Fisher). Tissues were homogenized by grinding between 75 mm frosted microscope slides. Homogenized tissues of each organ were transferred into 25 mL LB and supplemented with a 0.5 mL of log phase *E. coli* K12 ER2738. After incubation at 37 °C for 3 h, the amplified phage in the culture were isolated by centrifugation at 4500 g for 10 min. The supernatant was incubated with 5% PEG-8000, 0.5 M NaCl for 8 h at 4 °C, followed by 30 min centrifugation at 13, 000 ×g. The phage pellet was re-suspended in 1 mL PBS-Glycerol 50% and solution of phages (2 µL) was PCR amplified using barcoded sequencing primers using a protocol described in *"PCR Protocol Section*" and analyzed by Illumina sequencing.

### 2. Synthetic methods

#### **2.1.** *General Synthetic methods*

All reagents were purchased from commercial sources and used without further purification. Oven-dried glassware was used for all reactions. Reaction solvents were dried by passage through columns of alumina and copper under nitrogen. All reactions, unless stated otherwise, were carried out at rt under positive pressure of argon. Organic solutions were concentrated under vacuum below 40 °C using a rotary evaporator. Reaction progress was monitored by TLC on Silica Gel 60 $F_{254}$ (0.25 mm, E. Merck). Visualization of the TLC spots was done either under UV light or charring TLC plates with acidified *p*-anisaldehyde solution in ethanol or phosphomolybdic acid stain. Column chromatography was performed using Silica Gel 40–60 µM. $^{1}$H NMR spectra were recorded using 500 MHz or 400 MHz instruments, and the data are reported as if they were first order. $^{13}$C NMR (APT) spectra were recorded at 125 MHz. The details of LC-MS and HRMS was described in Section 1.1.

#### **2.2.** *Acylation of the asparagine amino group with 8-azido-octanoic acid*

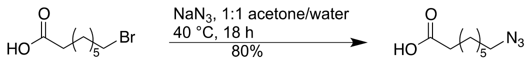

To a stirred solution of 8-bromooctanoic acid (2.00 g, 8.96 mmol) in acetone–water (1:1, 5 mL) was added sodium azide (1.16 g, 17.8 mmol). The solution was heated to 40 °C and stirred for 18 h. After cooling to rt, EtOAc (10 mL) was added and the organic phase separated from the aqueous phase. The aqueous phase was extracted with EtOAc (3 × 10 mL) and the organic phase and extractions were combined, washed with water (5 mL) and brine (5 mL), before being dried ($MgSO_4$) and filtered. The filtrate was concentrated to give 8-azidooctanoic acid as a colourless oil (1.28 g, 80%). $R_f$ 0.5 (3:1 hexane–EtOAc). This procedure was adapted from the reported literature[9]. HR ESIMS: *m/z* [M–H$^-$] calcd for $C_8H_{14}N_3O_2$: 184.1092. Found: 184.1093.

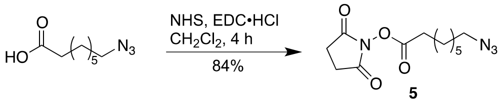

1-Ethyl-3-(3-dimethylaminopropyl)carbodiimide (EDC, 1.59 g, 8.3 mmol) was added to a suspension of 8-azidooctanoic acid (1.28 g, 6.9 mmol) and *N*-hydroxysuccinimide (NHS, 0.95 g, 8.3 mmol) in $CH_2Cl_2$ (35 mL) at rt, and the reaction was monitored by TLC (EtOAc–hexane, 1:3). After 5 h, 1 N HCl was added and the organic layer was separated, washed with saturated aqueous sodium bicarbonate, water, dried over sodium sulfate and filtered. The filtrate was concentrated and the resulting residue was purified by column chromatography (EtOAc–hexane, 1:2) to give 8-azidooctanoic acid NHS ester (**5**) as a colorless oil (1.08 g, 84%). $R_f$ 0.3 (3:1 hexane–EtOAc). This procedures were followed according to a reported literature[9]. HR ESIMS: *m/z* [M+Na$^+$] calcd for $C_{12}H_{18}N_4NaO_4$: 305.1220. Found: 305.1228.

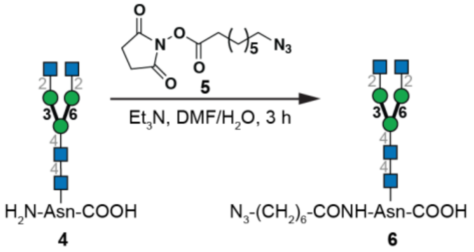

To a solution of **4** (2 mg, 0.001 mmol) in DMF (1.0 mL) and $H_2O$ (added dropwise until the solution turned clear) at rt was added 8-azidooctanoic acid NHS ester **5** (1 mg, 0.004 mmol). Triethylamine was added dropwise to make ensure the reaction was basic by using pH test paper rolls. The reaction was monitored by TLC (*i*-PrOH–$NH_4OH$–$H_2O$, 4:3:1). After 3 h, the solution was concentrated and the residue was purified by size exclusion chromatography (Sephadex$^{TM}$ G-25 Superfine Gel). The fractions containing **6** (1.6 mg) ($R_f$ 0.6, *i*-PrOH–$NH_4OH$–$H_2O$, 4:3:1) were collected and lyophilized. The results were confirmed by LC-MS (**Fig. S5**) and NMR analysis. Note: The reaction of NHS esters with amines is pH-dependent. At lower pH, the amino group is protonated, and no modification takes place. At higher-than-optimal pH, hydrolysis of the NHS ester is rapid and modification yield diminishes. The optimal pH value for this modification is 8.3–8.5. We have attempted the reaction in sodium bicarbonate buffer (pH=8.5); however, acylation proceeded slowly and the NHS ester was hydrolyzed. The use of trimethylamine in DMF with a small amount of $H_2O$ avoided these issues.

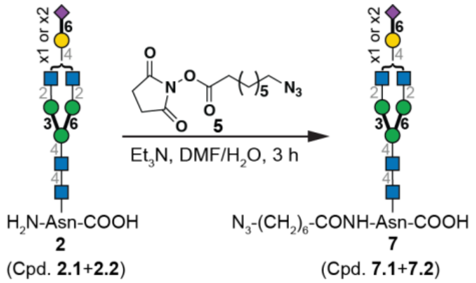

To a solution of heterogeneous *N*-glycans **2** (2 mg) in DMF (1.0 mL) and $H_2O$ (added dropwise until the solution turned clear) at rt was added 8-azidooctanoic acid NHS ester **5** (1 mg, 0.004 mmol). Triethylamine was added dropwise to make sure the reaction was basic by pH test paper rolls. The reaction was monitored by TLC until the spots of starting material have consumed. After 3 h, the solution was concentrated and the residue was purified by size exclusion chromatography (Sephadex™ G-25 Superfine Gel). The fractions containing the desired azido- *N*-glycans **7** (1.4 mg) was collected and lyophilized. The results were confirmed by LC-MASS (**Fig. S6**).

### 3. On-Phage enzymatic glycan modification

#### **3.1.** *Procedures for enzymatic reactions on the phage surface*

**Procedures for phage surface glycosidase treatment:**

*Cleavage of sialic acid in phage-displayed N-glycans by neuraminidase*
After modification of phage clones with **7** to afford the conjugated phage (60 µL, ~$10^{12}$–$10^{13}$ PFU/mL in PBS, pH7.4), 5% PEG-8000, 0.5 M NaCl (12 µL) was added and the solution was kept for 1.5 h at 0 °C. The solution was then centrifuged at 21000 g for 10 min at rt. The supernatant was removed, and the pellet was resuspended in sodium acetate buffer (40 µL, 50 mM, pH 5.5 containing 5 mM $CaCl_2$) followed by the addition of *Clostridium perfringens* neuraminidase (3 µL, 0.07 units). The reaction was incubated at 37 °C and monitored by MALDI-TOF MS. The reaction was complete in 1 h to afford galactose-terminating *N*-glycans (**Fig. 3**) on the phage surface.

*Cleavage of galactose in phage-displayed N-glycans by β-galactosidase*
To a solution of phage modified with terminal galactose containing mixtures of *N*-glycans I1 and I2 in **Fig. 4**, 5% PEG-8000, 0.5 M NaCl (12 µL) was added and the solution was kept for 1.5 h at 0 °C. The solution was centrifuged at 21000 g for 10 min at rt. The supernatant was removed, and the pellet was resuspended in sodium acetate buffer (40 µL, 50 mM, pH 4.5 containing 5 mM $CaCl_2$) followed by the addition of *Aspergillus niger* β-galactosidase (2 µL , 0.47 units). The reaction was incubated at 37 °C and monitored by MALDI-TOF MS every 1 h. The reaction was complete in 4 h to afford a symmetric biantennary GlcNAc-terminating *N*-glycan on the phage surface.

*Cleavage of acetylglucosamine in phage-displayed N-glycans by β-N-acetylglucosaminidase S*
To a solution of phage modified with the symmetric biantennary terminal GlcNAc *N*-glycan in **Extended Data Fig. 1**, 5% PEG-8000, 0.5 M NaCl (12 µL) was added and the solution was kept for 1.5 h at 0 °C. The solution was then centrifuged at 21000 g for 10 min at room temperature. The supernatant was removed, and the pellet was resuspended in nuclease free water (35 µL) and glycobuffer (4 µL, New England BioLabs Inc. #P0744S) followed by the addition of *Streptococcus pneumoniae β-N*-Acetylglucosaminidase S (1 µL, 4 units, New Englang BioLabs Inc. #P0744S). The reaction was incubated at 37 °C and monitored by MALDI-TOF MS. The reaction was complete in 1 h to afford mannose-terminating *N*-glycan on the phage surface.

**Procedures for phage surface glycosylation:**

*Installation of α-(2→6)-linked sialic acid using Pd26ST*
The conditions described below were used in the model study (**Fig. S15(B)**). After modification of phage clones with LacNAc (β-Gal-(1→4)-β-GlcNAc-$OCH_2CH_2N_3$) to afford the conjugated phage (60 µL, ~$10^{12}$–$10^{13}$ PFU/mL in PBS, pH 7.4), 5% PEG-8000, 0.5 M NaCl (12 µL) was added and the solution was kept for 1.5 h at 0 °C. After 1.5 h, the solution was centrifuged at 21000 g for 10 min at rt and supernatant was decanted. The pellet was resuspended in Tris-HCl buffer

(25 μL, 100 mM with 20 mM $MnCl_2$, pH8.5) containing CMP-Neu5Ac (300 μg), recombinant Shrimp Alkaline Phosphatase (rSAP, 0.5 μL) and Pd26ST (10 μL, 1.29 mg/mL). The reaction mixture was incubated at 37 °C and monitored by MALDI-TOF MS. Pd26ST (5 μL, 1.5 mg/mL) was added twice at 3 h and 6 h reaction time. The reaction was complete in 9 h and was validated by a β-galactosidase digestion experiment (**Extended Data Fig. 2**).

*Installation of β-(1→4)-linked galactose using B4GalT1*

The condition described below was used in the model study (**Extended Data Fig. 4**). After modification of phage clones with *N*-Acetylglucosamine (β-GlcNAc-$OCH_2CH_2N_3$) to afford the conjugated phage (60 μL, ~$10^{12}$-$10^{13}$ PFU/mL in PBS, pH7.4), 5% PEG-8000, 0.5 M NaCl (12 μL) was added and the solution was kept for 1.5 h at 0 °C. After 1.5 h, the solution was centrifuged at 21000 g for 10 min at rt and the supernatant was decanted. The pellet was resuspended in HEPES buffer (20 μL, 50 mM with 10 mM $MnCl_2$, pH 7.5) containing UDP-Gal (100 μg), Shrimp Alkaline Phosphatase (1 μL) and B4GalT1 (15 μL, 0.048 mg/mL). The reaction mixture was incubated at 37 °C and monitored by MALDI-TOF MS. The reaction was complete in 40 h and was validated by MALDI-TOF MS.

**a**

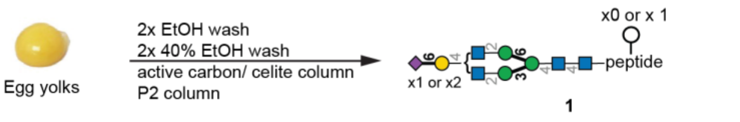

(Compounds. **1.1**+**1.2**+**1.3**+**1.4**)

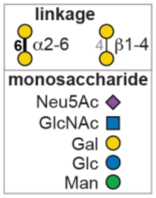

**b**

**Compound table**

| Compound Label | RT | Mass | Abund | Formula | Tgt Mass | Diff (ppm) |
|---|---|---|---|---|---|---|
| Compound1.1:C95H162N14O57 | 2.42 | 2411.0174 | 8317 | C95H162N14O57 | 2411.0208 | -1.43 |
| Compound1.2:C112H189N15O70 | 2.45 | 2864.1626 | 47419 | C112H189N15O70 | 2864.1691 | -2.25 |
| Compound1.3:C101H172N14O62 | 2.45 | 2573.071 | 6892 | C101H172N14O62 | 2573.0736 | -1.01 |
| Compound1.4:C118H199N15O7 | 2.47 | 3026.2159 | 6213 | C118H199N15O75 | 3026.2219 | -1.99 |

Compound 1.1

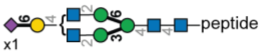

peptide = $NH_2$-Lys-Val-Ala-Asn-Lys-Thr-COOH

| Compound 1.1 | *m/z* | RT | Algorithm | Mass |
|---|---|---|---|---|
| $C_{95}H_{162}N_{14}O_{57}$ | 805.014 | 2.42 | Find by Formula | 2411.017 |

**Compound Chromatograms**

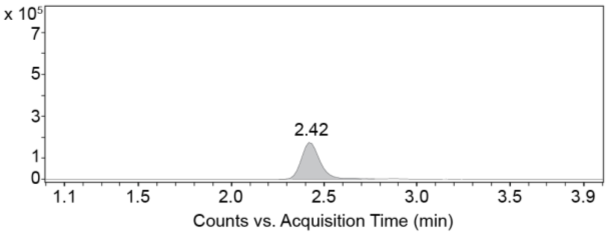

Compound 1.3

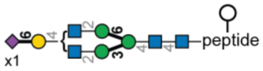

| Compound 1.3 | *m/z* | RT | Algorithm | Mass |
|---|---|---|---|---|
| $C_{101}H_{172}N_{14}O_{62}$ | 859.032 | 2.45 | Find by Formula | 2573.071 |

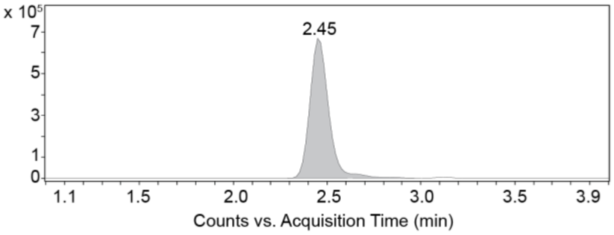

Compound 1.2

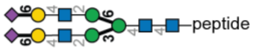

| Compound 1.2 | *m/z* | RT | Algorithm | Mass |
|---|---|---|---|---|
| $C_{112}H_{189}N_{15}O_{70}$ | 956.0627 | 2.45 | Find by Formula | 2864.1626 |

**Compound Chromatograms**

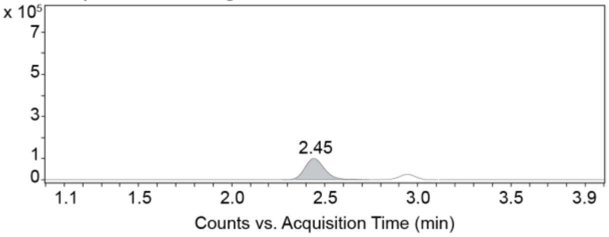

Compound 1.4

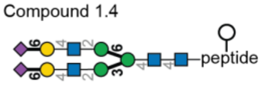

| Compound 1.4 | *m/z* | RT | Algorithm | Mass |
|---|---|---|---|---|
| $C_{118}H_{199}N_{15}O_{7}$ | 1010.0805 | 2.47 | Find by Formula | 3026.2159 |

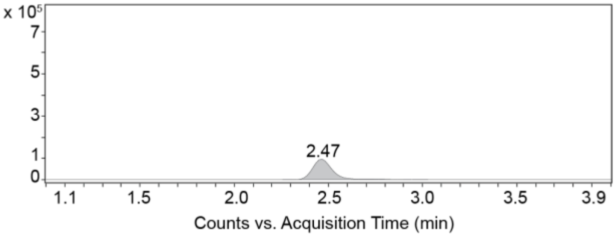

**Figure S1.** LC-MS analysis of isolated SGP **1**.

**a,** Scheme of SGP production from egg yolk powder. **b,** Analytical LC-MS chromatogram of isolated SGP showing four components, Compounds 1.1–1.4 with different retention times.

**a**

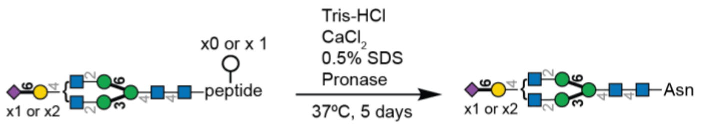

**1**

**2**
(Compounds. **2.1**+**2.2**)

**Compound table**

| Compound Label | RT | Mass | Abund | Formula | Tgt Mass | Diff (ppm) |
|---|---|---|---|---|---|---|
| Compound2.1:C88H144N8O64 | 6.08 | 2336.8233 | 1279 | C88H144N8O64 | 2336.8259 | 1.11 |
| Compound2.2:C71H117N7O51 | 5.09 | 1883.674 | 850 | C71H117N7O51 | 1883.6777 | 1.95 |

**b**

Compound 2.1

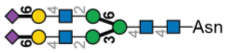

Compound 2.2

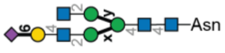

**x, y = 3 or 6**

| Compound 2.1 | *m/z* | RT | Algorithm | Mass |
|---|---|---|---|---|
| $C_{88}H_{144}N_8O_{64}$ | 780.2822 | 6.08 | Find by Formula | 2336.8233 |

| Compound 2.2 | *m/z* | RT | Algorithm | Mass |
|---|---|---|---|---|
| $C_{71}H_{117}N_7O_{51}$ | 942.8444 | 5.09 | Find by Formula | 1883.674 |

**Compound Chromatograms**

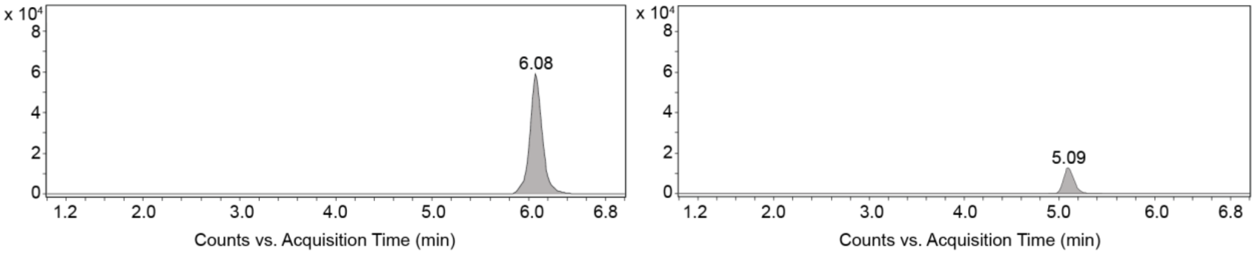

**Figure S2.** LC-MS results of pronase treated SGP **2** after purification.
**a,** Scheme for treating SGP **1** with pronase to trim the peptide down to Asparagine (Asn). **b,** LC-MS analysis of pronase treated SGP **2** showing two components with different retention times. Compound 2.1 is a homogeneous product with terminal sialic acid and Compound 2.2 is a mixture of heterogeneous structures.

**a**

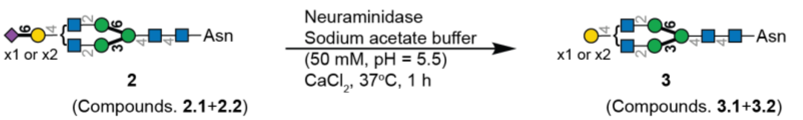

**b**

**Compound table**

| Compound Label | RT | Mass | Abund | Formula | Tgt Mass | Diff (ppm) |
|---|---|---|---|---|---|---|
| Compound3.1: C60H100N6O43 | 5.80 | 1592.5823 | 3741 | C60H100N6O43 | 1592.5823 | -0.02 |
| Compound3.2: C66H110N6O48 | 6.04 | 1754.6327 | 12078 | C66H110N6O48 | 1754.6351 | -1.34 |

Compound 3.1

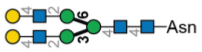

Compound 3.2

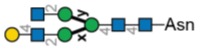

**x, y = 3 or 6**

| Compound 3.1 | *m/z* | RT | Algorithm | Mass |
|---|---|---|---|---|
| $C_{88}H_{144}N_8O_{64}$ | 797.2912 | 5.80 | Find by Formula | 1592.5823 |

| Compound 3.2 | *m/z* | RT | Algorithm | Mass |
|---|---|---|---|---|
| $C_{71}H_{117}N_7O_{51}$ | 878.3164 | 6.04 | Find by Formula | 1754.6327 |

**Compound Chromatograms**

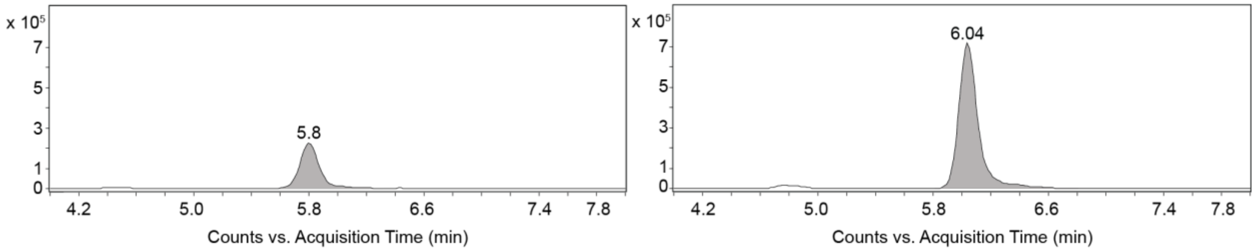

**Figure S3.** LC-MS results after neuraminidase treatment of **2**, giving **3**.
**a,** Scheme for cleaving the non-reducing end sialic acid in **2** by neuraminidase to give a hetereogenous mixture **3** (Compounds 3.1 and 3.2). **b,** LC-MS analysis showing two components with different retention times.

**a**

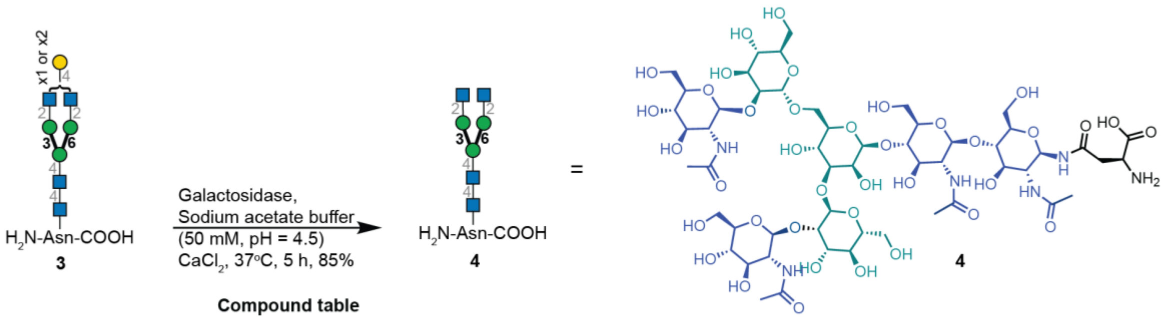

**b**

**Compound table**

| Compound Label | m/z | RT | Algorithm | Mass |
|---|---|---|---|---|
| C54H90N6O38 | 716.2722 | 5.33 | Find by Formula | 1431.5368 |

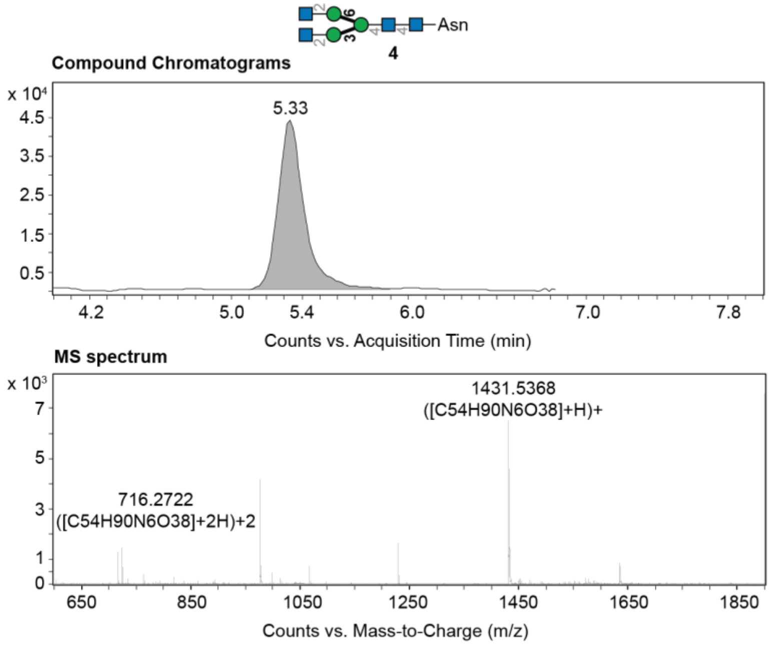

**Figure S4.** LC-MS results after β-galactosidase treatment of **3**, giving **4**.
**a,** Scheme for cleaving the non-reducing end galactose residues in **3** by β-galactosidase to give a homogenous biantennary *N*-glycan terminating in GlcNAc **3. b,** LC-MS result of compound **4**.

**a**

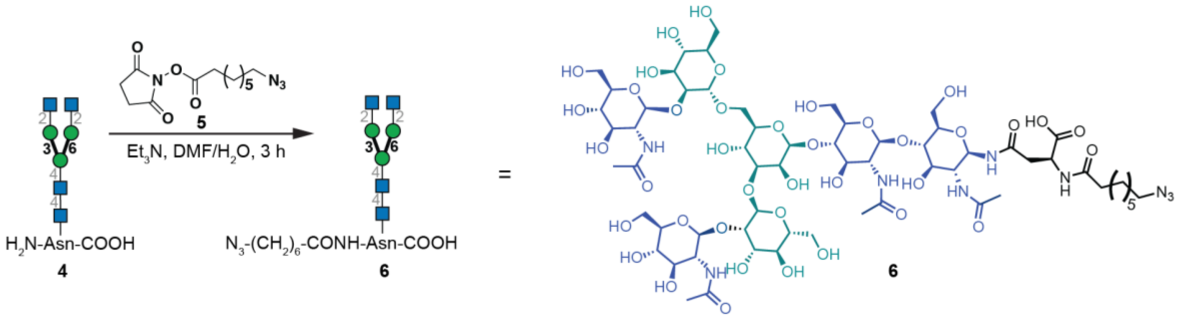

**b**

**Compound table**

| Compound Label | m/z | Tgt Mass | Algorithm | Diff (ppm) |
|---|---|---|---|---|
| C62H103N9O39 | 1597.6365 | 1597.6365 | Find by Formula | -0.76 |

**MS spectrum**

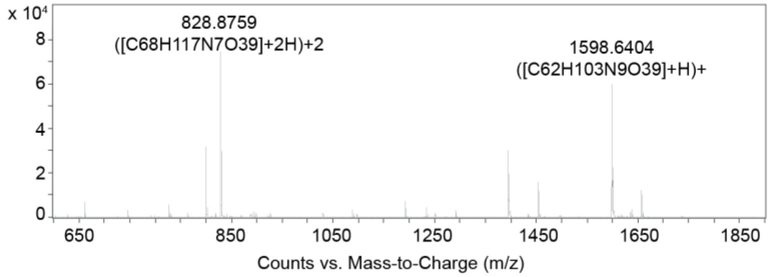

**Figure S5.** LC-MS results of modification of **4** with 8-azido-octanoic acid to give **6**. **a,** Scheme for preparing **6** by reaction of **4** with **5**. **b,** LC-MS result of compound **6**.

**a**

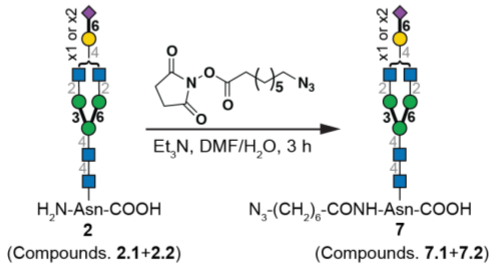

**b**

**Compound table**

| Compound Label | RT | Mass | Abund | Formula | Tgt Mass | Diff (ppm) |
|---|---|---|---|---|---|---|
| Compound7.1: C96H157N11O65 | 3.54 | 2503.9273 | 54234 | C96H157N11O65 | 2503.9318 | -1.8 |
| Compound7.2: C79H130N10O48 | 3.70 | 2050.7842 | 12104 | C79H130N10O52 | 2050.7836 | -0.34 |

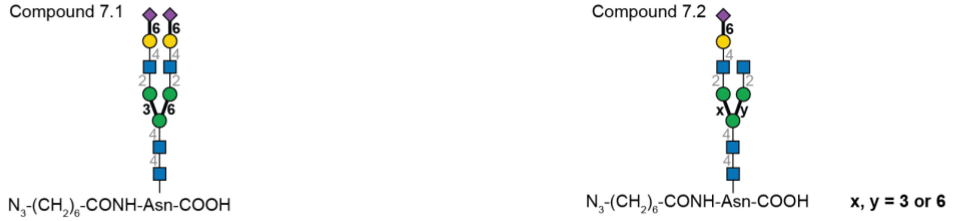

| Compound 7.1 | *m/z* | RT | Algorithm | Mass |
|---|---|---|---|---|
| $C_{96}H_{157}N_{11}O_{65}$ | 1253.4721 | 3.54 | Find by Formula | 2503.9273 |

| Compound 7.2 | *m/z* | RT | Algorithm | Mass |
|---|---|---|---|---|
| $C_{79}H_{130}N_{10}O_{52}$ | 1026.3994 | 3.70 | Find by Formula | 2050.7842 |

**Compound Chromatograms**

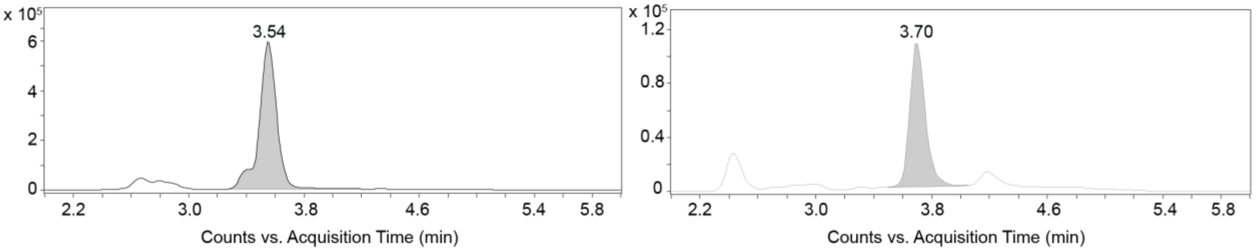

**Figure S6.** LC-MS results modification of **2** with 8-azido-octanoic acid to give **7**.
**a,** Scheme for preparing **7** via reaction of **2** with **5** (procedure analogous to **Fig. S5.**). **b,** LC-MS analysis of Compounds 7.1+7.2 showing 2 components with different retention time.

**a**

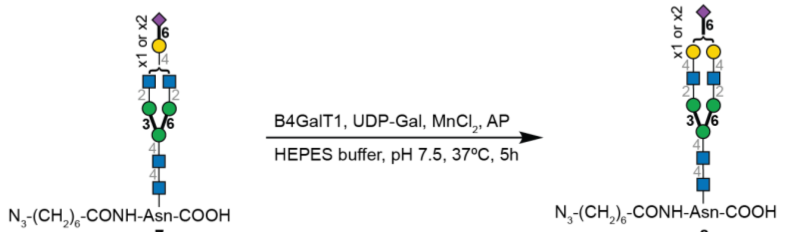

**7**

**8**

(Compounds. **8.1**+**8.2**)

**b**

**Compound table**

| Compound Label | RT | Mass | Abund | Formula | Tgt Mass | Diff (ppm) |
|---|---|---|---|---|---|---|
| Compound8.1: C96H157N11O65 | 5.94 | 2503.9295 | 4740 | C96H157N11O65 | 2503.9318 | -0.9 |
| Compound8.2: C85H140N10O57 | 5.98 | 2212.8358 | 1642 | C85H140N10O57 | 2212.8364 | -0.26 |

Compound 8.1

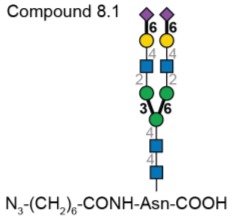

Compound 8.2

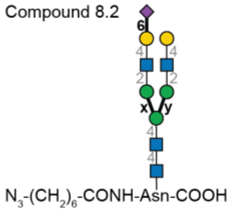

**x, y = 3 or 6**

| Compound 8.1 | *m/z* | RT | Algorithm | Mass |
|---|---|---|---|---|
| $C_{96}H_{157}N_{11}O_{65}$ | 835.9852 | 5.94 | Find by Formula | 2503.9295 |

| Compound 8.2 | *m/z* | RT | Algorithm | Mass |
|---|---|---|---|---|
| $C_{85}H_{140}N_{10}O_{57}$ | 1107.9275 | 5.98 | Find by Formula | 2212.8358 |

**Compound Chromatograms**

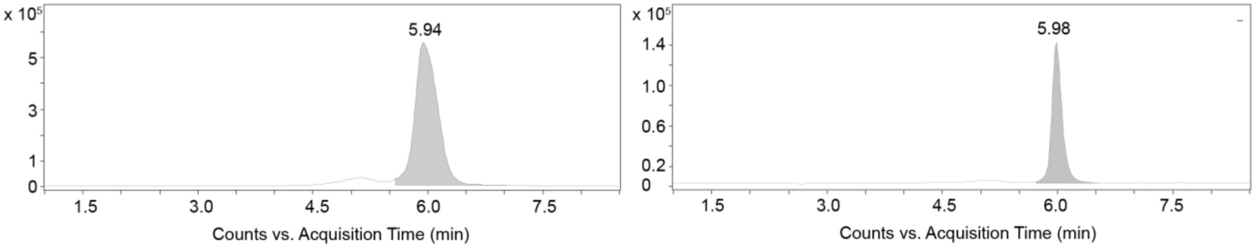

**Figure S7.** LC-MS results after B4GalT1-catalyzed reaction of mixtures **7**, giving **8**.
**a,** Scheme of B4GalT1-catalyzed reaction on mixtures **7** to generate desired structures of **8**. Enzymatic reaction in solution proved the possibility of installing galactose using heterogeneous mixtures **7** as starting material to afford mixtures **8** (Compounds 8.1+8.2). **b,** LC-MS analysis confirmed the completion of B4GalT1-catalyzed reaction to afford mixtures **8**.

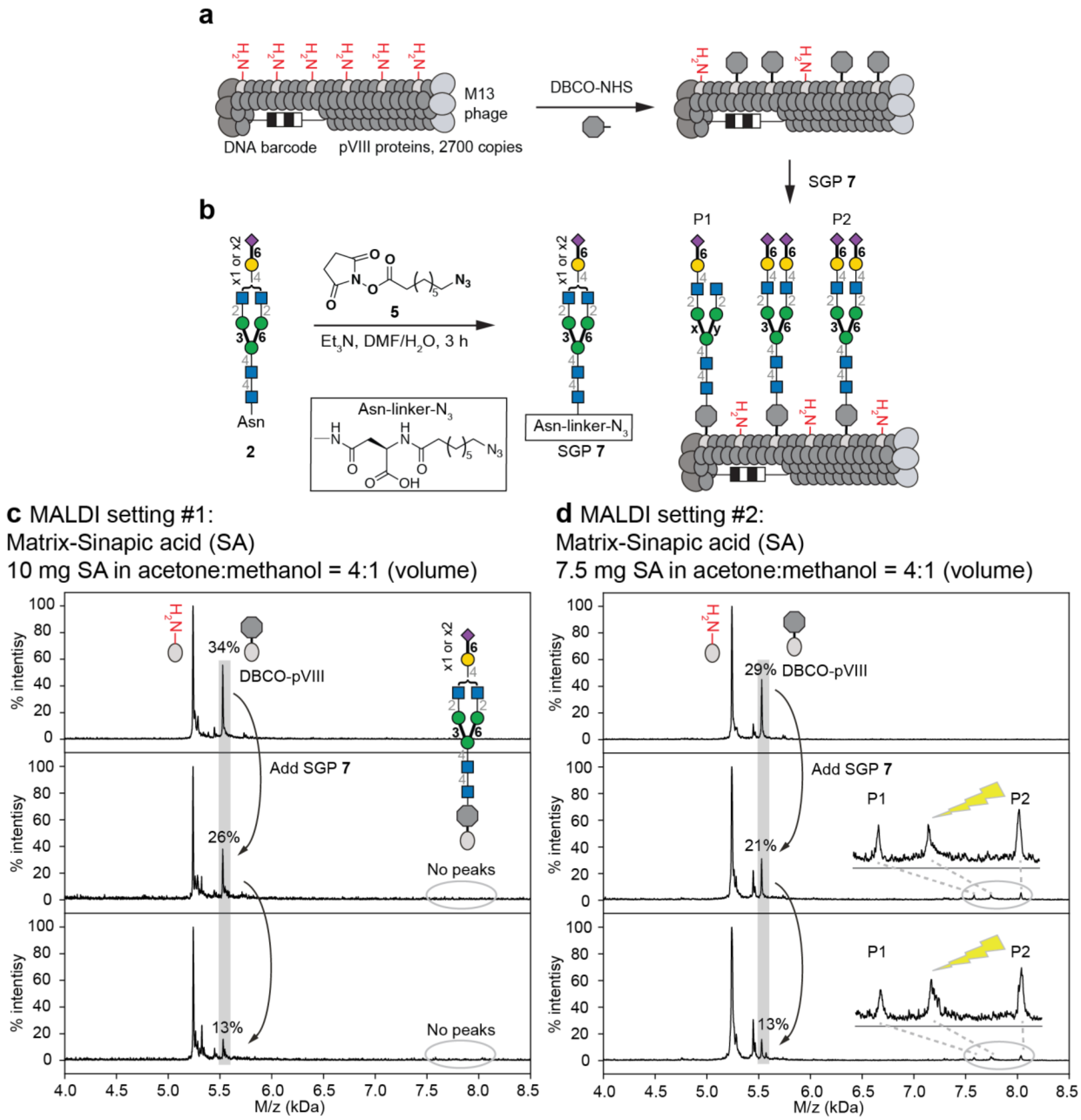

**Figure S8.** Optimization of MALDI-TOF of sialylated biantennary *N*-linked oligosaccharides **7** on phage.

**a,** Two steps chemical glycosylation of **7** on phage. **b,** Preparation of heterogeneous azido-*N*-glycan **7**. **c,** MALDI-TOF MS spectrometry characterized the decreasing intensity of alkyne-functionalized product (DBCO-pVIII) after incorporation of azido-modified biantennary *N*-linked oligosaccharides 7, which indicated the chemical glycosylation step did work; however glycosylated pVIII has not been observed under previously reported MALDI conditions[2]. The problem was solved by changing the MALDI matrix; minor attenuation in the amount of the sinapic acid made it possible to observe the glycosylated pVIII (**d**).

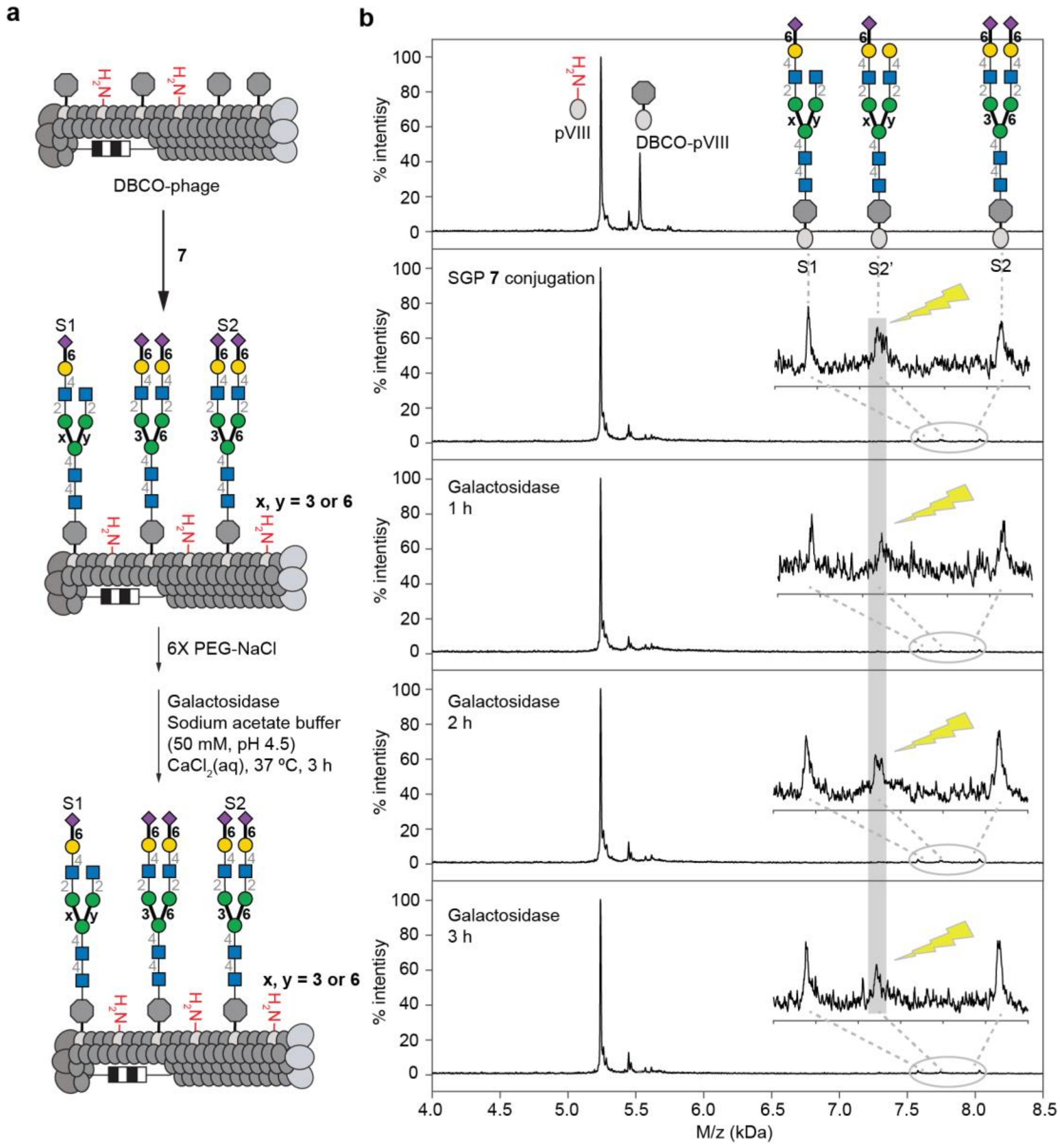

**Figure S9.** Direct *β*-galactosidase treatment of SGP **7** to confirm the corresponding peak S2' are indeed "ghost" species.

**a,** Scheme for SGP **7** conjugation followed by β-galactosidase digestion for 3 h. **b,** Time course results indicated the S2' peak is not responding to β-galactosidase treatment indicating that the species with terminal galactose are not present and they were generated due sialic acid cleavage during MALDI-TOF MS analysis.

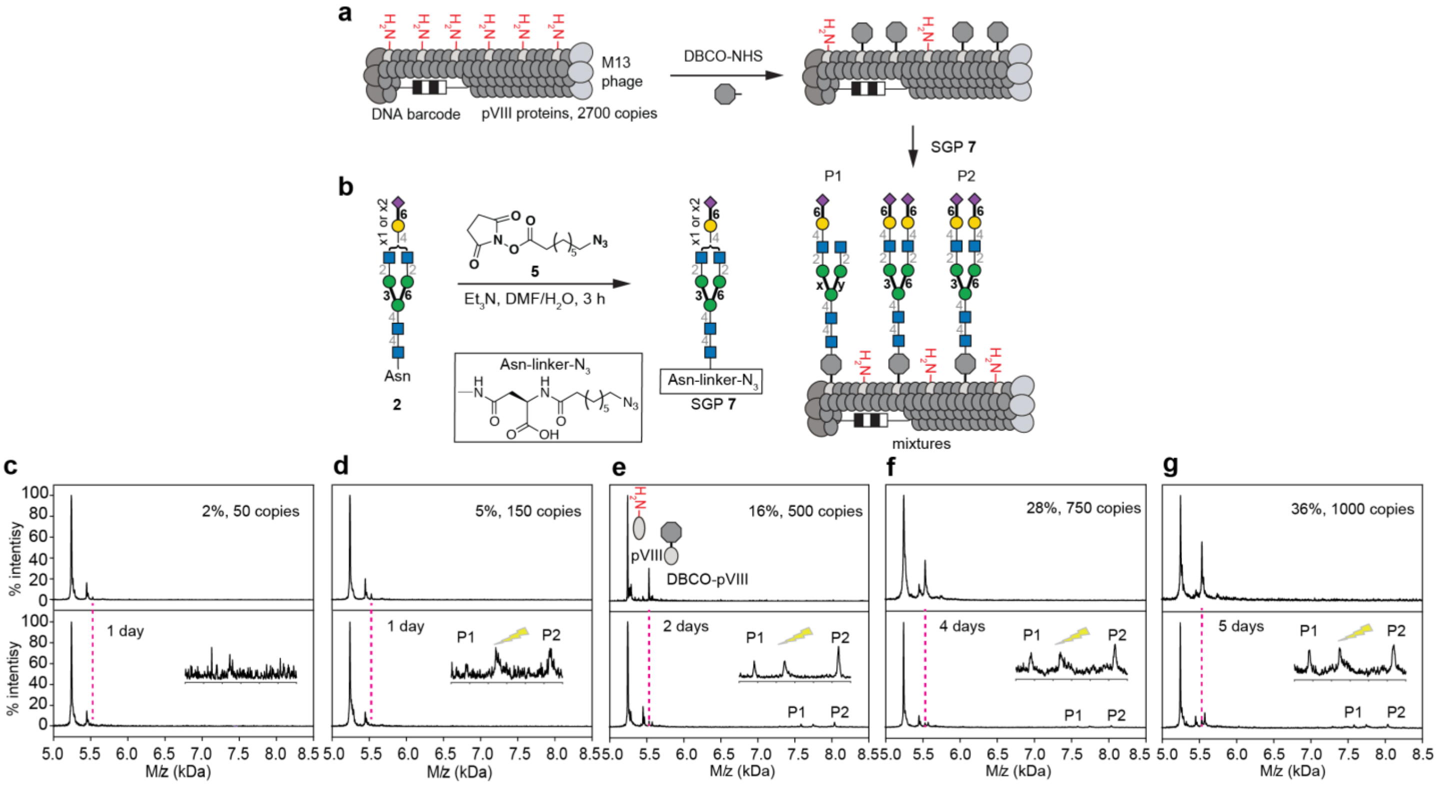

**Figure S10.** Conjugation of SGP glycan to phage at 5 different densities as monitored by MALDI TOF MS.

**a,** Phage was modified by 2-7 uL of 50 mM DBCO for 4 min-65 min at room temperature as detailed below to achieve the indicated copy number of DBCO molecules on phage surface. **b,** Modification of DBCO phage by azido-SGP (see Figure S6 for synthesis) resulted in complete (**c-f**) of nearly complete **g,** disappearance of DBCO-pVIII peak in MALDI indicating the completion of the reaction. Reaction was run for 1-5 days and higher DBCO density required longer completion times as indicated in (**c-g**). All conjugates were exposed to azido ethanol to cap any unreacted cyclooctyne residues.

Experimental details:

Combine clonal phage and DBCO in EpiTube according to desired density:
a) 200uL SDB + 2uL DBCO = 2% to 4% density (50 copies p8)
b) 200uL SDB + 2uL DBCO = 4% to 10% density (150 copies p8)
c) 200uL SDB + 4uL DBCO = 16% to 18% density (500 copies p8)
d) 200uL SDB + 4uL DBCO = 26% to 28% density (750 copies p8)
e) 200uL SDB + 7uL DBCO = 36% to 38% density (1000 copies p8)

Incubate at room temperature on shaker for time according to desired density:
a) 4 minutes = 2% to 4% density (50 copies p8)
b) 10 minutes = 4% to 10% density (150 copies p8)
c) 17.5 minutes = 16% to 18% density (500 copies p8)
d) 45 minutes = 26% to 28% density (750 copies p8)
e) 65 minutes = 36% to 38% density (1000 copies p8)

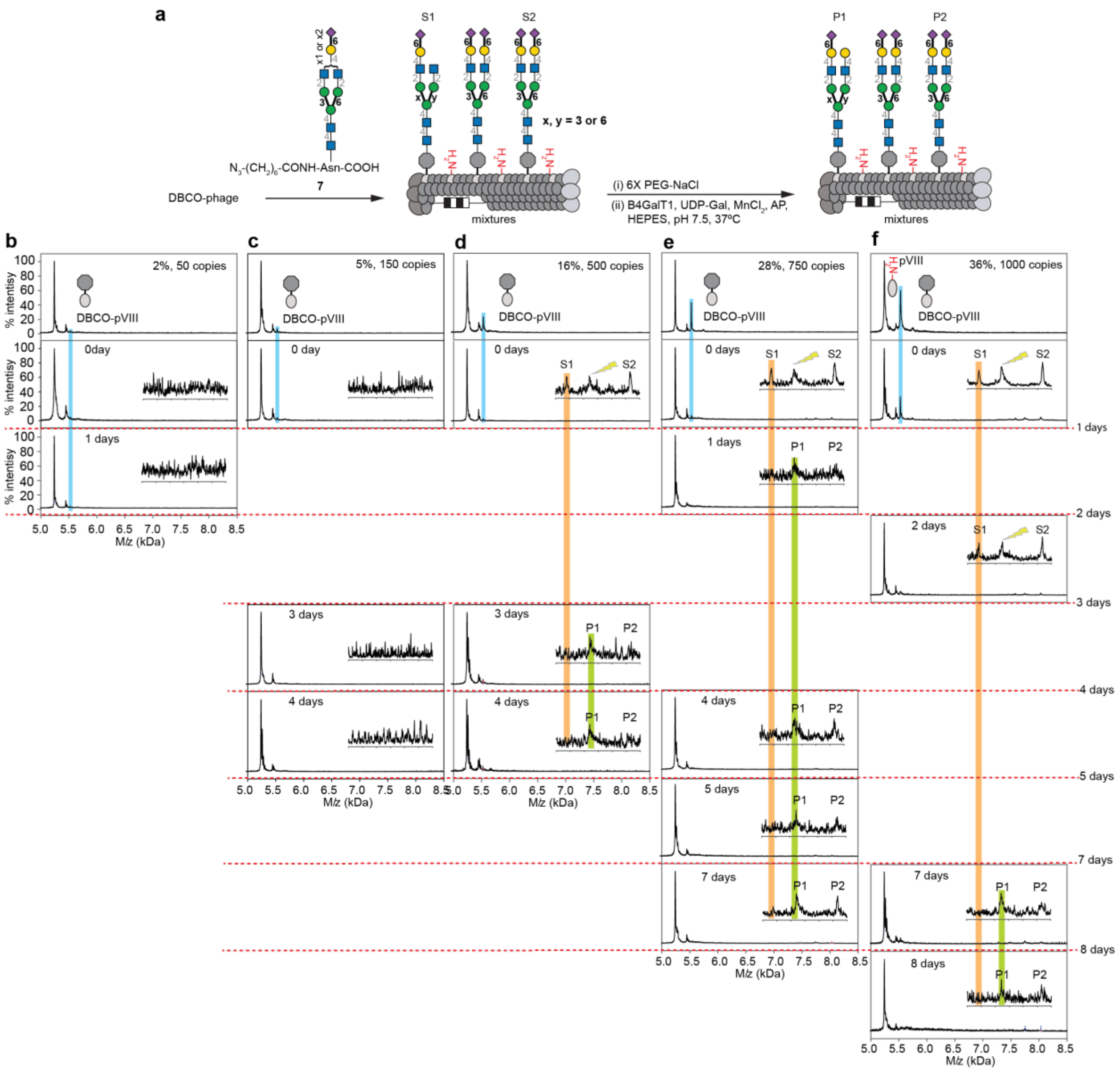

**Figure S11.** Enzymatic extension of SGP glycan (by B4GalT1) displayed on phage at 5 different densities monitored by MALDI TOF MS.

**a,** Phage was modified by 2-7 uL of 50 mM DBCO for 4 min-65 min at room temperature to achieve the indicated copy number of DBCO molecules on phage surface (see **Figure S10** for details). **b,** After modification of DBCO phage by azido-SGP, B4GalT1 was used to catalyze the transfer of galactose (Gal) from UDP-Gal. Reaction was run for 1-8 days and higher DBCO density required longer completion times as indicated in (**c-f**).

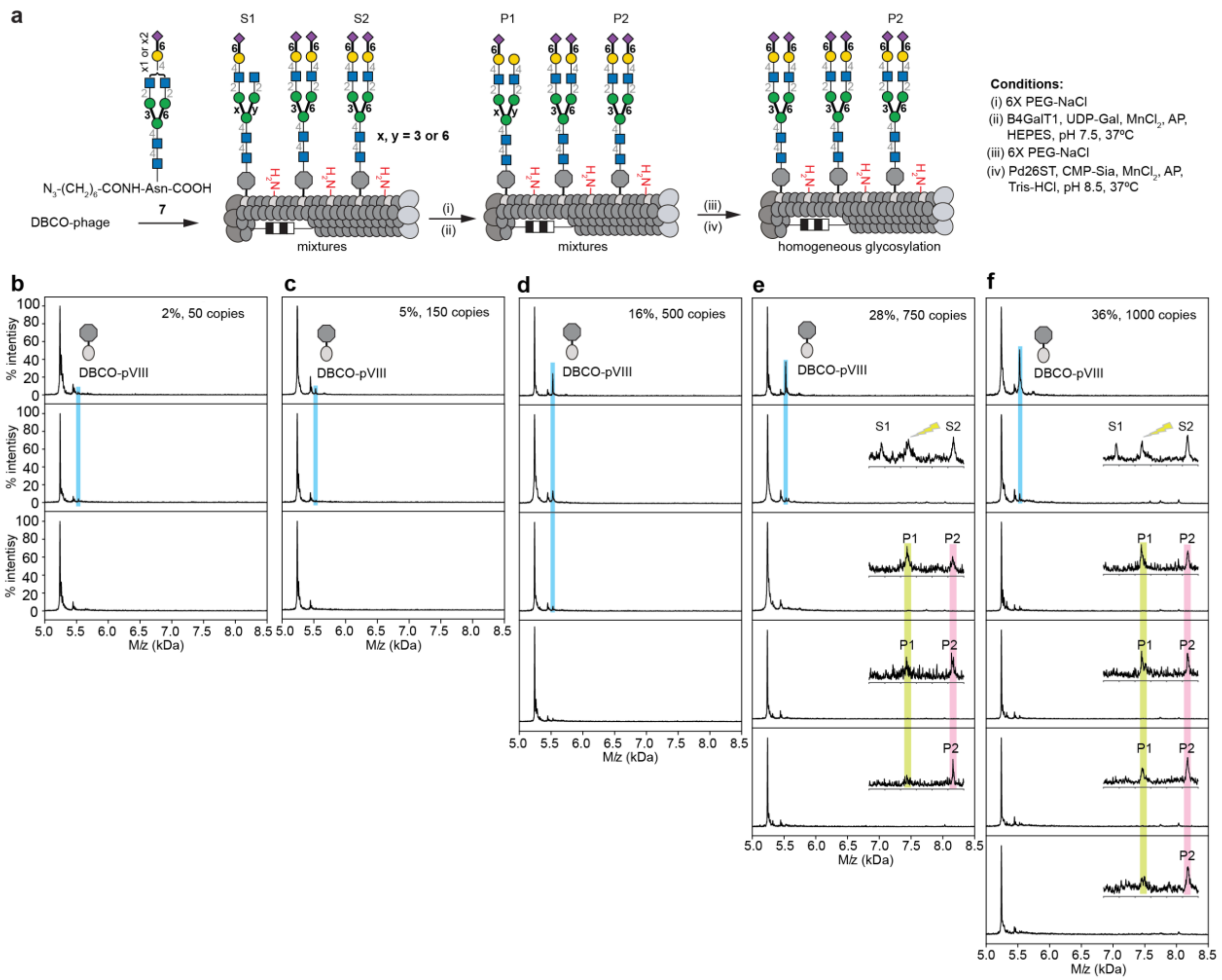

**Figure S12.** Two steps enzymatic extension (by B4GalT1 and Pd26ST) of SGP glycan displayed on phage at 5 different densities monitored by MALDI TOF MS.

**a,** Phage was modified by 2-7 uL of 50 mM DBCO for 4 min-65 min at room temperature to achieve the indicated copy number of DBCO molecules on phage surface (see **Figure S10** for details). **b,** After modification of DBCO phage by azido-SGP, β-1,4-galactosylation by B4GalT1 and α-2,6-linked sialylation by Pd26ST afforded homogeneous sialylated product P2. Reaction was run for 1-8 days; higher DBCO density required longer completion times as indicated in (**c-f**).

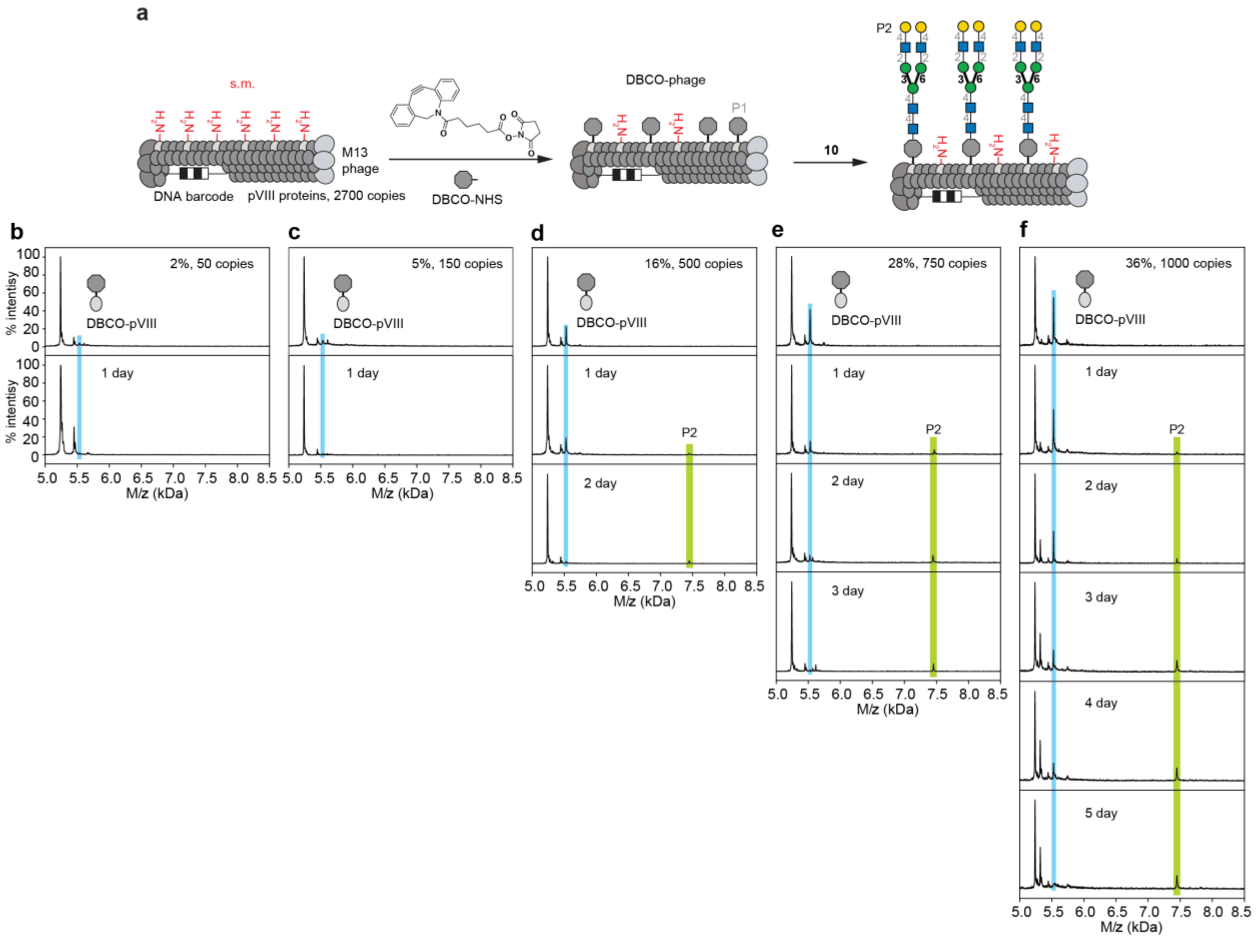

**Figure S13.** Conjugation of *N*-glycan **10** to phage at 5 different densities as monitored by MALDI TOF MS.

**a,** Phage was modified by 2-7 uL of 50 mM DBCO for 4 min-65 min at room temperature to achieve the indicated copy number of DBCO molecules on phage surface. **b,** Modification of DBCO phage by *N*-glycan **10** resulted in complete (**b-f**) disappearance of DBCO-pVIII peak in MALDI indicating the completion of the reaction. Reaction was run for 1-5 days; higher DBCO density required longer completion times as indicated in (**b-f**). All conjugates were exposed to azido ethanol to cap any unreacted cyclooctyne residues.

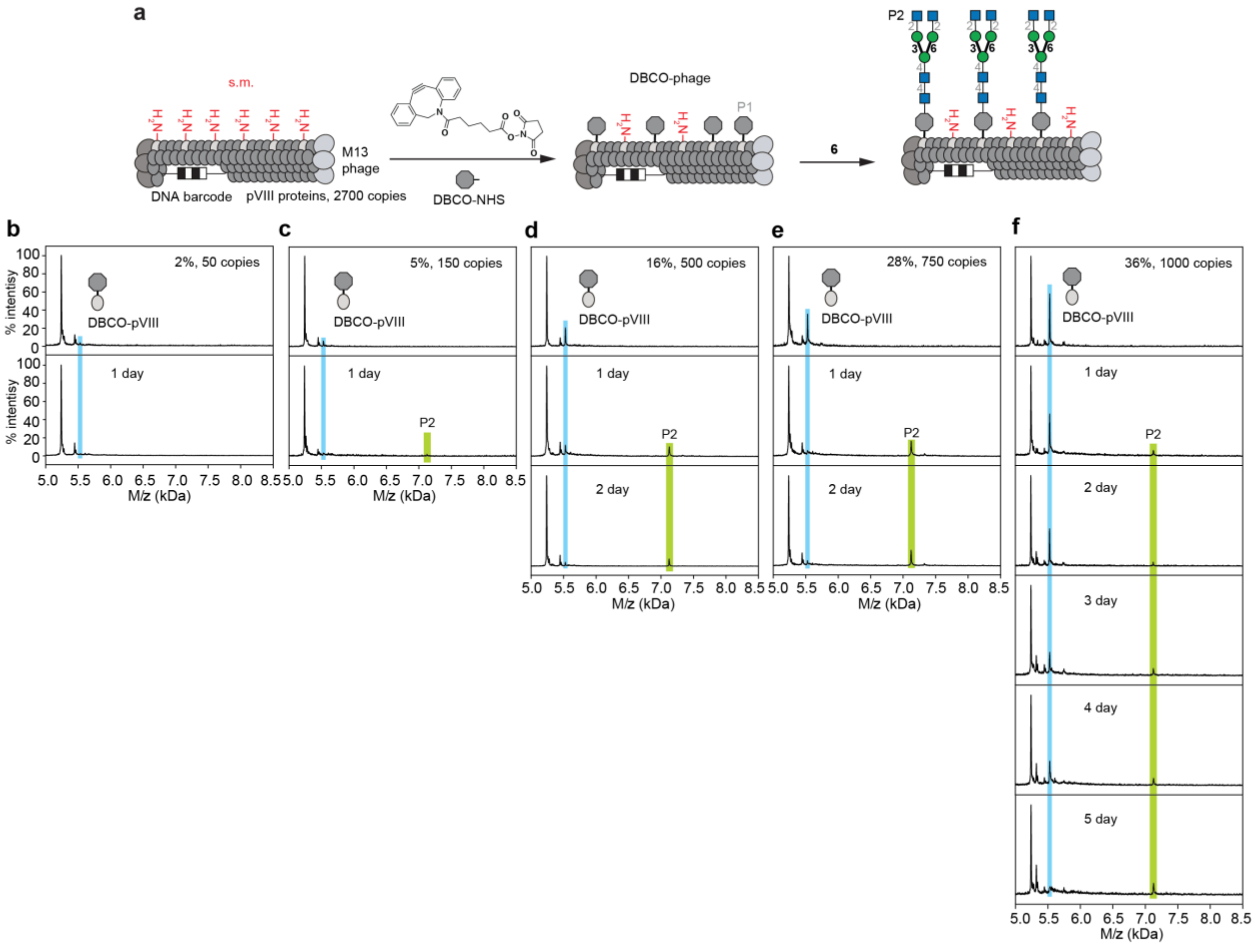

**Figure S14.** Conjugation of *N*-glycan **6** to phage at 5 different densities as monitored by MALDI TOF MS.

**a,** Phage was modified by 2-7 uL of 50 mM DBCO for 4 min-65 min at room temperature to achieve the indicated copy number of DBCO molecules on phage surface. **b,** Modification of DBCO phage by N-glycan **6** resulted in complete (**b-f**) disappearance of DBCO-pVIII peak in MALDI indicating the completion of the reaction. Reaction was run for 1-5 days; higher DBCO density required longer completion times as indicated in (**b-f**). All conjugates were exposed to azido ethanol to cap any unreacted cyclooctyne residues.

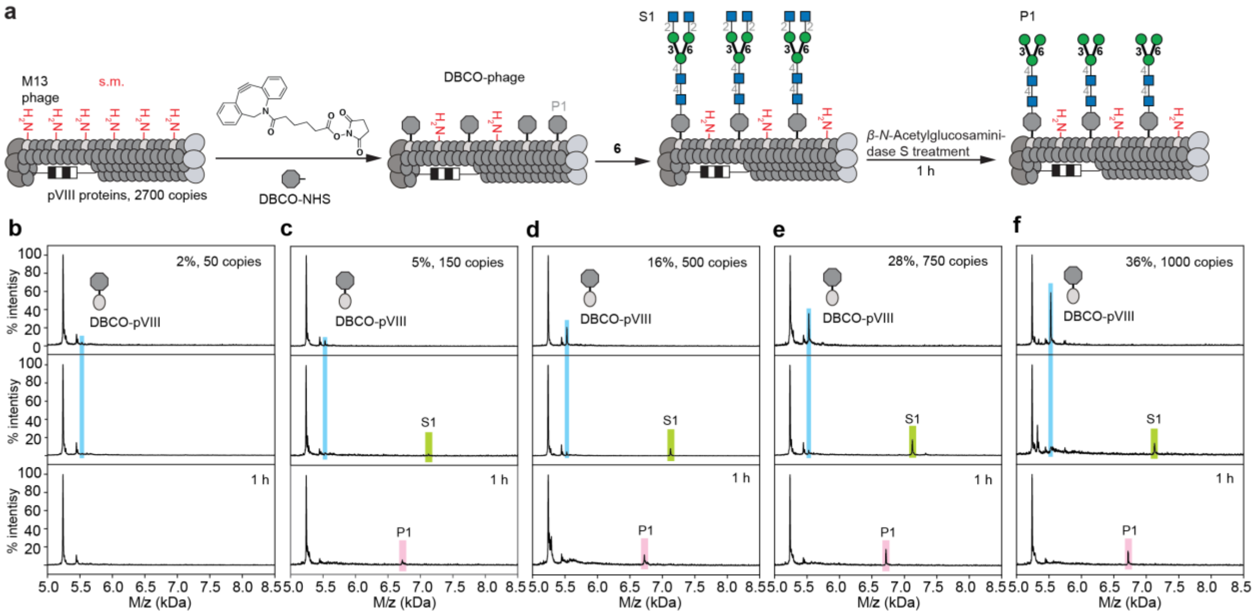

**Figure S15.** On-phage enzymatic trimming of **6** to generate terminal mannose *N*-glycan ($Man_3GlcNAc_2$) at 5 different densities as monitored by MALDI TOF MS.

**a,** Phage was modified by 2-7 uL of 50 mM DBCO for 4 min-65 min at room temperature to achieve the indicated copy number of DBCO molecules on phage surface. **b-f,** After modification of DBCO phage by S1 glycan, β-N-Acetylglucosaminidase S treatment of S1 provided desired P1 on pVIII. Each reaction was complete in 1 h.

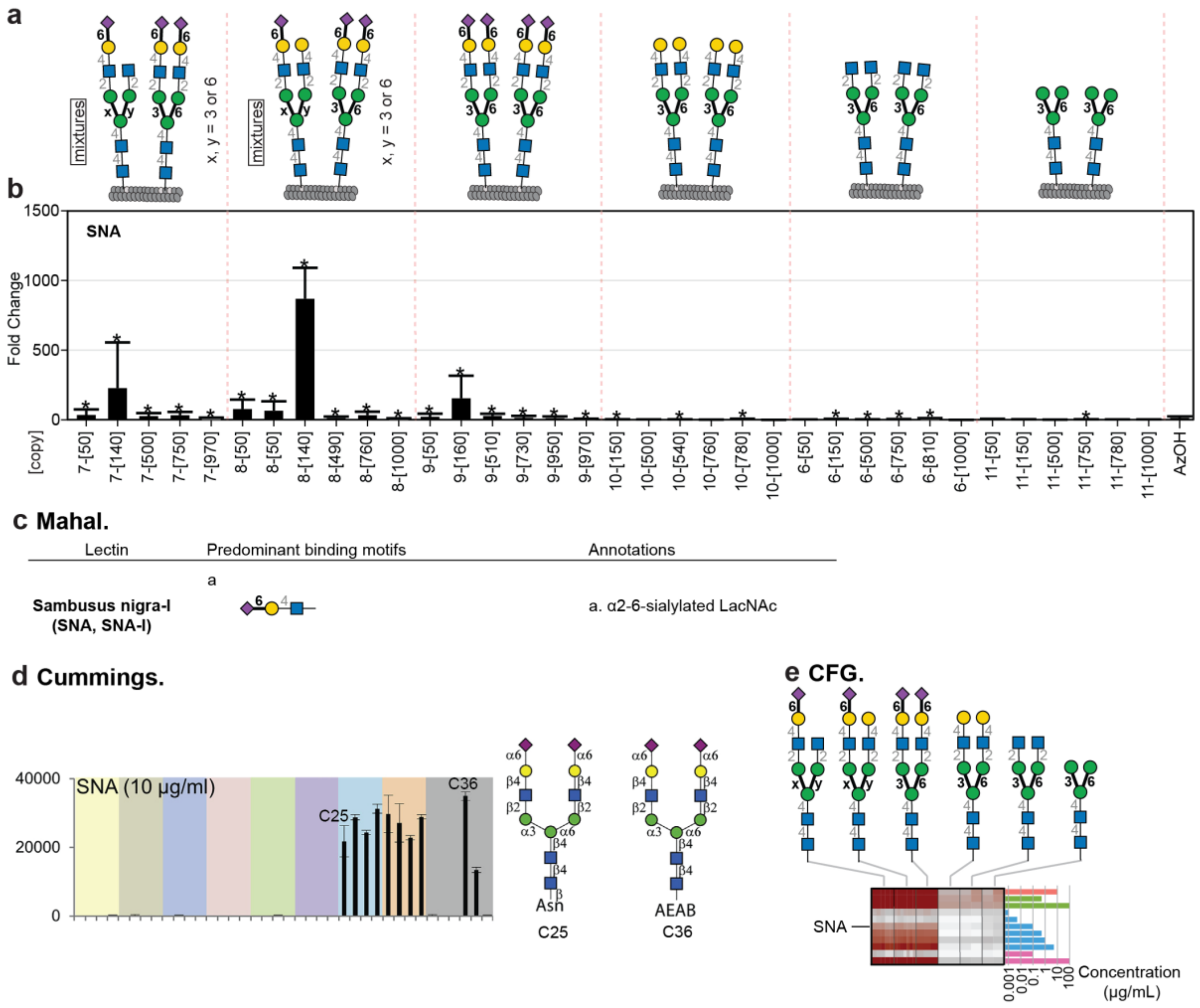

**Figure S16.** Binding of LiGA6×5 library to SNA-I.

**a,** LiGA6×5 components, same as **Figure 5**. **b,** Binding to SNA-I was calculated as fold change (FC) differential enrichment (DE) of each glycophage clone in SNA-I coated wells when compared to BSA coated wells. *denotes significantly enriched reads in DE analysis with FDR≤0.05 (n = 5 independent binding experiments). Error bars represent standard deviation propagated from the variance of the Trimmed Mean of M-values (TMM)-normalized sequencing data. **c,** Expected SNA-I binding specificities as described by Mahal and co-workers[10]. **d,** SNA-I binding to glass-based N-glycan array measured by Cummings and co-workers[11]. **e,** SNA-I binding to N-glycans on glass-based array produced by Consortium of Functional Glycomics[10].

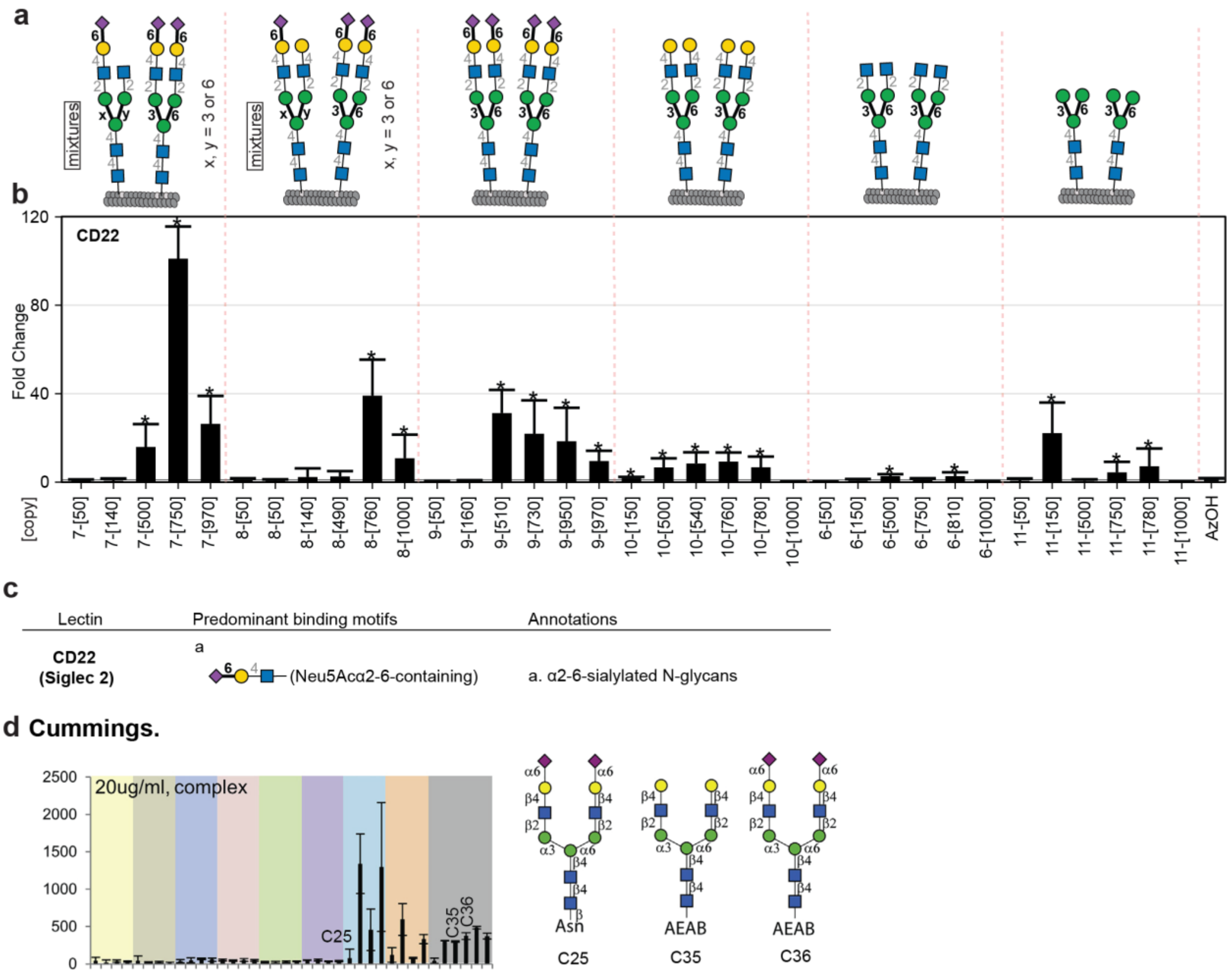

**Figure S17.** Binding of LiGA6×5 library to CD22 (Siglec-2).

**a,** The component of LiGA6×5, same as **Figure 5**. **b,** Binding to CD22 was calculated as fold change (FC) differential enrichment (DE) of each glycophage clone in CD22 coated wells when compared to BSA coated wells. * denote significantly enriched reads in DE analysis with FDR≤0.05 (n=5 independent binding experiments). Error bars represent s.d. propagated from the variance of the TMM-normalized sequencing data. **c,** Expected CD22 binding specificities as described by Cummings and co-workers[11]. **d,** CD22 binding to glass-based N-glycan array measured by Cummings and co-workers[11].

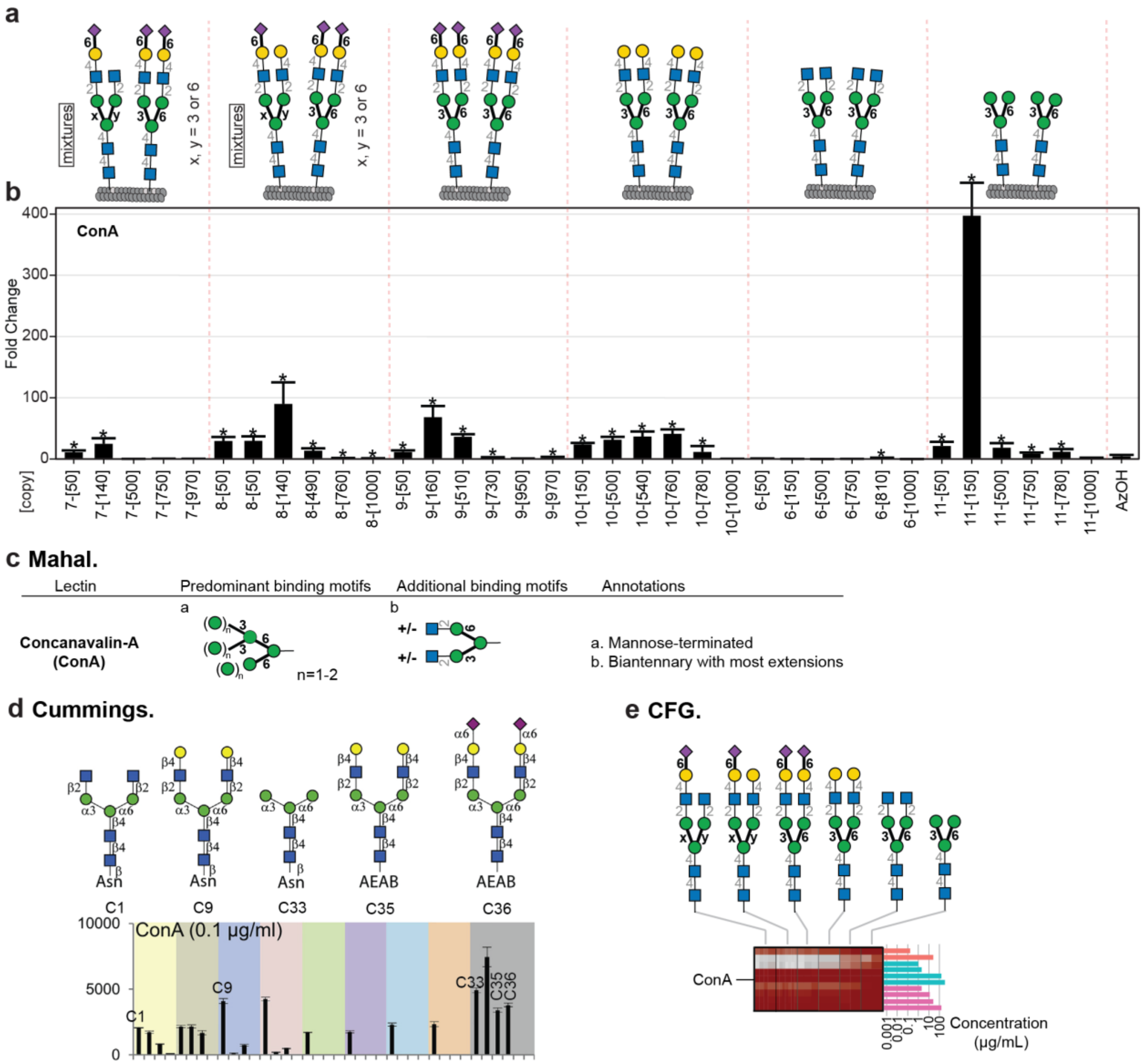

**Figure S18.** Binding of LiGA6×5 library to ConA.

**a,** The component of LiGA6×5, same as **Figure 5**. **b,** Binding to ConA was calculated as FC of each glycophage clone in ConA coated wells when compared to BSA coated wells. * denote significantly enriched reads (FDR≤0.05, n=5 independent binding experiments). Error bars represent s.d. propagated from the variance of the TMM-normalized sequencing data. **c,** Expected ConA binding specificities as described by Mahal and co-workers[10]. **d,** ConA binding to glass-based N-glycan array measured by Cummings and co-workers[11]. **e,** ConA binding to N-glycans on glass-based array produced by Consortium of Functional Glycomics[10].

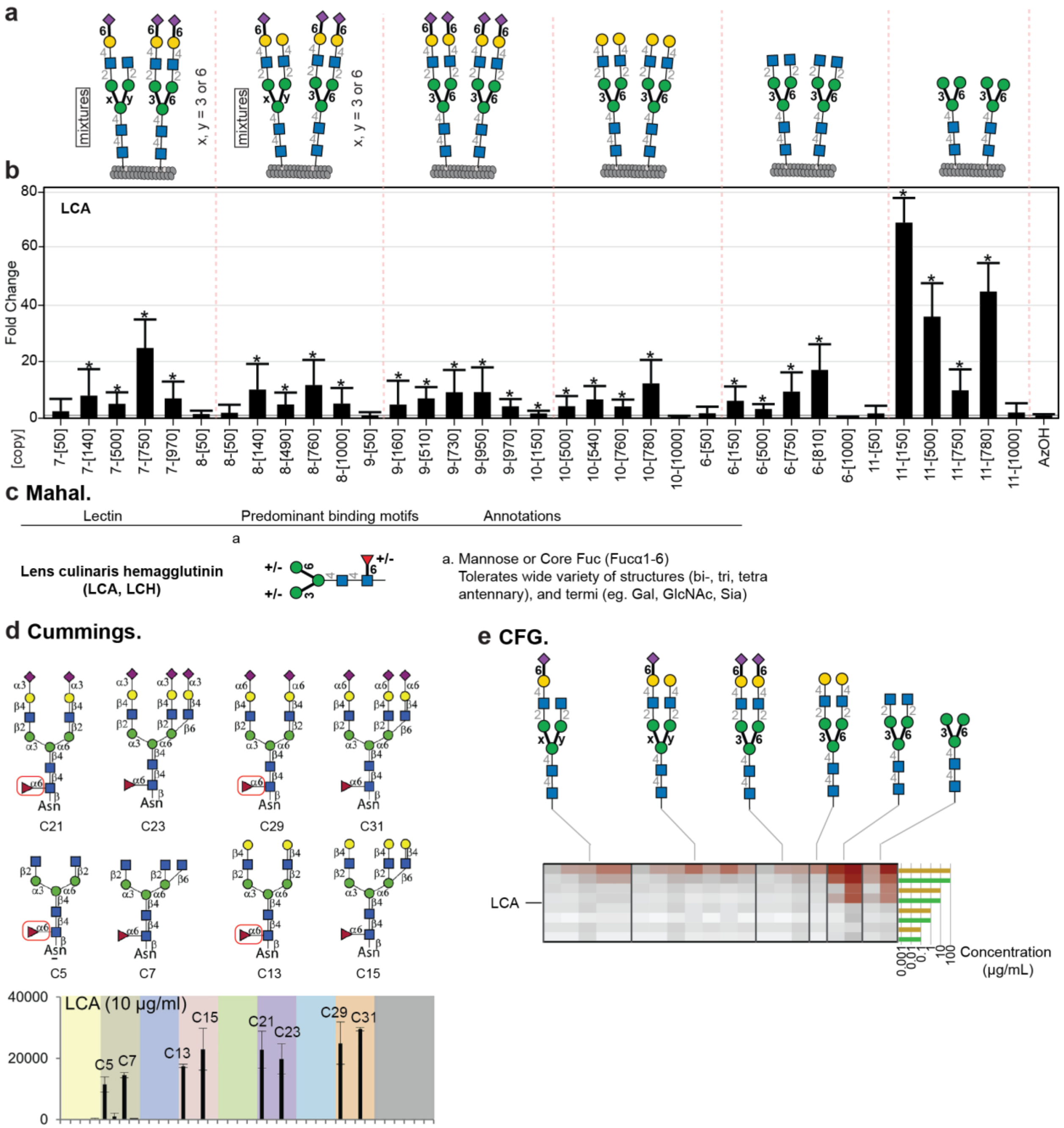

**Figure S19.** Binding of LiGA6×5 library to LCA.

**a,** The component of LiGA6×5, same as **Figure 5**. **b,** Binding to LCA was calculated as FC of each glycophage clone in LCA coated wells when compared to BSA coated wells. * denote significantly enriched reads with FDR≤0.05. (n=5 independent binding experiments). Error bars represent s.d. propagated from the variance of the TMM-normalized sequencing data. **c,** Expected LCA binding specificities as described by Mahal and co-workers[10]. **d,** LCA binding to glass-based N-glycan array measured by Cummings and co-workers[11]. **e,** LCA binding to N-glycans on glass-based array produced by Consortium of Functional Glycomics[10].

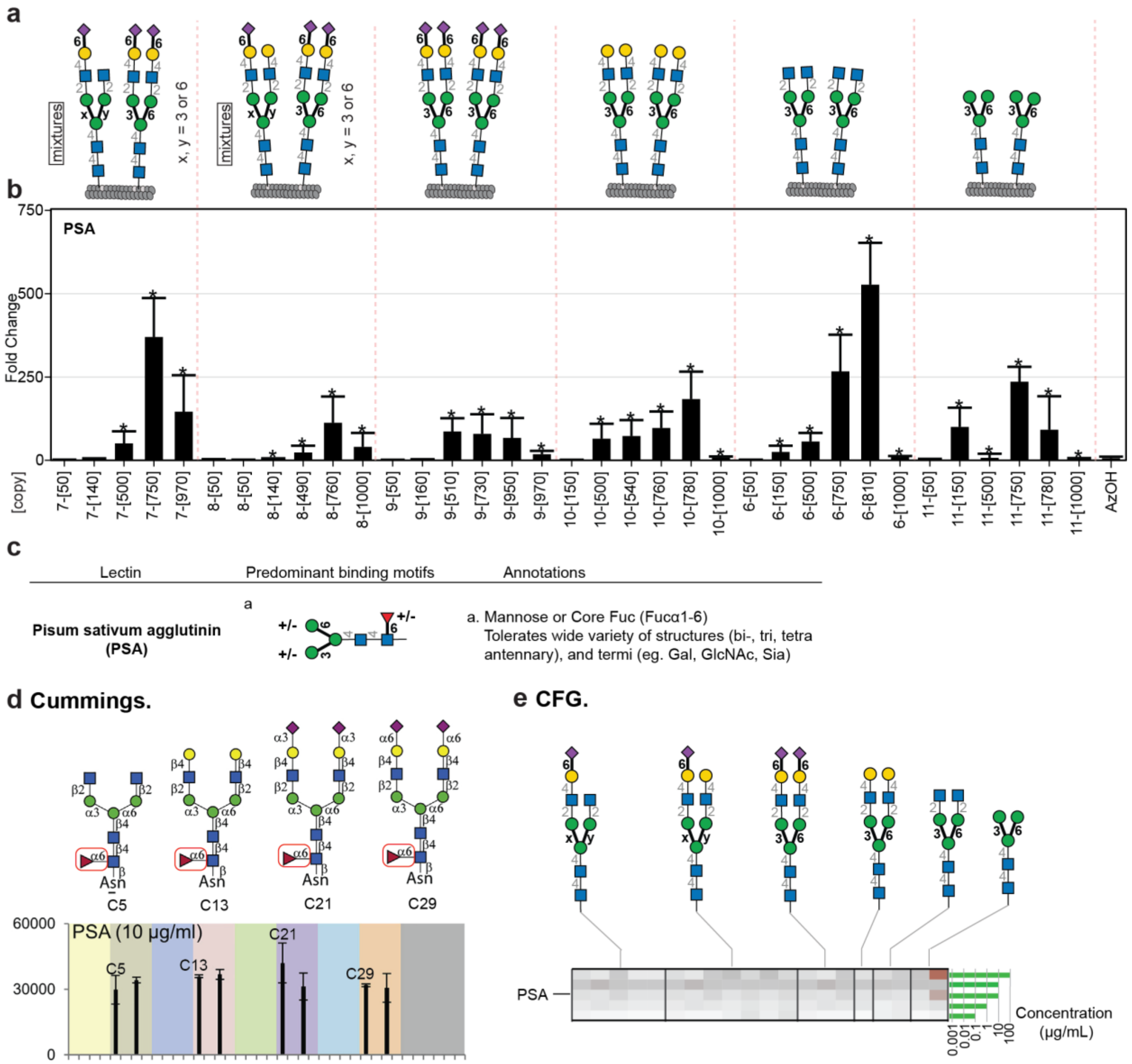

**Figure S20.** Binding of LiGA6×5 library to PSA.
**a,** The component of LiGA6×5, same as **Figure S5**. **b,** Binding to PSA was calculated as FC of each glycophage clone in PSA coated wells when compared to BSA coated wells. * denote significantly enriched reads with FDR≤0.05 (n=5 independent binding experiments). Error bars represent s.d. propagated from the variance of the TMM-normalized sequencing data. **c,** Expected PSA binding specificities as described by Mahal and co-workers[10]. **d,** PSA binding to glass-based N-glycan array measured by Cummings and co-workers[11]. **e,** PSA binding to N-glycans on glass-based array produced by Consortium of Functional Glycomics[10].

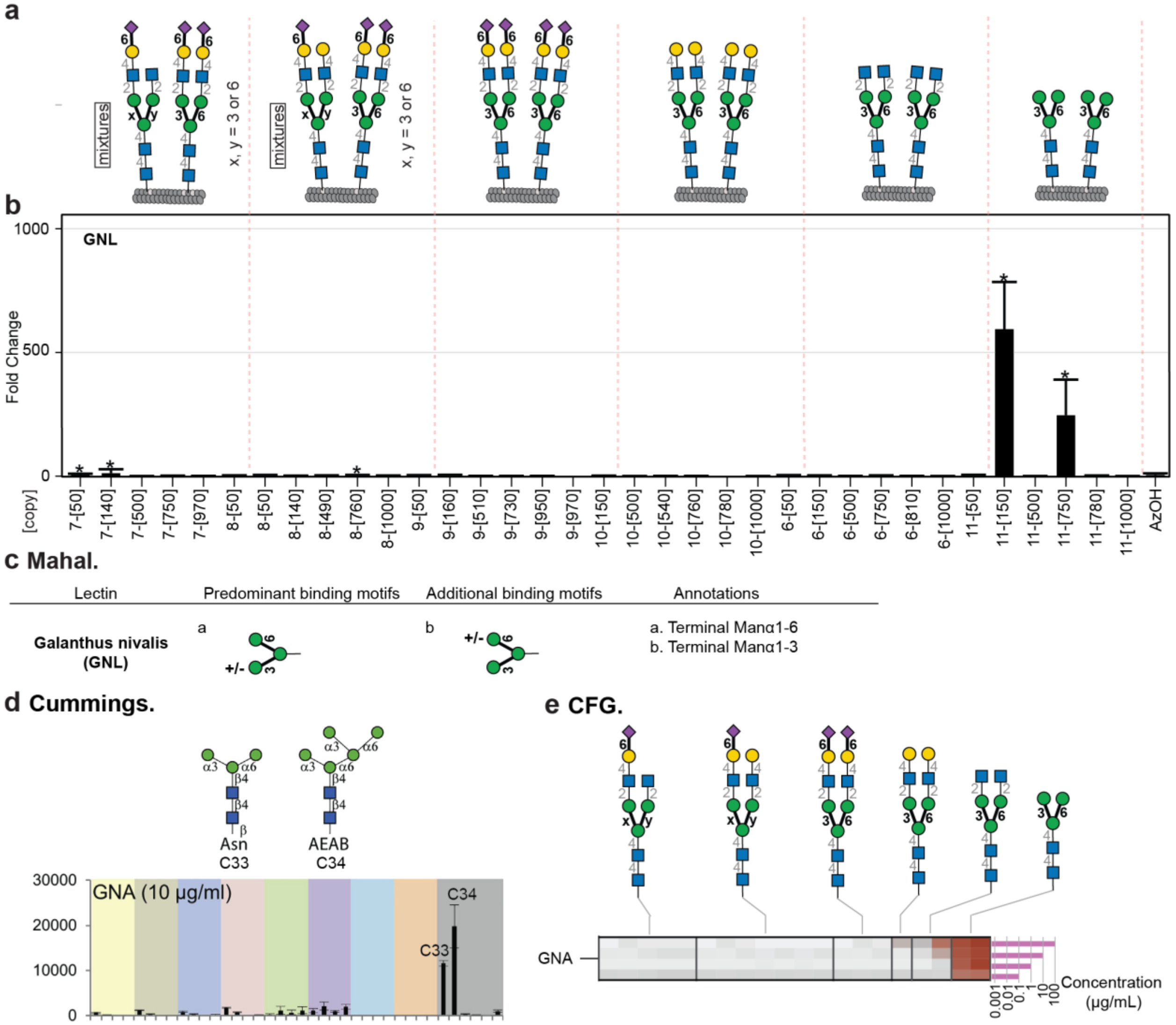

**Figure S21.** Binding of LiGA6×5 library to GNL (GNA).

**a,** The component of LiGA6×5, same as **Figure S5**. **b,** Binding to GNL was calculated as FC of each glycophage clone in GNL coated wells when compared to BSA coated wells. * denote significantly enriched reads with FDR≤0.05 (n=5 independent binding experiments). Error bars represent s.d. propagated from the variance of the TMM-normalized sequencing data. **c,** Expected GNL binding specificities as described by Mahal and co-workers[10]. **d,** GNL binding to glass-based N-glycan array measured by Cummings and co-workers[11]. **e,** GNL binding to N-glycans on glass-based array produced by Consortium of Functional Glycomics[10].

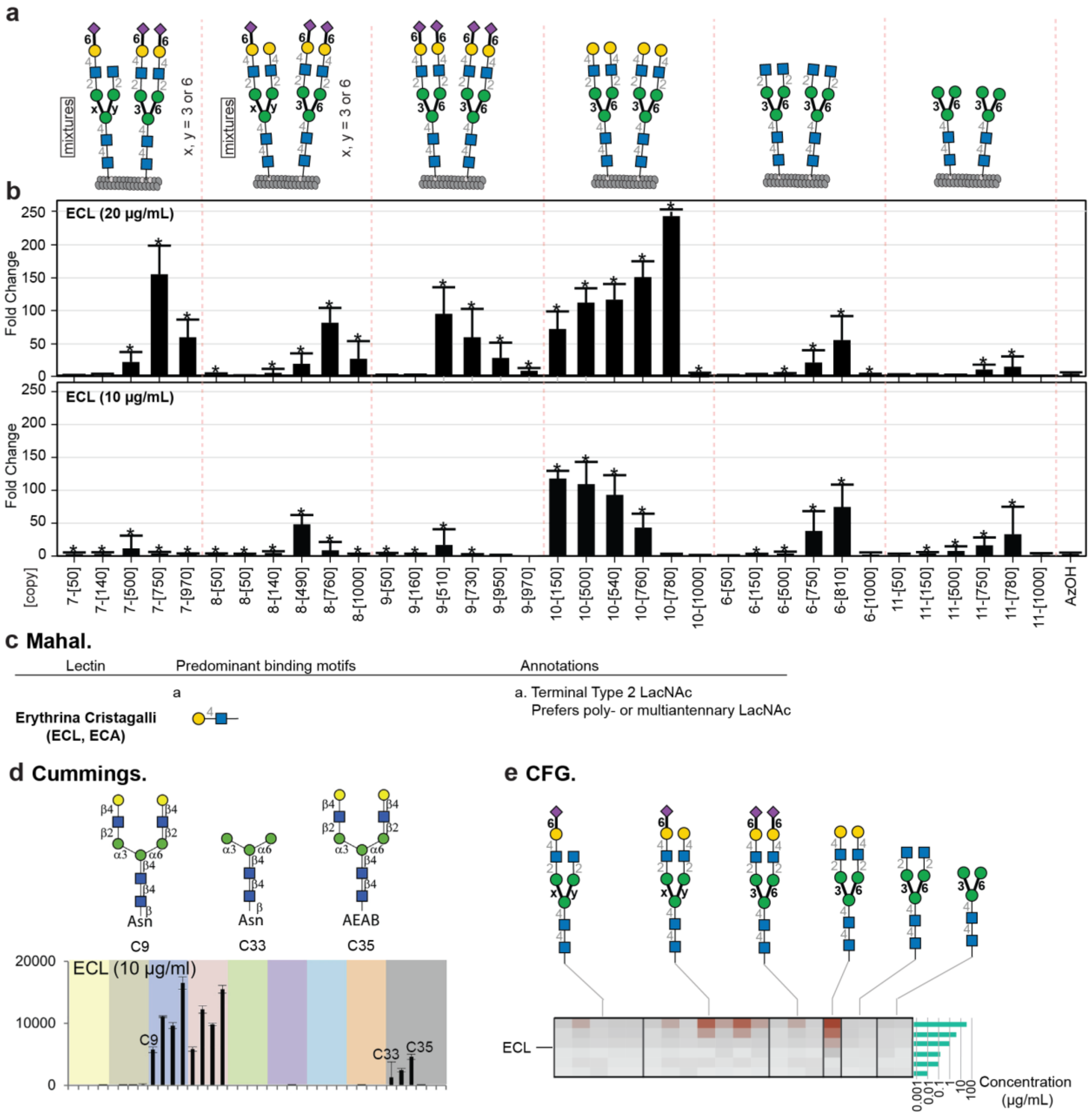

**Figure S22.** Binding of LiGA6×5 library to ECL.

**a,** The component of LiGA6×5, same as **Figure S5**. **b,** Binding to ECL was calculated as fold change (FC) differential enrichment (DE) of each glycophage clone in ECL coated wells when compared to BSA coated wells. * denote significantly enriched reads with FDR≤0.05 (n=5 independent binding experiments). Error bars represent s.d. propagated from the variance of the TMM-normalized sequencing data. **c,** Expected ECL binding specificities as described by Mahal and co-workers[10]. **d,** ECL binding to glass-based N-glycan array measured by Cummings and co-workers[11]. **e,** ECL binding to N-glycans on glass-based array produced by Consortium of Functional Glycomics[10].

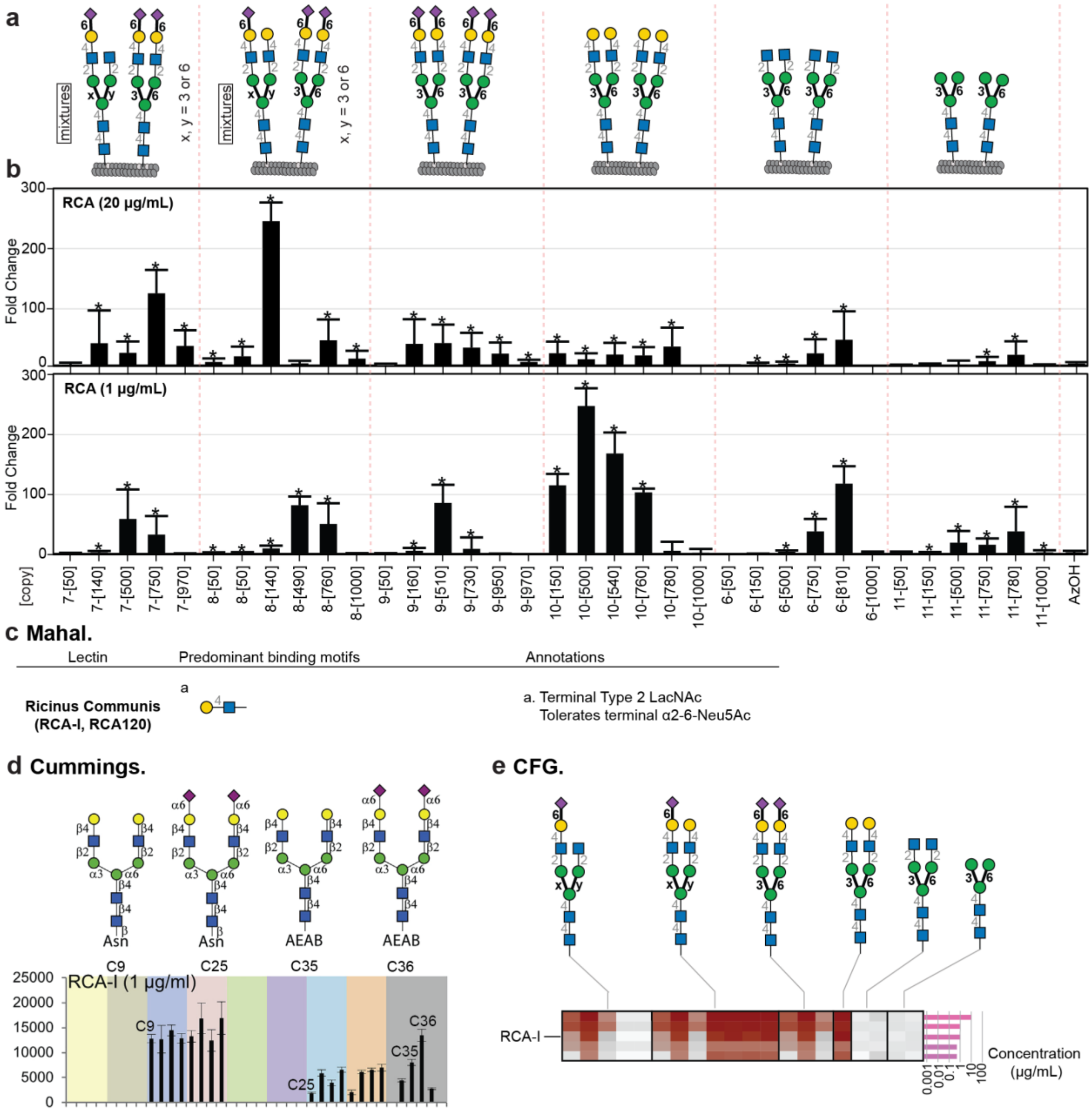

**Figure S23.** Binding of LiGA6×5 library to RCA-I.

**a,** The component of LiGA6×5, same as **Figure S5**. **b,** Binding to RCA-I was calculated as fold change (FC) differential enrichment (DE) of each glycophage clone in RCA-I coated wells when compared to BSA coated wells. * denote significantly enriched (black) or depleted (pink) reads in DE analysis with FDR≤0.05 (n=5 independent binding experiments). Error bars represent s.d. propagated from the variance of the TMM-normalized sequencing data. **c,** Expected RCA-I binding specificities as described by Mahal and co-workers[10]. **d,** RCA-I binding to glass-based N-glycan array measured by Cummings and co-workers[11]. **e,** RCA-I binding to N-glycans on glass-based array produced by Consortium of Functional Glycomics[10].

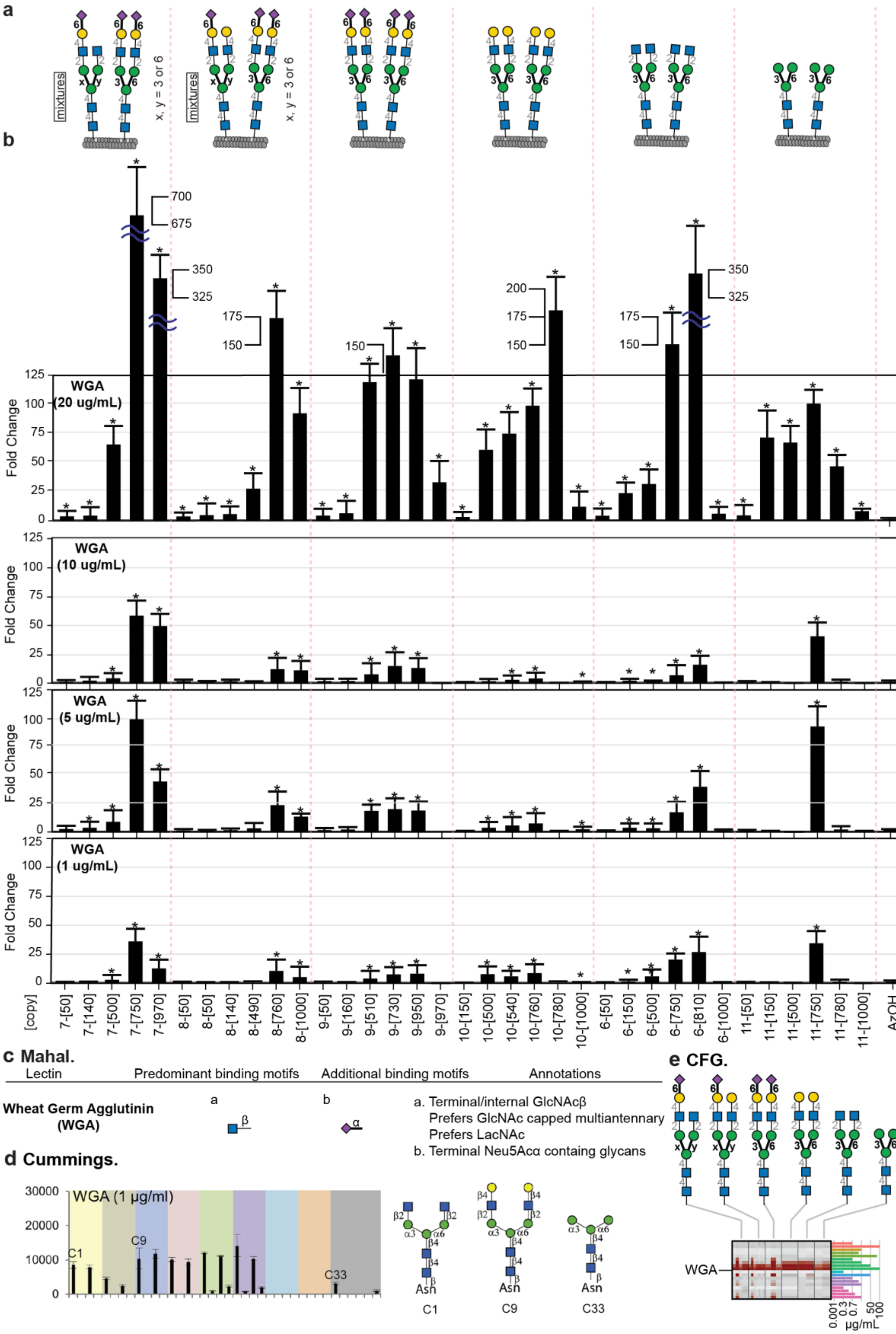

**(legend on the next page)**

**Figure S24.** Binding of LiGA6×5 library to WGA.

**a,** The component of LiGA6×5, same as **Figure S5**. **b,** Binding to WGA was calculated as FC of each glycophage clone in WGA coated wells when compared to BSA coated wells. * denote significantly enriched reads with FDR≤0.05 (n=5 independent binding experiments). Error bars represent s.d. propagated from the variance of the TMM-normalized sequencing data. **c,** Expected WGA binding specificities as described by Mahal and co-workers[10]. **d,** WGA binding to glass-based N-glycan array measured by Cummings and co-workers[11]. **e,** WGA binding to N-glycans on glass-based array produced by Consortium of Functional Glycomics[10].

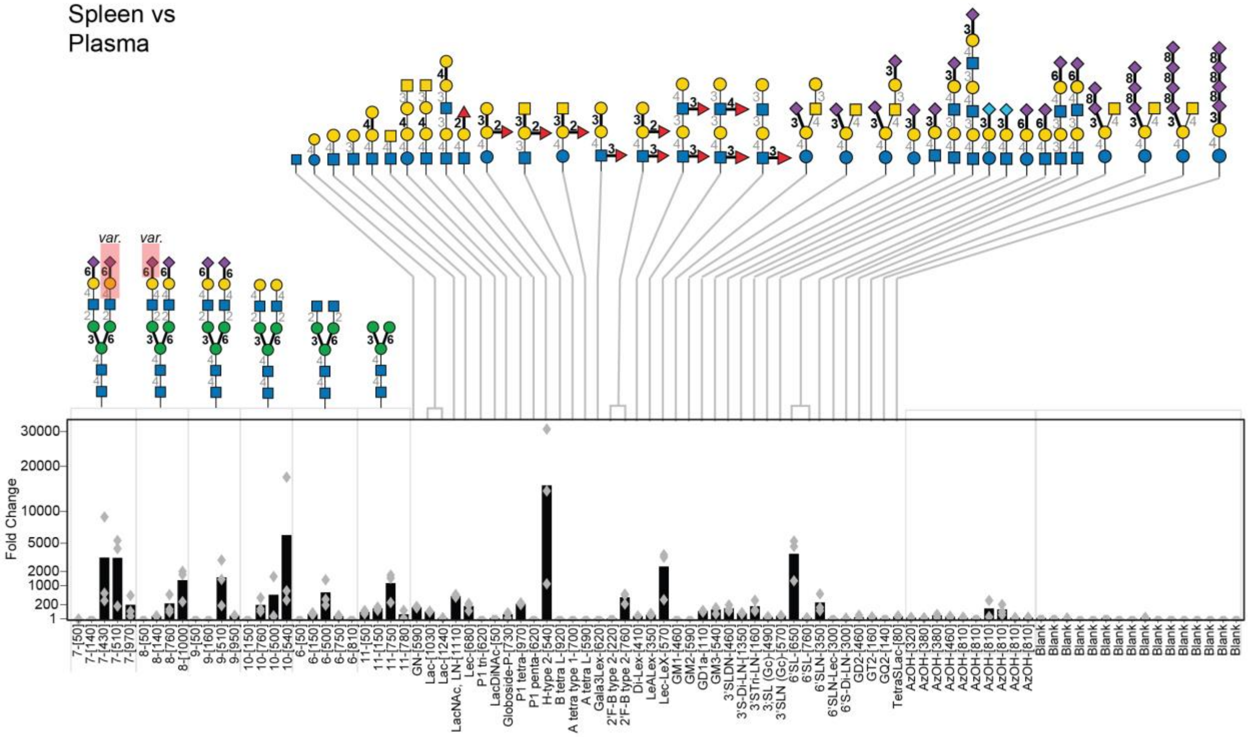

**Figure S25.** Summary of LiGA interaction with spleen *in-vivo* described as fold change (FC) enrichment in spleen with respect to plasma from the same animal.

The injection and processing of LiGA is described in Figure 6 (main text); the N-glycan and blank (unmodified) phage data is mirrored from Figure 6b for consistency and replotted as bar chart describing an average FC from n=3 mice and an overlaid scatter plot describing an FC from each mouse.

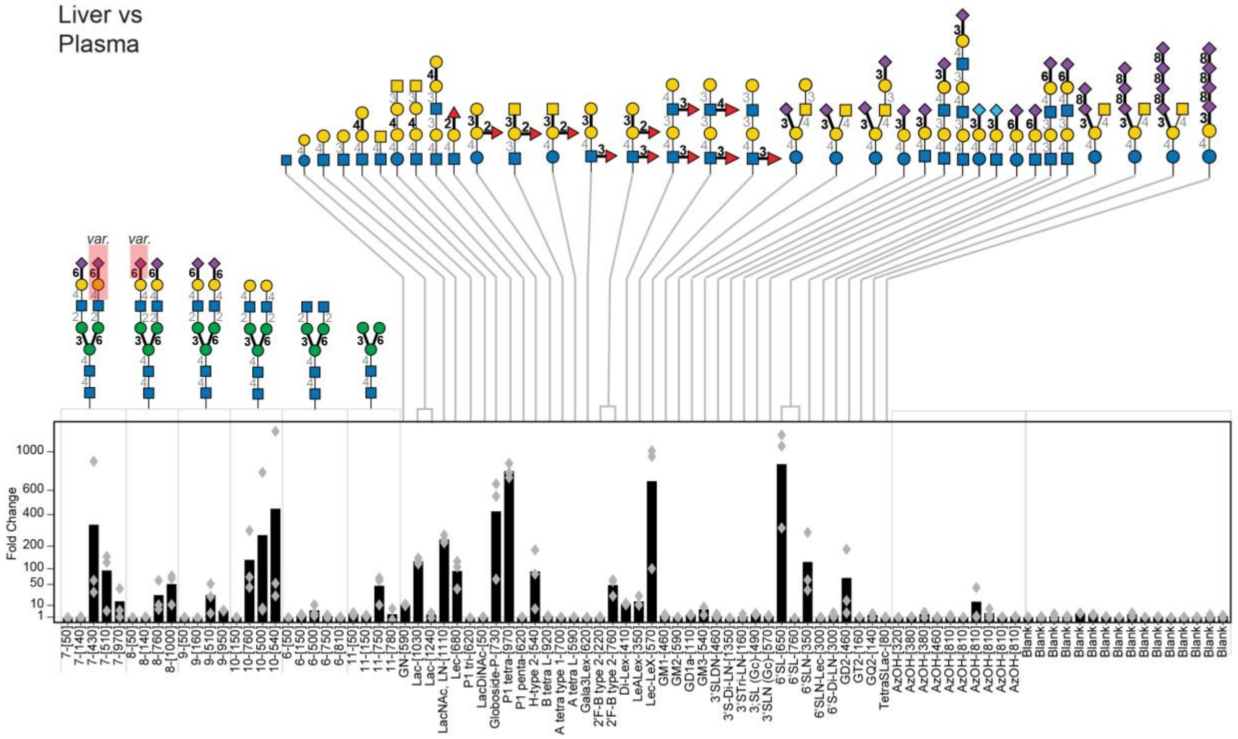

**Figure S26.** Summary of LiGA interaction with liver *in-vivo* described as fold change (FC) enrichment in liver with respect to plasma from the same animal.

The injection and processing of LiGA is described in Figure 6 (main text); the N-glycan and blank (unmodified) phage data is mirrored from Figure 6b for consistency and replotted as bar chart describing an average FC from n=3 mice and an overlaid scatter plot describing an FC from each mouse.

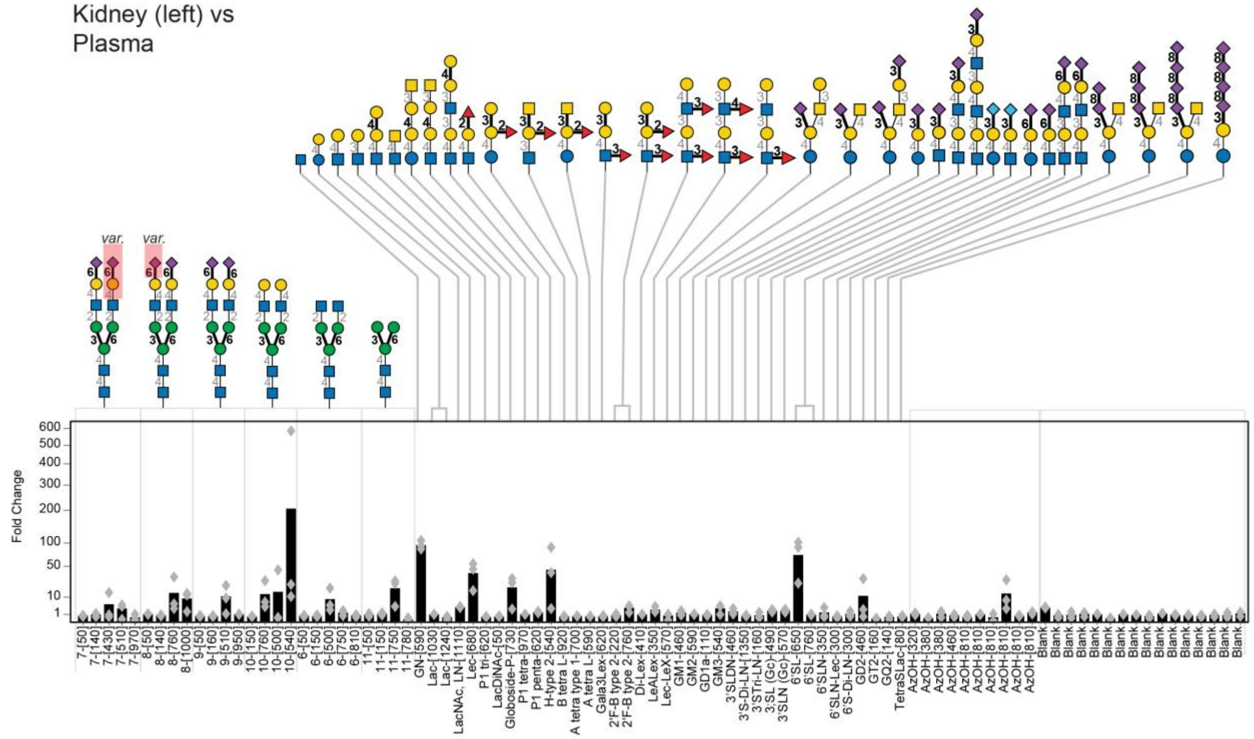

**Figure S27.** Summary of LiGA interaction with left kidney *in-vivo* described as fold change (FC) enrichment in left kidney with respect to plasma from the same animal.

The injection and processing of LiGA is described in Figure 6 (main text); the N-glycan and blank (unmodified) phage data is mirrored from Figure 6b for consistency and replotted as bar chart describing an average FC from n=3 mice and an overlaid scatter plot describing an FC from each mouse.

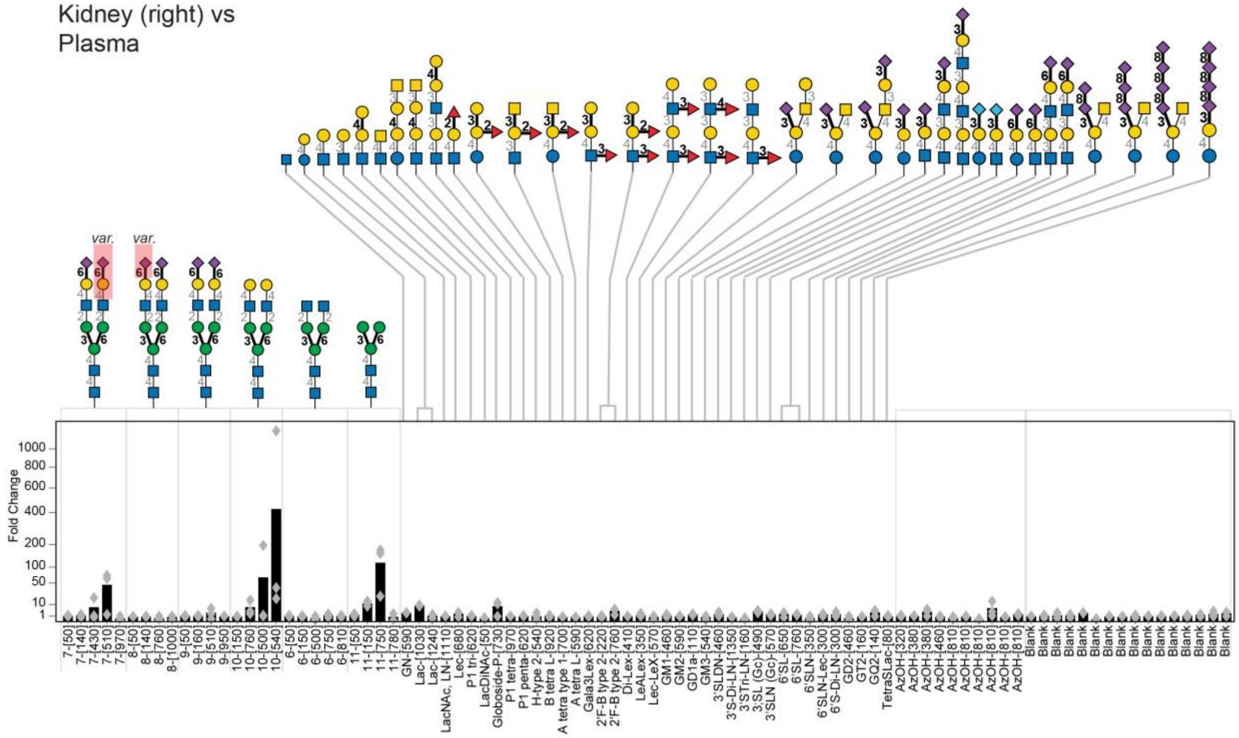

**Figure S28.** Summary of LiGA interaction with right kidney *in-vivo* described as fold change (FC) enrichment in right kidney with respect to plasma from the same animal.
The injection and processing of LiGA is described in Figure 6 (main text); the N-glycan and blank (unmodified) phage data is mirrored from Figure 6b for consistency and replotted as bar chart describing an average FC from n=3 mice and an overlaid scatter plot describing an FC from each mouse.

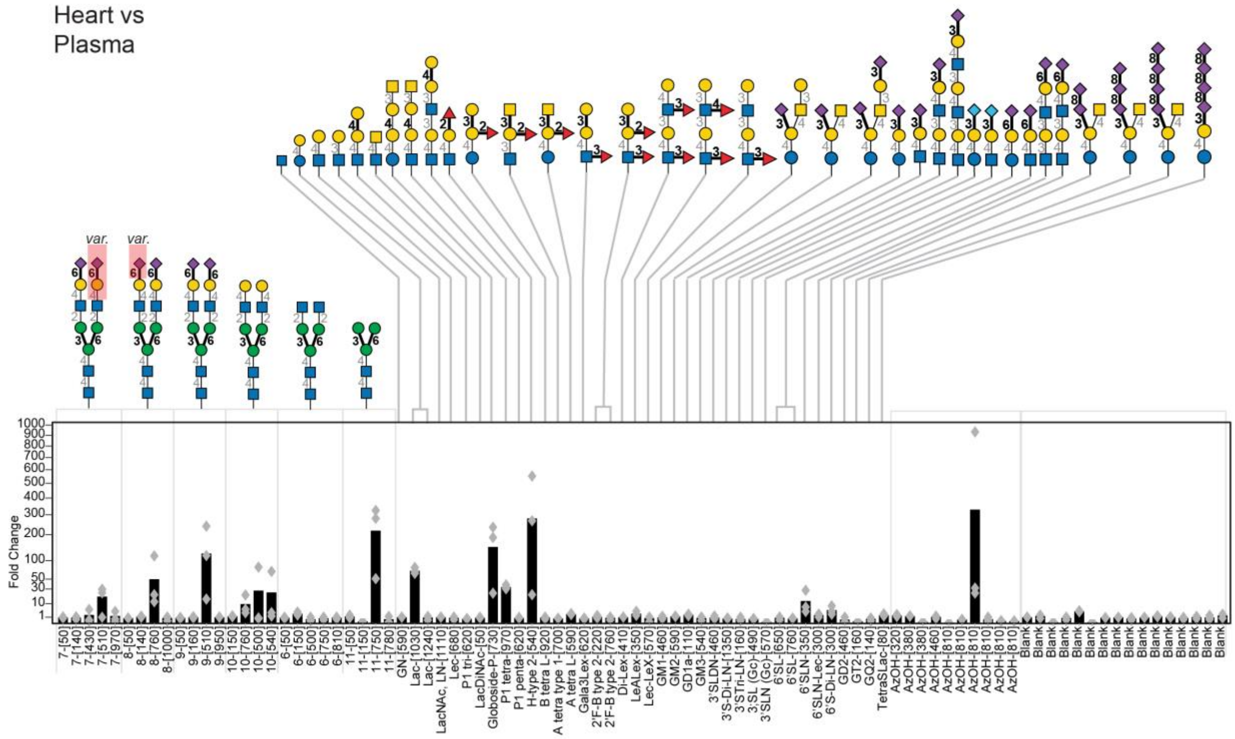

**Figure S29.** Summary of LiGA interaction with heart *in-vivo* described as fold change (FC) enrichment in heart with respect to plasma from the same animal.

The injection and processing of LiGA is described in Figure 6 (main text); the N-glycan and blank (unmodified) phage data is mirrored from Figure 6b for consistency and replotted as bar chart describing an average FC from n=3 mice and an overlaid scatter plot describing an FC from each mouse.

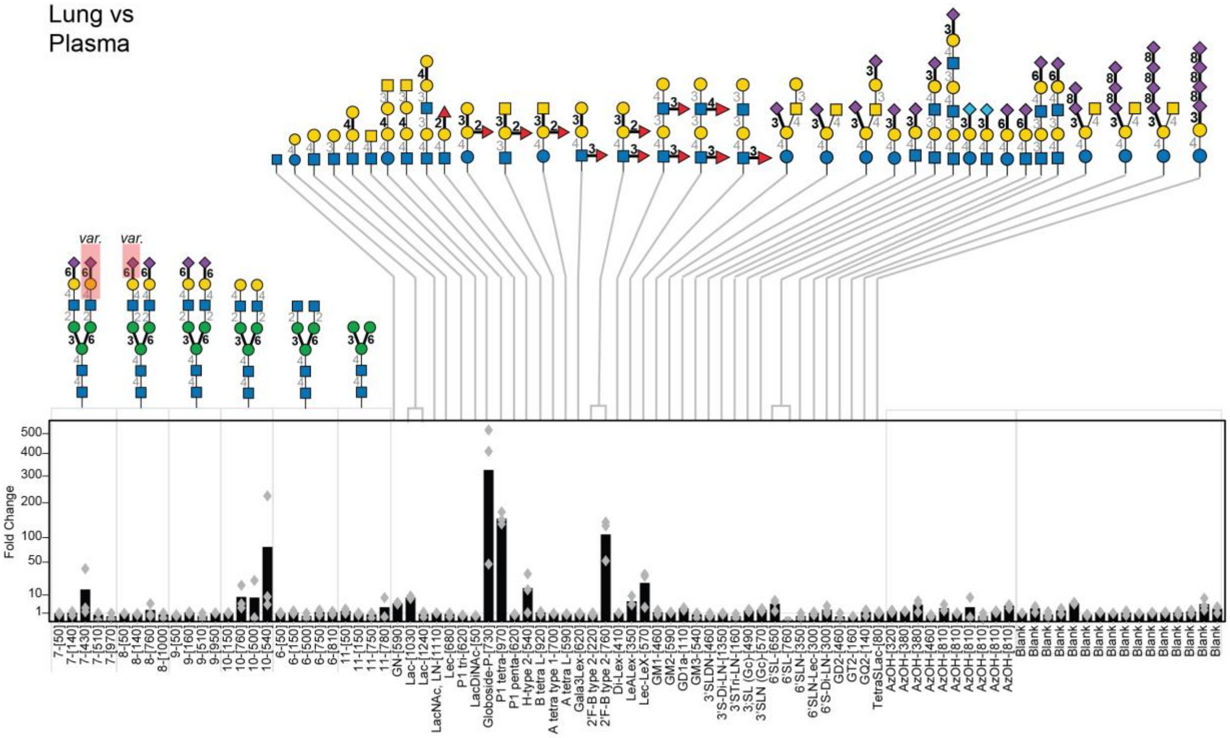

**Figure S30.** Summary of LiGA interaction with lungs *in-vivo* described as fold change (FC) enrichment in lungs with respect to plasma from the same animal.

The injection and processing of LiGA is described in Figure 6 (main text); the N-glycan and blank (unmodified) phage data is mirrored from Figure 6b for consistency and replotted as bar chart describing an average FC from n=3 mice and an overlaid scatter plot describing an FC from each mouse.

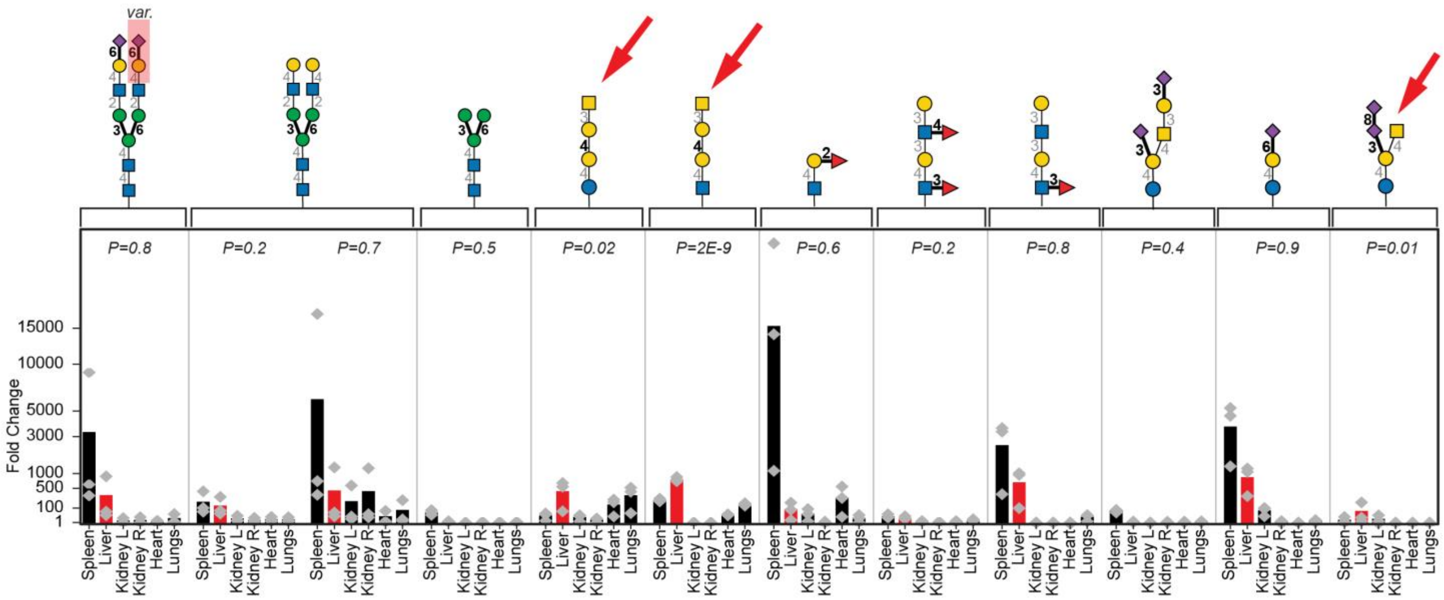

**Figure S31.** Summary of glycans enriched in liver compared to plasma.

Among the glycans enriched in liver compared to plasma, we observed 11 glycans (12 glycan density pair) with FDR<0.05. The FC for terminal GalNAc containing glycans (Globoside-P, P1-tetra and GD2) was highest and statistically significant compared to other organs. Thus highlighting that GalNAc in LiGA can be used for liver targeting, analogous to the many of the GalNAc-siRNA drug candidates in clinical trials.[12] For each of the glycan density-pairs in the plot, whether the fold change is significantly different in liver compared to other tissues was assessed by a one-way ANOVA test (implemented in R version 3.5.2).

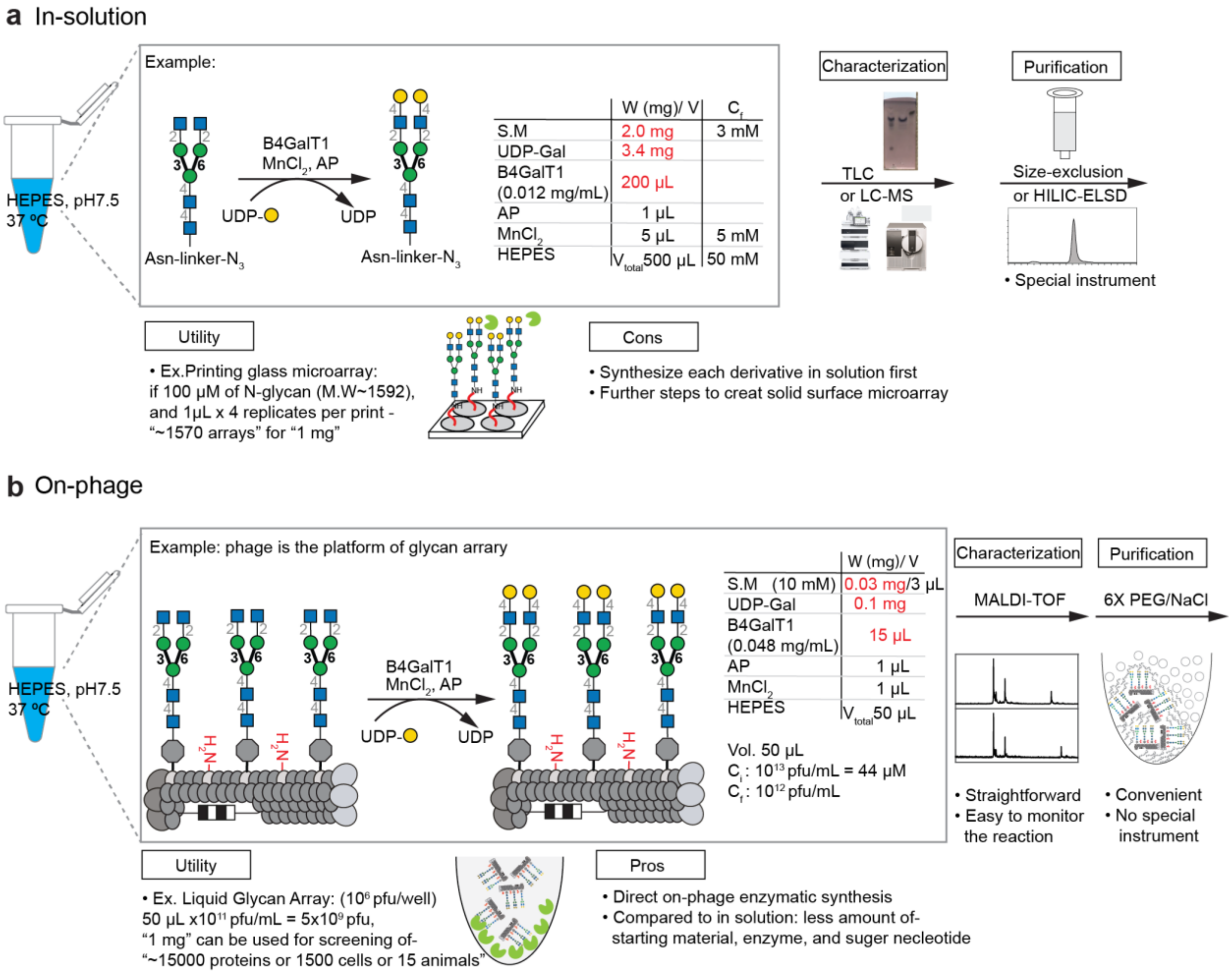

**Figure S32.** Comparison of "on-phage" enzymatic synthesis of liquid glycan array (**a**) with in solution enzymatic synthesis of glycans used to construct glass-based glycan arrays (**b**).

Both **a** and **b** have end point utility section which describes how many arrays can be generated from a fixed amount of each glycan (e.g., 1 mg of glycans can be used to print 1570 glass-based arrays). **b,** Example of on-phage enzymatic synthesis with detailed reaction conditions. The utility describes the amount of glycans needed to generate DNA-coded glycan needed to conduct a certain number of LiGA experiments (e.g., 1 mg of glycan yields enough LiGA for 15,000 protein binding experiments). While **a** and **b** are only an "order of magnitude"-type estimates, they illustrate that LiGA manufacturing requires an order of magnitude less material than the traditional glass-based glycan microarrays.
