## Extended Data Figures for "Chemoenzymatic Synthesis of Genetically-Encoded Multivalent Liquid *N*-glycan Arrays"

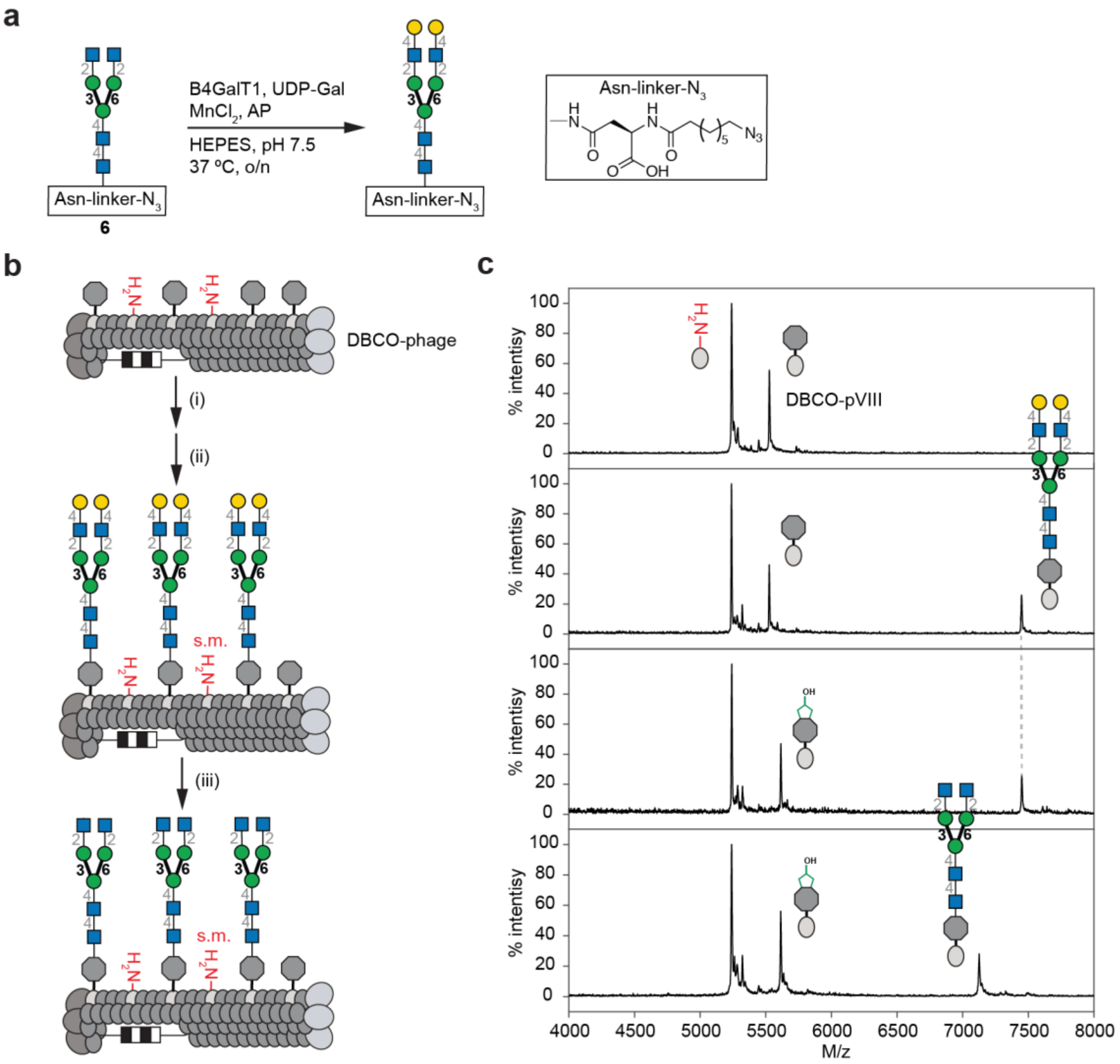


**Extended Data Fig. 1: On phage trimming of "homogeneous" N-linked terminal galactose structure. a,** Compound **10** was used as starting material to prepare "homogeneous" *N*-linked terminal galactose structure. **b,** Scheme of galactosidase treatment of homogeneous *N*-linked galactosylated derivative to afford terminal *N*-acetylglucosamine moieties. **c,** MALDI-TOF MS characterized the azidoethanol capping step after chemical glycosylation to block the unreacted DBCO-pVIII intermediate, and the peak of the desired product was generated after the cleavage of galactose.

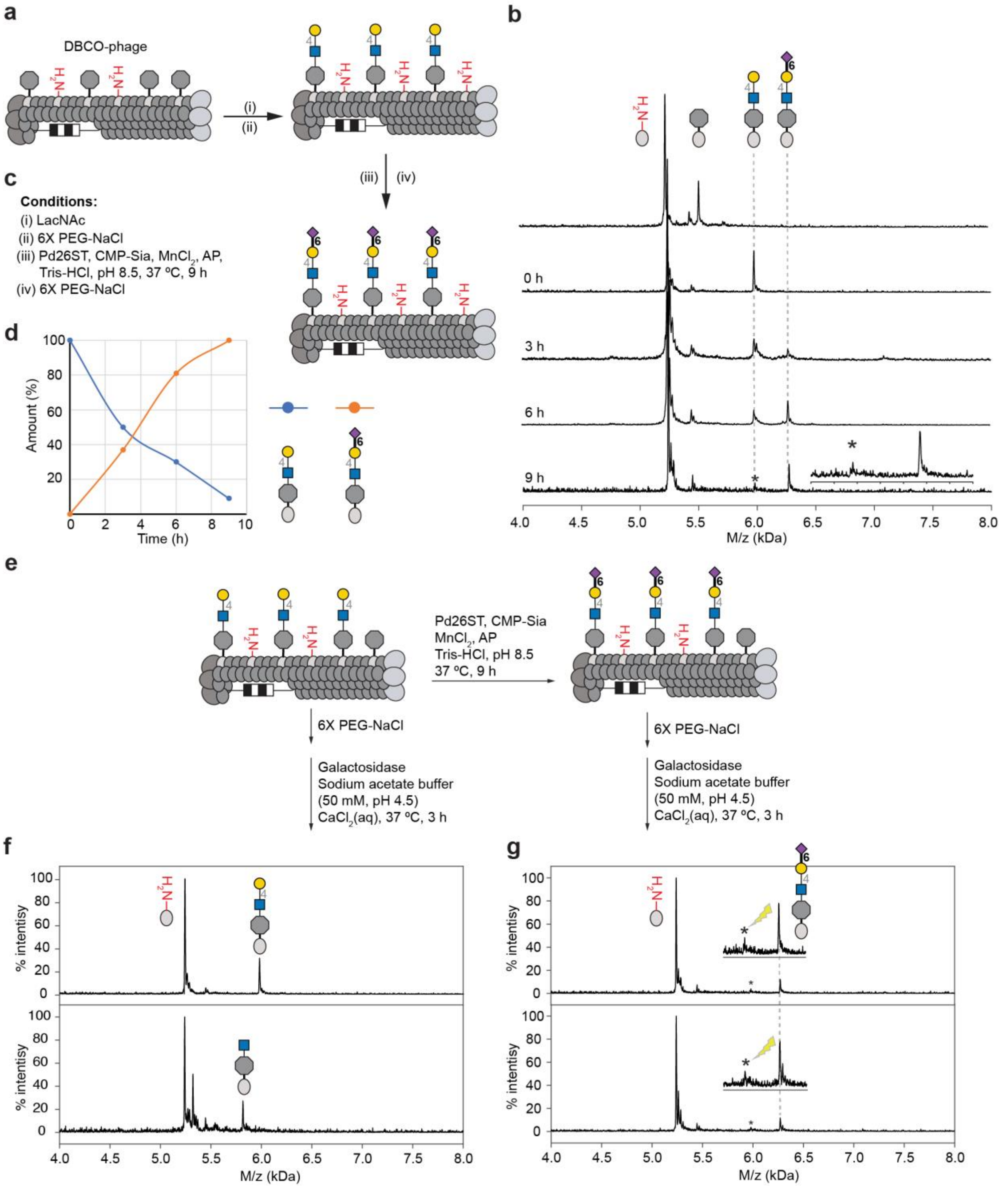


**Extended Data Fig. 2: Model study of Pd26ST-catalyzed sialylation on LacNAc-modified pVIII. a,** Scheme for α-(2 →6)-linked sialylation, with reaction pushed to completion by a 9 h incubation. **b,** Monitoring progress by MALDI-TOF MS indicated detailed time course changes. **c,** Conditions for Pd26ST-catalyzed sialylation. **d,** Plot of time course results with the amount (%) of LacNAc and α-(2 →6)-sialylated LacNAc over time (hours). **e,** Scheme for galactosidase digestion to confirm complete Pd26ST-catalyzed sialylation of LacNAc modified pVIII. (Identical strategy as Extended Data Fig. 5) **f,** MALDI-TOF MS of the positive control. **g,** With the positive control in parallel, MALDI-TOF MS validated the completion of Pd26ST-catalyzed sialylation.

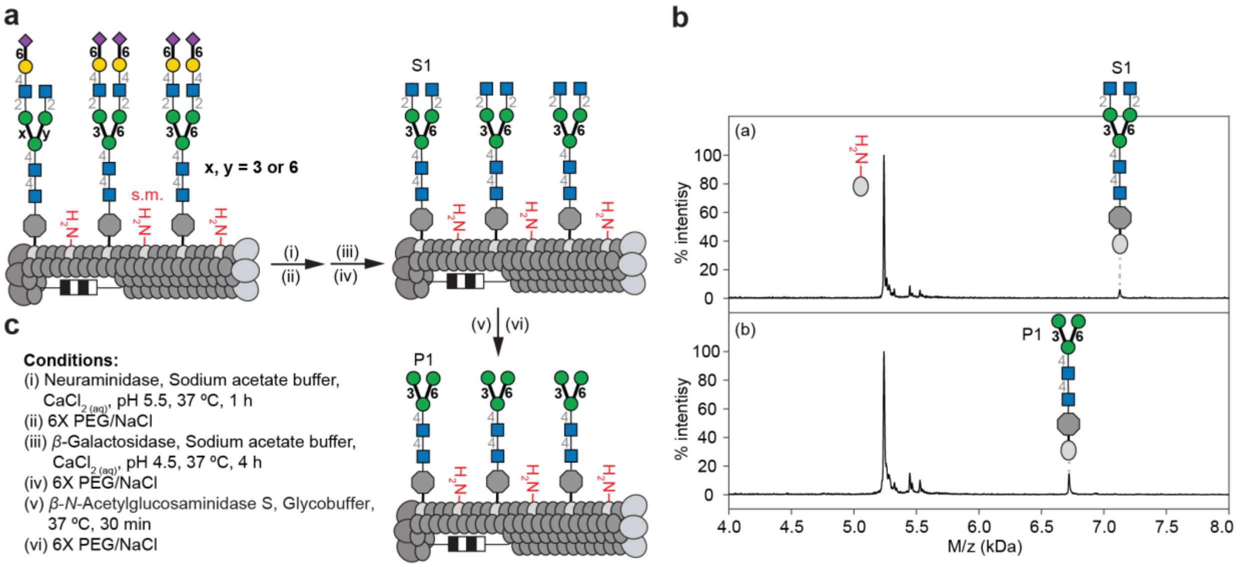


**Extended Data Fig. 3: On-phage enzymatic trimming of biantennary glycosyl-asparagines to generate terminal mannose *N*-glycans ($Man_3GlcNAc_2$). a,** Glycosidase treatment as described in Fig. 4 to afford conjugate P1 with terminal mannose on pVIII. **b,** MALDI-TOF MS characterization: (a) Substrate S1 was obtained as described in Fig. 4. (b) *β*-N-Acetylglucosaminidase S treatment of S1 provided P1 on pVIII. **c,** Conditions of the three glycosidase treatments.

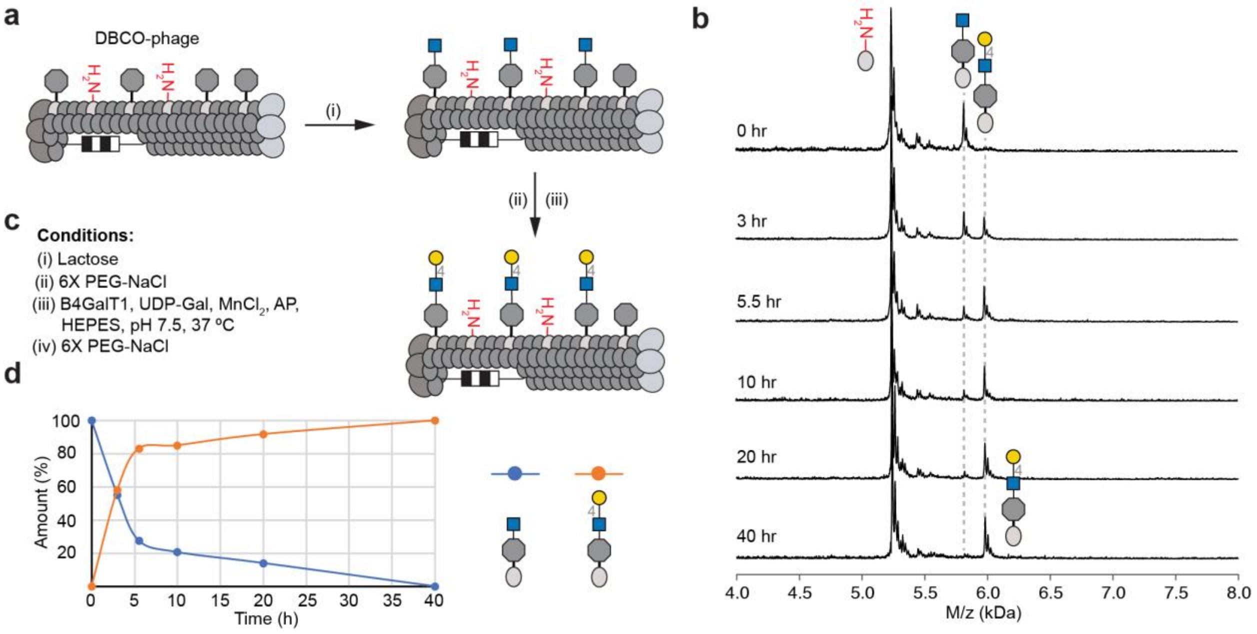


**Extended Data Fig. 4: Model study of B4GalT1-catalyzed galactosylation of GlcNAc to generate LacNAc-modified pVIII. a,** Addition of *N*-acetylglucosamine (GlcNAc) to phage by SPAAC and then $\beta$-(1→4)-galactosyltransferase (B4GalT1)-catalyzed transfer of galactose (Gal) from UDP-Gal. **b,** Progress of the $\beta$-(1→4)-galactosylation as followed by MALDI-TOF MS to indicate time course changes. **c,** Conditions of B4GalT1-catalyzed galactosylation. **d,** Plot of the amount (%) of GlcNAc substrate and LacNAc product over time (h).

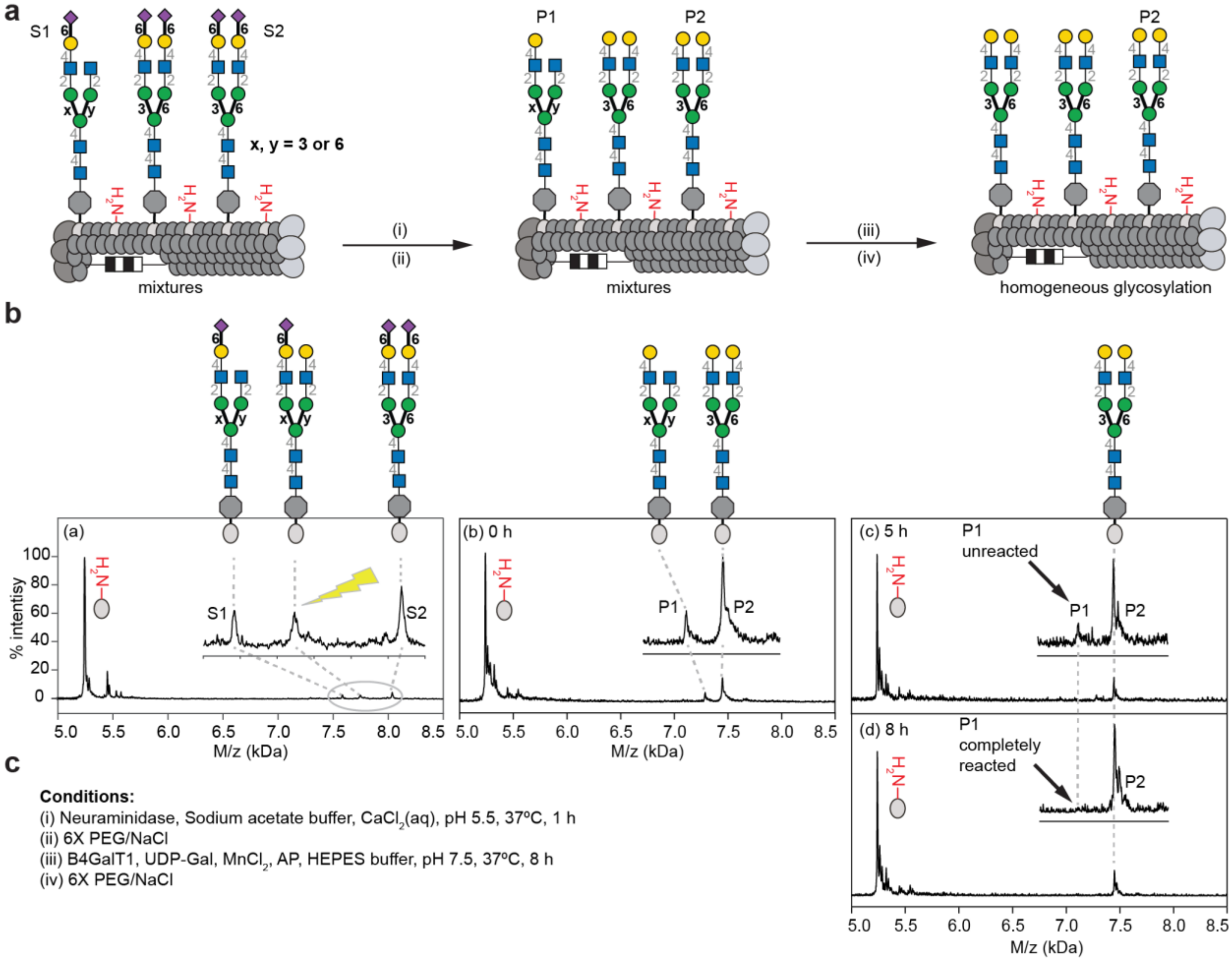


**Extended Data Fig. 5: On-phage B4GalT1-catalyzed galactosylation of biantennary N-glycans. a,** Neuraminidase trimming followed by B4GalT1-catalyzed galactosylation to afford galactose terminated glycosyl-asparagine structure P2 on pVIII. **b,** MALDI-TOF MS characterization: (a) substrates S1, S2 were trimmed by neuraminidase to yield (b) a mixture of products P1 and P2. (c) B4GalT1-catalyzed galactosylation converted P1 to P2 yielding a homogeneous display of P2 on pVIII of M13 phage. **c,** Conditions of each enzymatic treatment step.

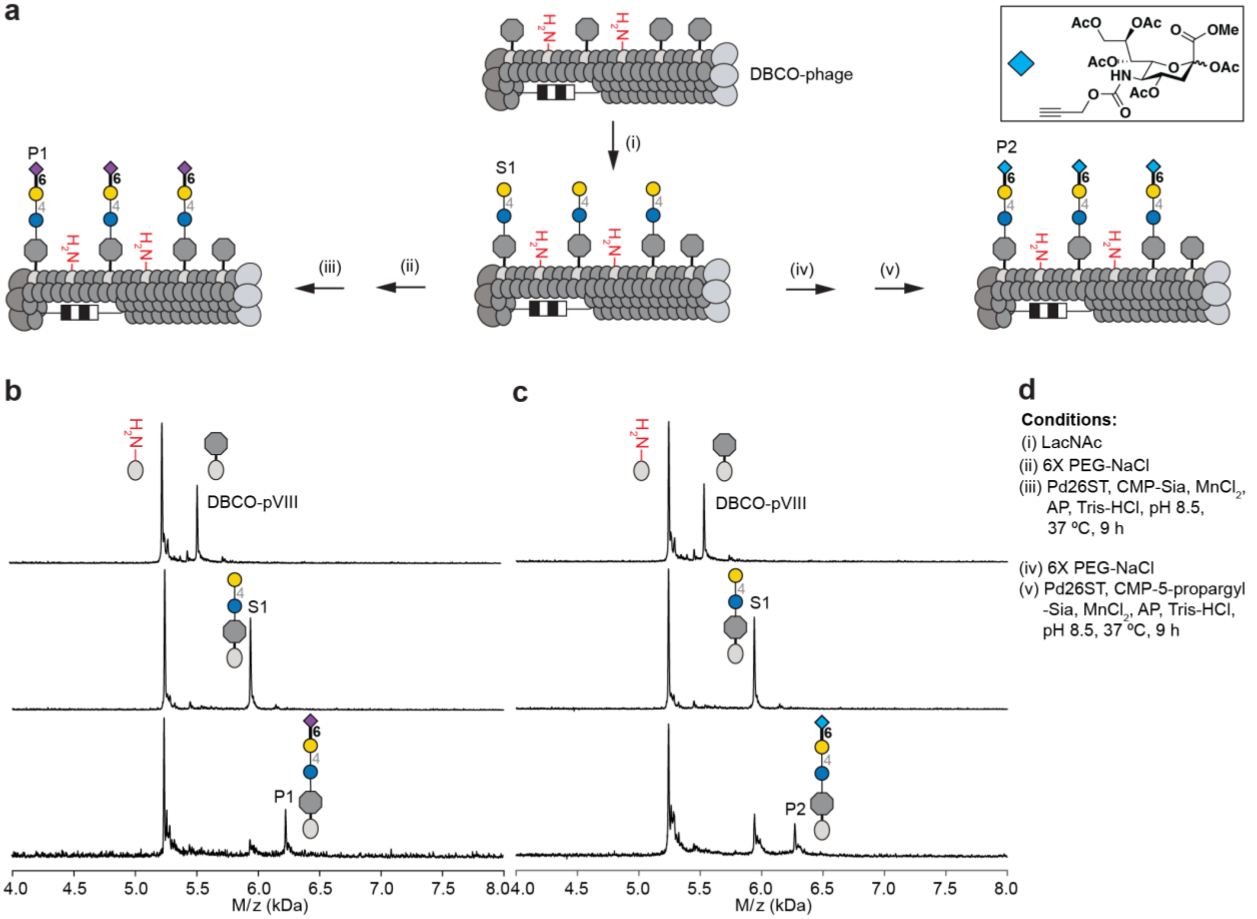


**Extended Data Fig. 6: Model study of Pd26ST-catalyzed sialylation of lactose-modified pVIII and use unnatural CMP-sialic acid as a donor. a,** Scheme for α-(2 →6)-linked sialylation and use of unnatural CMP-sialic acid to modify biantennary N-glycans on pVIII. **b,** MALDI-TOF MS indicates that Pd26ST catalyzes the transfer of N-acetylneuramic acid (Neu5Ac) from CMP-Neu5Ac to lactose (S1) to give α-(2 →6)-sialyl-lactose (P1). **c,** MALDI-TOF MS indicates that Pd26ST S1 transfers a C5-modified-sialic acid residue from C5-propargylamido-CMP-sialic acid to S1 to form product P2 on pVIII. **d,** Conditions for α-(2 →6)-sialylation.

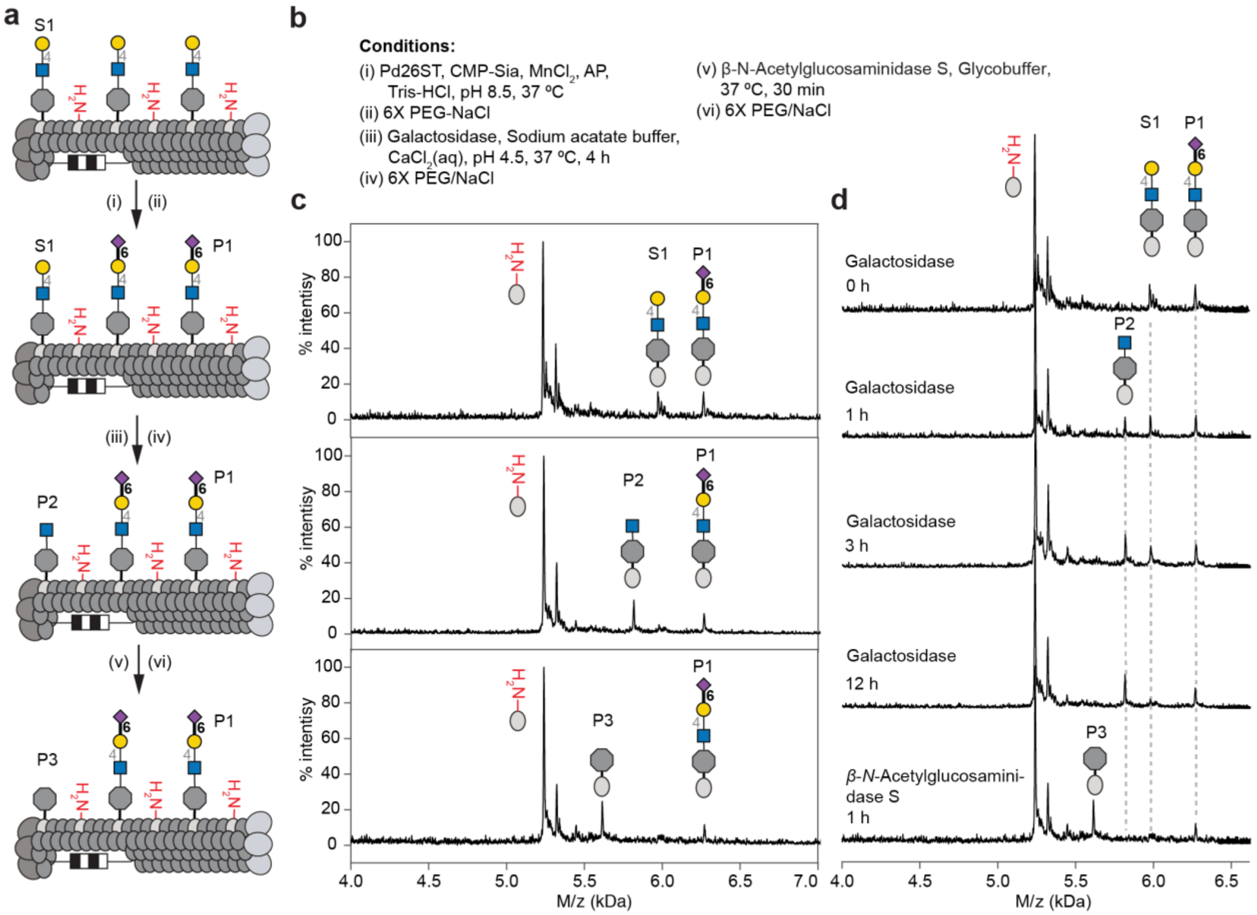


**Extended Data Fig. 7: Glycosidase digestion of incomplete Pd26ST-catalyzed sialylation on LacNAc-modified pVIII. A,** Scheme for α-(2 →6)-linked sialylation, galactosidase treatment and then β-N-acetylglucosaminidase S treatment of LacNAc-modified pVIII. **b,** Conditions for glycosidase digestion. **c,** MALDI-TOF MS of each enzyme treatment step indicate that S1 was converted to P1 by Pd26ST; the reaction was left incomplete on purpose by incubation for only 1 h. Unreacted S1 can be digested to P2 and then P3 by glycosidase treatment, but the sialylated product P1 was not digested. **d,** Progress of glycosidase treatment by MALDI-TOF MS indicated detailed time course changes.

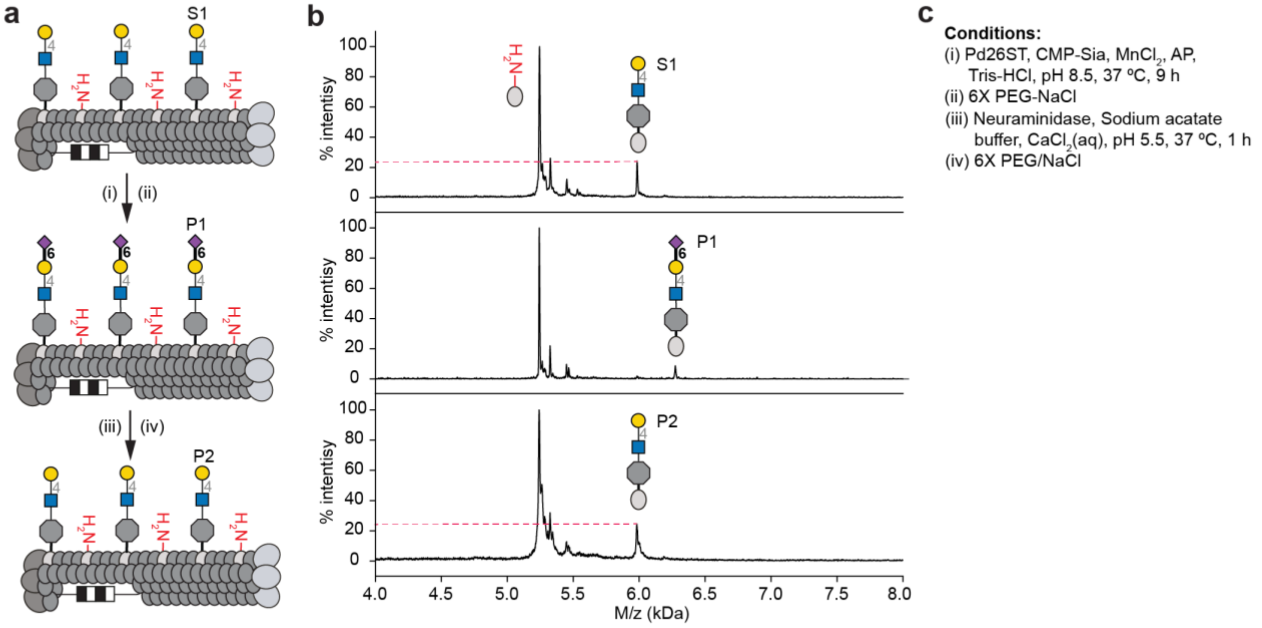


**Extended Data Fig. 8: Comparison of the intensity before enzymatic glycan extension and after enzymatic glycan cleavage on pVIII. a,** Scheme for the intensity comparison study. Neuraminidase was used to cleave sialic acid after Pd26ST-catalyzed sialylation on pVIII. **b,** P1 was generated, followed by neuraminidase cleavage to form P2 (Same structure as S1). e MALDI-TOF MS indicated the corresponding intensity of LacNAc-modified peak S1 and P2 were the same. **c,** Conditions for each step.

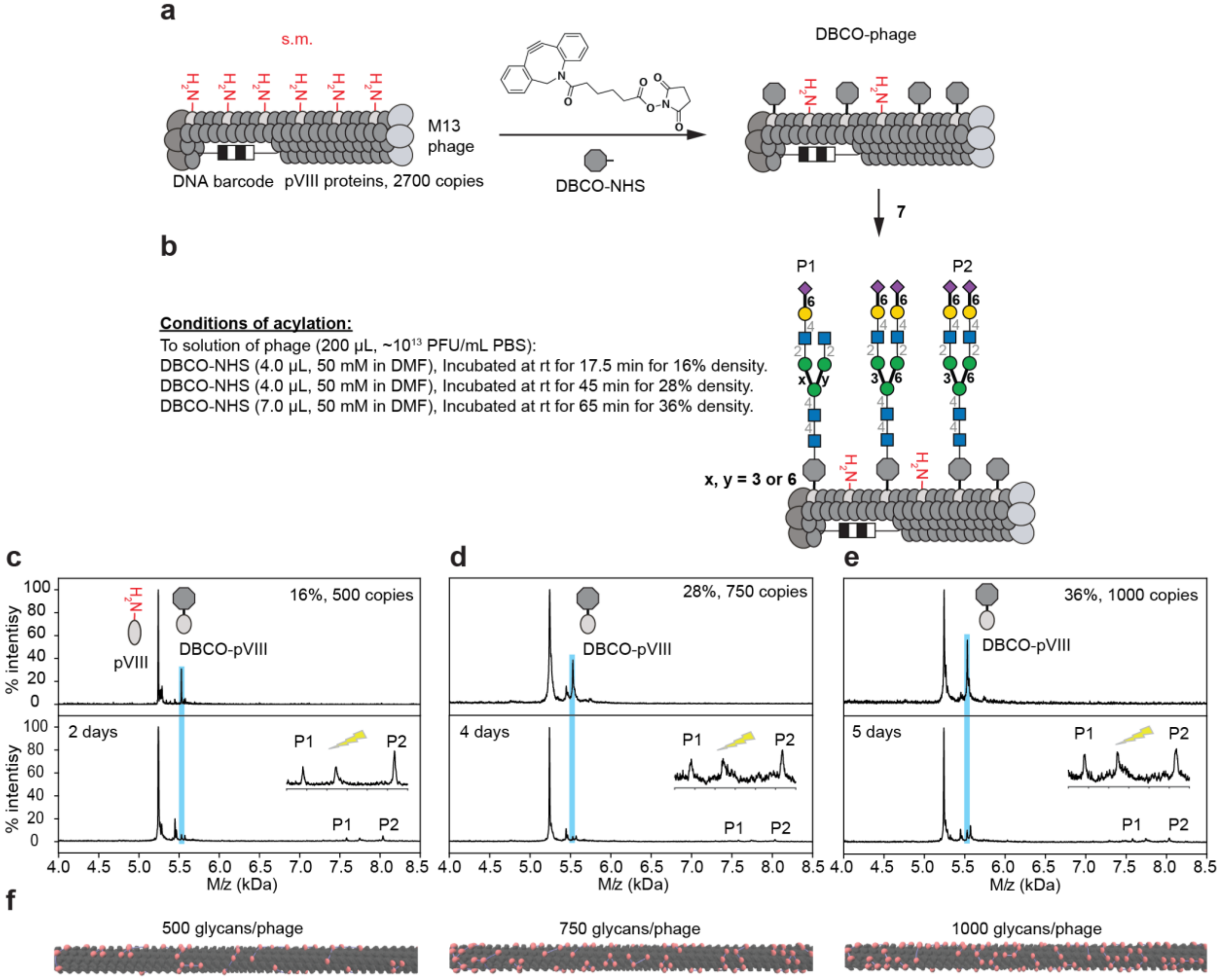


**Extended Data Fig. 9: Example of density control based on the acylation of phage particles to create multivalent display of glycans on M13 phage.** PlotMALDI.m MatLab script were used to process raw MALDI *.txt files to analyze images and integration data. **a,** Example of DBCO-NHS modification followed by glycan conjugation. **b,** To control the number of acylated sites, detailed conditions were shown in **b**. The reaction was desalted using a Zeba filter and analyzed by MALDI-TOF MS. **c,** Out of 2700 copies of pVIII proteins, 16% (~500 copies) were acylated. **d,** Out of 2700 copies of pVIII proteins, 28% (~750 copies) were acylated. **e,** Out of 2700 copies of pVIII proteins, 36% (~1000 copies) were acylated.
